# 1,2,4-Triazole-based first-in-class non-nucleoside inhibitors of bacterial enzyme MraY

**DOI:** 10.1101/2025.01.30.635793

**Authors:** Tomayo Berida, Tzu-Yu Huang, Stefanie C. Weck, Marcel Lutz, Samuel R. McKee, Nathalie Kagerah, Destinee Manning, Mohamed E. Jahan, Sushil K. Mishra, Robert J. Doerksen, Christina L. Stallings, Christian Ducho, Sudeshna Roy

**Affiliations:** Department of BioMolecular Sciences, University of Mississippi, University, MS, 38677, USA; Department of Molecular Microbiology, Center for Women’s Infectious Disease Research, Washington University School of Medicine, St. Louis, MO, 63110, USA; Department of Pharmacy, Pharmaceutical and Medicinal Chemistry, Saarland University, Saarbrücken, 66123, Germany; Department of Biomedical Engineering, University of Mississippi, University, MS, 38677, USA; Department of Pharmaceutical Sciences, University of Tennessee Health Science Center, Memphis, TN, 38103, USA

**Author notes:** Warren Center for Neuroscience Drug Discovery and Department of Pharmacology, Vanderbilt University, Nashville, Tennessee 37232, USA.

## Abstract

MraY, a bacterial enzyme crucial for the synthesis of peptidoglycans, represents a promising yet underexplored target for the development of effective antibacterial agents. Nature has provided several classes of nucleoside inhibitors of MraY and scientists have modified these structures further to obtain natural product-like inhibitors of MraY. The natural products and their synthetic analogs suffer from non-optimal *in vivo* efficacy, and the synthetic complexity of the structures renders the synthesis and structure-activity relationship (SAR) studies of these molecules particularly challenging. In this study, we present our findings on the discovery of first-in-class 1,2,4-triazole-based MraY inhibitors that are not nucleoside-derived. A series of 1,2,4-triazole analogous were identified by a structure–activity-relationship (SAR) study using a structure-based drug design strategy. Compound **1**, with an IC_50_ of 171 µM against MraY from *Staphylococcus aureus* (MraY*_SA_*), was optimized to compound **12a**, exhibiting an IC_50_ of 25 µM. Molecular docking studies against MraY*_SA_* provided insights into these compounds’ binding interactions and activity. Furthermore, screening against the ESKAPE bacterial panel was also conducted, through which we discovered compounds demonstrating broad-spectrum antibacterial activity against *E. faecium*, methicillin-resistant *S. aureus* (MRSA), vancomycin-resistant Enterococci (VRE) strains and *Mycobacterium tuberculosis*. The novel, first-in-class non-nucleoside inhibitors of MraY highlighted in this work provide a strong proof-of-concept of how to leverage structural information of the protein to develop future antibacterial agents targeting MraY.

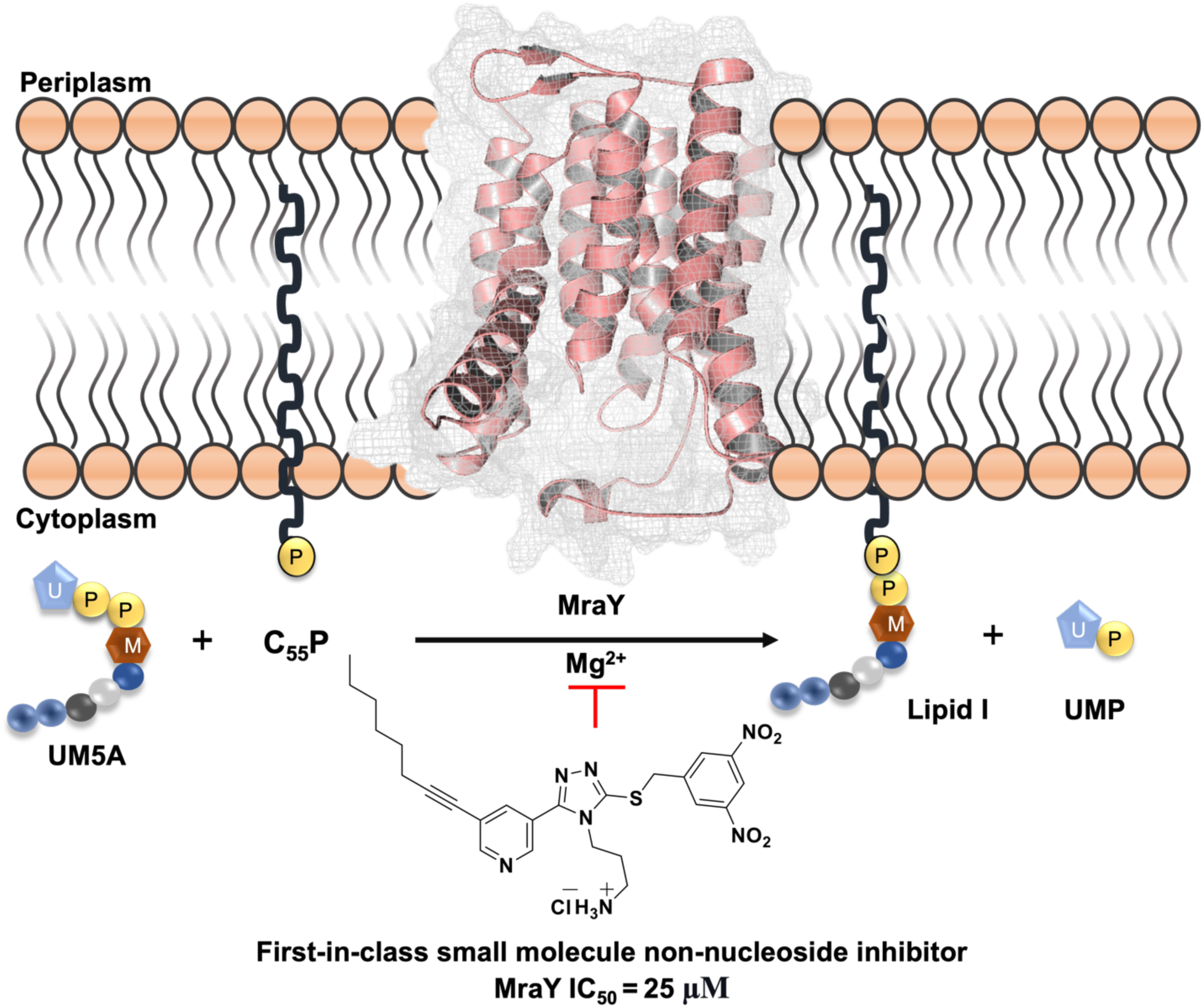

## INTRODUCTION

The growing threats of antibiotic-resistant bacterial infections underscore the critical need for new antibacterial agents with novel mechanisms of action. MraY (phospho-MurNAc-pentapeptide translocase or translocase I), also known as MurX in *Mycobacterium tuberculosis* (*Mtb*), is an integral membrane protein that plays a key role in peptidoglycan biosynthesis and is a promising candidate target for the development of new drugs for the treatment of drug-resistant bacterial infections.^1,2^ The enzyme is responsible for the transfer of phospho-MurNAc-pentapeptide (called Park’s nucleotide) from UDP-MurNAc-pentapeptide (UM5A) to undecaprenyl phosphate (lipid carrier, C_55_-P) to form undecaprenyl-pyrophosphoryl-MurNAc-pentapeptide, a precursor of peptidoglycan also called lipid I (Figure 1).^3–5^

**Figure 1.**
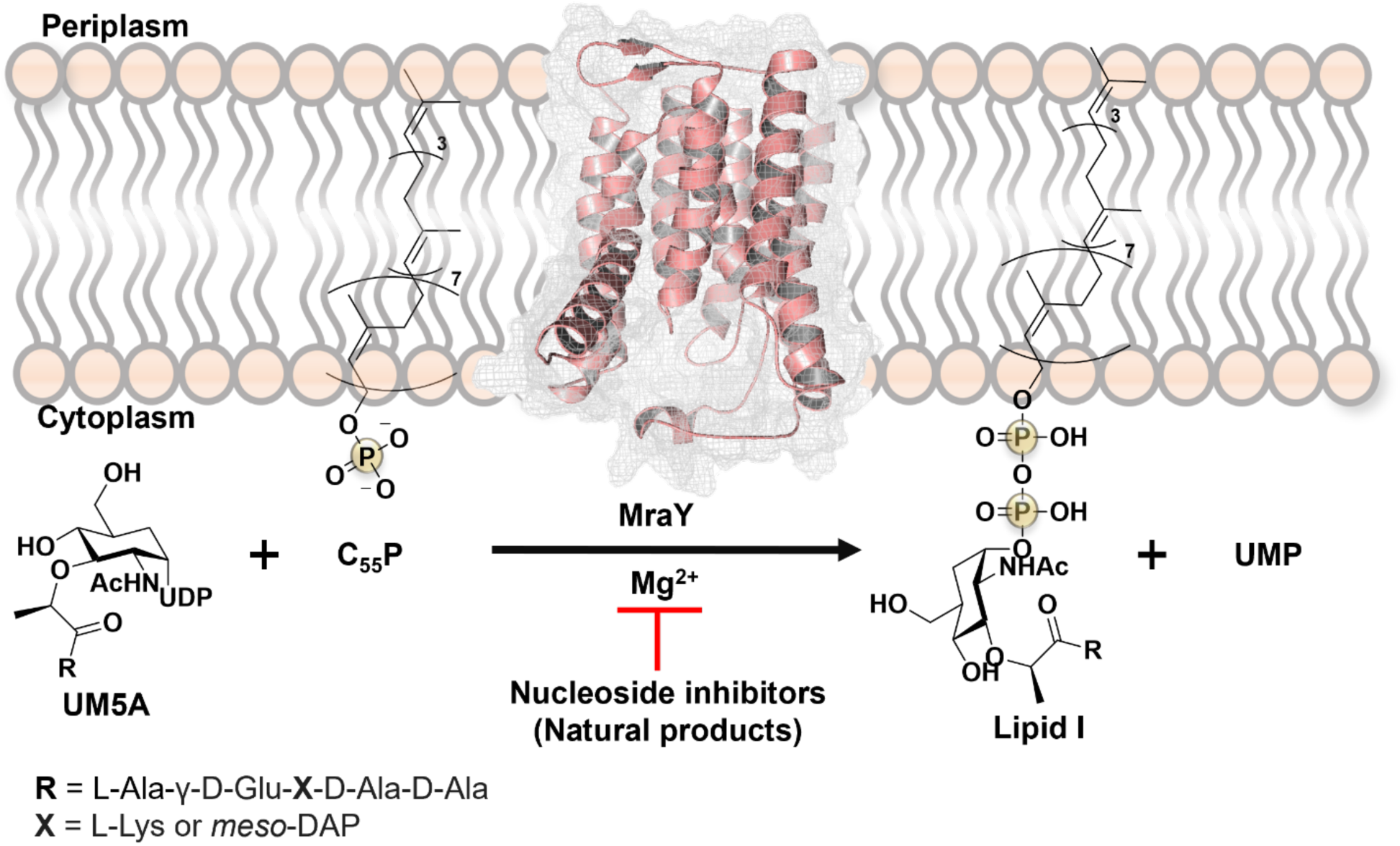
The MraY enzyme facilitates the transfer of phospho-MurNAc-pentapeptide from UDP-MurNAc-pentapeptide (UM5A) to undecaprenyl phosphate (C_55_-P). Using Mg^2+^ as a cofactor, this reaction forms the peptidoglycan precursor, lipid I.

MraY transferase is highly conserved across a wide variety of bacteria, including Gram-negative, Gram-positive, and mycobacterial species.^6,7^ The high conservation of MraY suggests that it plays a key biological function shared among different bacteria and makes it a promising target for broad-spectrum antibiotic agent development. Sequence alignment of the MraY proteins shows 65 amino acid residues are 100% identical, ∼34 amino acid residues are 80% identical, and ∼69 amino acid residues are 60% identical across different bacteria: *Aquifex aeolicus* (MraY*_AA_*, 359 residues), *Escherichia coli* (MraY*_E. coli_*, 360 residues), *Staphylococcus aureus* (MraY*_SA_*, 321 residues), *Mycobacterium tuberculosis* (MraY*_Mtb_*, 359 residues), and *Enterococcus faecium* (MraY*_E. faecium_*, 320 residues) (Figure 2).^8–12^ Among the 27 hotspot amino acid residues in the active site of MraY, 17 amino acid residues are 100% identical, 5 amino acid residues are 80% identical, and 2 amino acid residues are 60% identical across various bacterial species.

**Figure 2.**
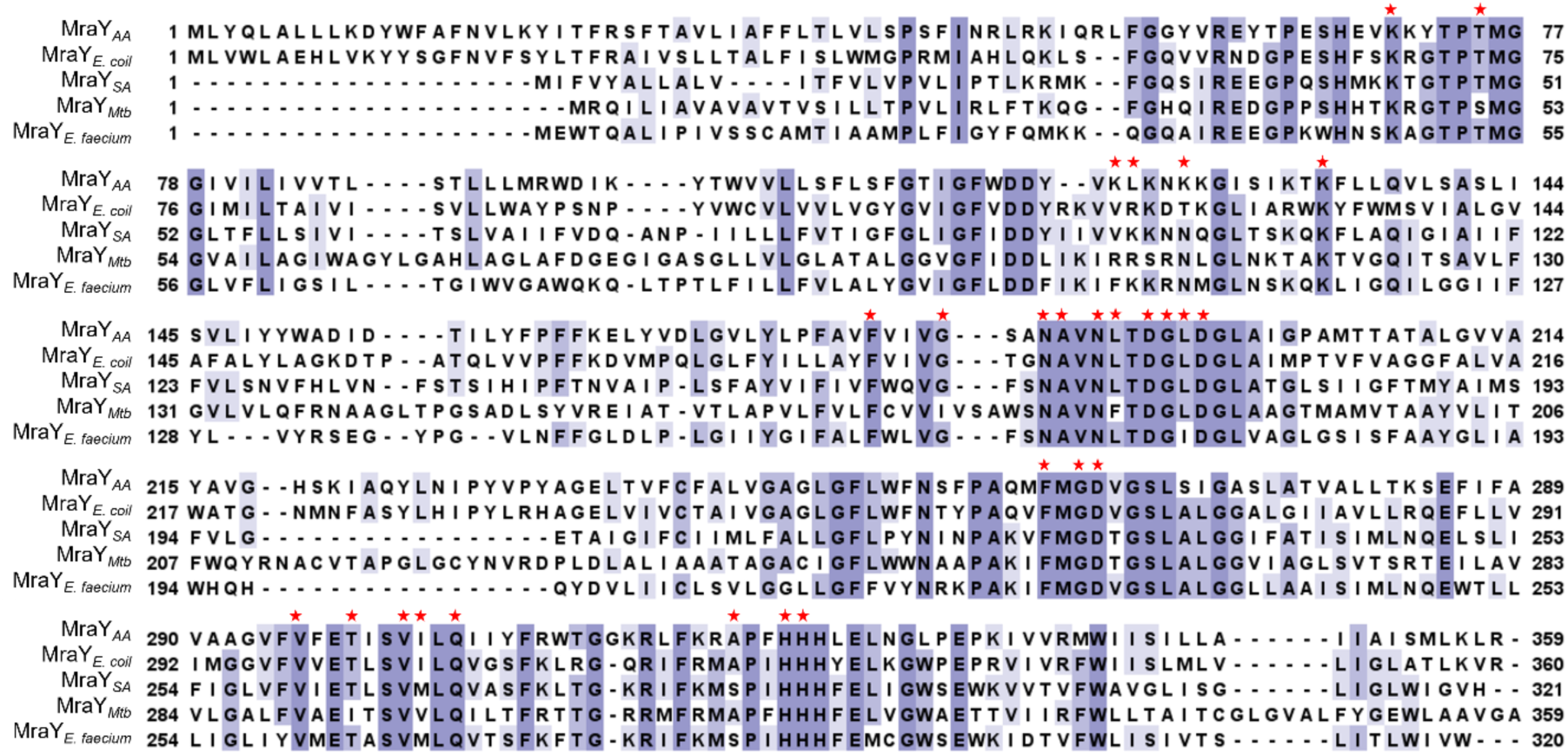
MraY sequence alignments of different bacterial species: *Aquifex aeolicus* strain VF5 (O66465); *Escherichia coli* strain K12 (P0A6W3); *Staphylococcus aureus* strain NCTC 8325/PS 47 (Q2FZ93); *Mycobacterium tuberculosis* strain ATCC 25618/H37Rv (P9WMW7); *Enterococcus faecium* strain D344RRF x C68 (A0A132P3E2), respectively. Fully conserved amino acids across species are in dark purple. The lighter the purple highlight, the less conserved the amino acid residues are. The red asterisks indicate the hotspot amino acid residues in the active site of the MraY-bound inhibitors (nucleoside natural products) with reference to MraY *Aquifex aeolicus* (MraY*_AA_*).^3^ Sequence alignment was performed using UniProt and Clustal W in Jalview.^13,14^

MraY transferase is crucial for bacterial survival, and it is absent in eukaryotes, making it a promising target for antibacterial drug discovery. Six classes of natural product nucleoside inhibitors have been reported that target MraY: liposidomycins/caprazamycins, capuramycins, mureidomycins and related compounds, muraymycins, tunicamycins, and sphaerimicins (Figure 3A).^2,3,15–21^ These nucleoside inhibitors and their synthetic analogs have been the center of extensive research efforts to develop MraY inhibitors as new antibacterial chemotherapeutics.^3,5,15,22^ A breakthrough study by Lee in 2013 reported the first crystal structure of the apo form of MraY*_AA_*, followed in quick succession by reports on co-crystal structures of nucleoside inhibitors bound to MraY from bacterial species *Aquifex aeolicus* (AA) and *Clostridium bolteae* (CB) and its human paralog, GlcNAc-1-P-transferase (GPT), provided much-needed understanding of the structural basis of inhibition of MraY.^3,8,23^ The comparative structural analysis showcases the interactions of each nucleoside inhibitor within the active site of MraY, broken down into different sub-pockets and hotspots of amino acids (Figure 3B & 3C). Overall, there are seven sub-pockets in the MraY active site that are important for its inhibition. These structural insights about MraY catalysis and inhibition provide an avenue to design molecules that can systematically engage in key binding interactions similar to representative classes of nucleoside inhibitors. The deep binding sub-pocket accommodating uracil (Uridine sub-pocket; salmon) is essential for MraY inhibition, engaging in π-π and hydrogen bonding interactions. The uridine moiety is present in both the endogenous MraY substrate UDP-MurNAc pentapeptide and all classes of natural product inhibitors, making them all nucleoside-based. The two other hotspots essential for MraY inhibition are the uridine-adjacent sub-pocket (pale pink) occupied by the 5- aminoribosyl (5ARM) unit found in carbacaprazamycin and muraymycin D2 and the hydrophobic subpocket (cadet gray) occupied by the aliphatic chain, such as those of liposidomycins and tunicamycins. Other hotspots comprise a sub-pocket in the transmembrane 9b (TM9b) and Loop E region (lavender), and the Mg^2+^ coordination site (gold). Two other sub-pockets, the caprolactam (apricot) and tunicamycin sub-pockets (mint), are uniquely occupied by capuramycin and tunicamycin. These pioneering studies elucidated how the nucleoside natural products overcome the challenges of targeting the complex binding site in MraY.

**Figure 3.**
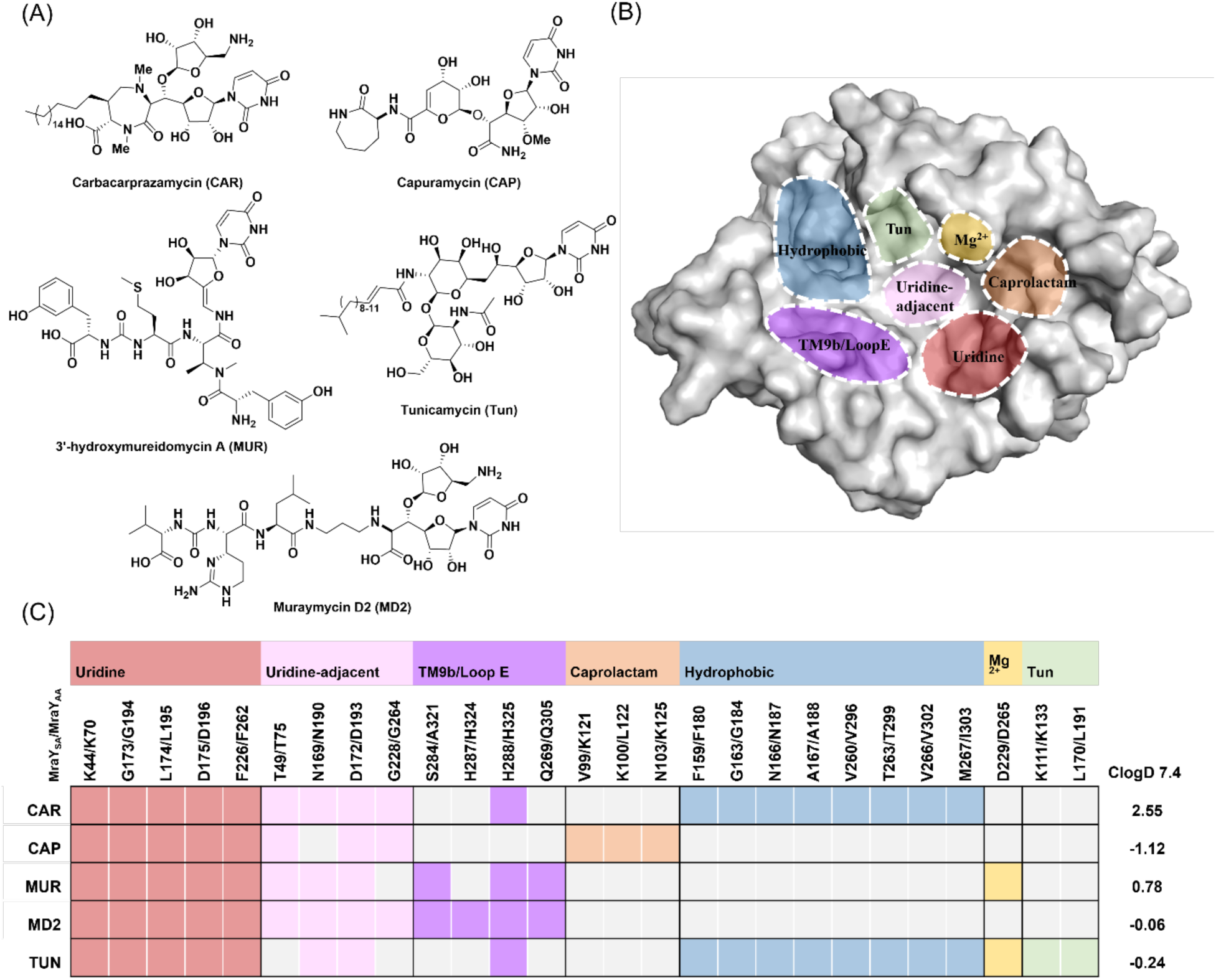
Summary of MraY nucleoside inhibitors, key inhibition hotspots, and their interactions with MraY: **(A)** The structures of MraY nucleoside inhibitors include: carbacaprazamycin (**CAR**), capuramycin (**CAP**), 3′-hydroxymureidomycin A (**MUR**), muraymycin D2 (**MD2**), and tunicamycin (**TUN**). **(B)** The surface representation of MraY*_AA_* highlights inhibitor binding site hotspot subpockets, color-coded as follows: uridine (salmon), uridine-adjacent (pale pink), TM9b/LoopE (lavender), caprolactam (apricot), hydrophobic (cadet gray), Mg²⁺ (gold), and tunicamycin (mint). **(C)** The table summarizes the interactions between MraY*_AA_* and various nucleoside inhibitors, along with their ClogD 7.4 values. Amino acid residue numbers are provided for MraY*_SA_* and MraY*_AA_* based on their sequence alignment. The color-coded squares indicate interactions between specific amino acid residues and the inhibitors, with light gray squares denoting the absence of an interaction. The geometric criteria for interactions are as follows: H bond: the donor and acceptor atoms within 3.3 Å; cation–π: aromatic and charged groups within 4.5 Å; π–π stacking: two aromatic groups stacked face-to-face or face-to-edge; hydrophobic: hydrophobic side chains within 3.6 Å. ClogD 7.4 values of nucleoside inhibitors were calculated using ADMETlab 3.0.^24^ The hotspot analysis figure is inspired by Mashalidis et al.^3^

Nucleoside inhibitors of MraY have been extensively studied for several decades, resulting in several seminal works in this field, which provided the much-needed scientific premise and validated MraY as a viable therapeutic antibacterial target.^7,17,19,25–31^ However, poor pharmaceutical properties of these nucleoside inhibitors barred their clinical development and use, for instance the halted phase I/IIb clinical trials for the semi-synthetic capuramycin analog SQ-641.^32–34^ The major problems associated with the natural product nucleosides include (a) limited *in vivo* efficacy and (b) the complex architecture of these structures making the synthesis and structure-activity relationship (SAR) studies intrinsically challenging.^5,35^ Due to these limitations, there is an unmet need for non-nucleoside small-molecule inhibitors (MW <500 Da) of MraY that can offer better drug-like characteristics with improved pharmacokinetic/pharmacodynamic (PK/PD) properties and potential oral bioavailability. Adjusting the lipophilicity of the small molecule inhibitors is crucial for maintaining both MraY target activity and the corresponding antibacterial activity. This has been a significant issue for nucleoside inhibitors; their MraY target activity often does not correlate with the antibacterial activity. The polar functional groups in the nucleoside inhibitors render a low CLogD value at pH 7.4 (Figure 3C), indicating their high polarity and low lipophilicity. This limits their ability to effectively penetrate the lipophilic bacterial membranes. Chemically diverse small molecules with feasible synthetic access could address this limitation by enabling the design of inherently more lipophilic, non-nucleoside MraY inhibitors. This approach will also allow easier tuning of the structure-activity and structure-property relationships and will be more cost-effective for scale-up. This area has long been and continues to be an unexplored frontier, presenting a significant opportunity for scientific innovation, particularly with the recent advances in structural insights on MraY catalysis and inhibition. The only report of non-nucleoside MraY inhibitors published in this space is from the Bugg laboratory.^31^ Through the screening of the National Cancer Institute (NCI) diversity set, Bugg and co-workers identified phloxine B, a xanthene dye, inhibiting MraY in *E. coli* with an IC_50_ of 32 μM and other bacterial species with an IC_50_ of 128–254 μM. Despite its moderate inhibitory potency, this work is notable as it represents one of the earliest screening efforts specifically designed to identify non-nucleoside inhibitors of MraY, which was identified when very limited structural information of MraY were known.

Based on our previous work and those of others, utilizing a structure-based drug design strategy, this work highlights our proof-of-concept study to create new classes of small molecule inhibitors of MraY that are non-nucleosides.^3,5,8,15,22,23,35–37^ We hypothesized that hybridizing key pharmacophores of the known nucleoside MraY inhibitors and mapping them onto antibacterial agents reported previously in our laboratory (for example, *Mtb* inhibitor structures related to compound **1**, Figure 4) can lead to 1,2,4-triazole-based MraY inhibitors.^38^ Capitalizing on the structural plasticity of MraY to accommodate a wide range of pharmacophores and the reported hotspots and binding interactions in the MraY active site offered a roadmap for designing novel non-nucleoside small molecule inhibitors of MraY.

**Figure 4.**
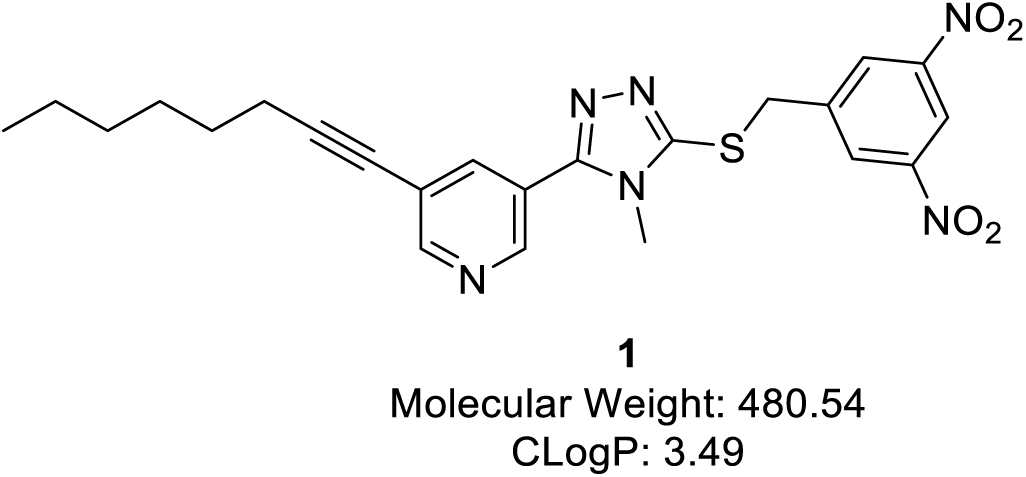
Representative *Mtb* inhibitor **1** from the 1,2,4-triazole class of compounds was used as a proof-of-concept study for the design of non-nucleoside MraY inhibitors.

## RESULTS AND DISCUSSION

Recently, we reported a new series of selective growth inhibitors of *Mtb* with activity in the nanomolar to micromolar range, belonging to the class of 1,2,4-triazoles.^38^ Because of our ongoing efforts to utilize underexploited antibacterial targets, such as MraY, we tested selected potent compounds from the 1,2,4-triazole series for their ability to inhibit MraY*_SA_* in an established fluorescence-based *in vitro* assay.^39^ In brief, overexpressed MraY (usually used as a crude membrane preparation in this assay) was incubated with dansylated Park’s nucleotide, resulting in an increase of fluorescence due to the conversion of the substrate to dansylated lipid I. This increase of fluorescence serves as a measure of MraY activity.^39^ The overexpression of the integral membrane protein MraY is not trivial and needs careful optimization whenever a different homolog is expressed. With respect to the pronounced sequence homology of MraY in relevant bacteria (cf. Figure 2), we, therefore, decided to select one MraY homolog with a robust protocol for overexpression as a representative example and to conduct all *in vitro* MraY assays with this protein, i.e., MraY from *S. aureus* (MraY*_SA_*, overexpressed in *E. coli*). This strategy is also validated by previous studies that had shown different MraY homologs to furnish very similar assay results with a range of uridine-derived natural product MraY inhibitors.^40^ Through this screening of the 1,2,4-triazole series, we identified compound **1** as a weak inhibitor of MraY (IC_50_ = 171 µM).

To further understand its potential mode of binding and molecular interactions compared to known MraY nucleoside inhibitors, compound **1** and tunicamycin (**TUN)** were docked into the active site of MraY*_SA_* (Figure 5) using AutoDock Vina.^41,42^ No crystal structure of MraY*_SA_* has been reported yet. Therefore, an AlphaFold model was retrieved from the AlphaFold Protein Structure Database (AF-Q2FZ93-F1-v4) for the docking study.^43,44^ The AlphaFold model is overall reliable, but the loop residues F29–K43 show a low confidence score (pLDDT < 70) and the conformation of the loop partially covers the active site entrance. Based on the crystal structure of MraY,^1^ in which the coordinates of the amino acids of this loop could not be resolved, we consider this loop to be very flexible and unlikely to block the entrance of molecules to the active site. Since the loop residues do not appear to directly interact with any nucleoside inhibitor,^1,35^ we excluded this loop during docking by removing it from the model. Residue T45 was retained despite its low pLDDT score since it is necessary for maintaining the connection with K44, which appears important for the binding of nucleoside inhibitors. The uracil moiety found in the nucleoside inhibitors and UM5A, the natural substrate of MraY, is known to occupy the uridine sub-pocket formed by residues K44, G173, L174, D175, and F226 (residue numbers based on MraY*_SA_*, which corresponds to the residue numbers K70, G194, L195, D196, and F262 in MraY*_AA_*, PDB ID: 5CKR).^1^ For the rest of this discussion, wherever pertinent, MraY*_SA_* residue numbers will be used.

**Figure 5.**
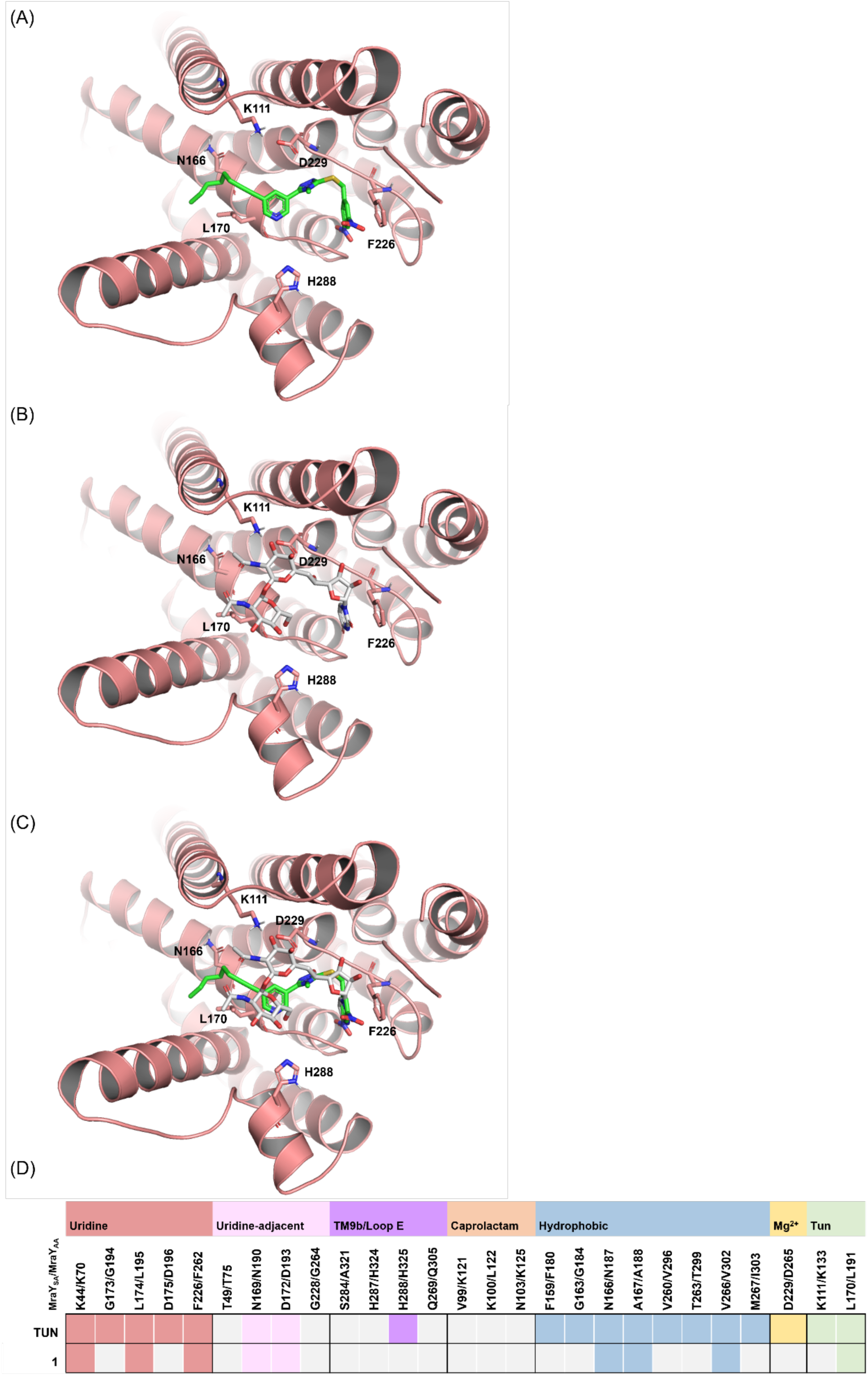
Binding mode of **(A)** compound **1** (green) and **(B) TUN** (gray) in the binding site of MraY*_SA_* as predicted using AutoDock Vina.^41,42^ **(C)** Superimposition of compound **1** and **TUN** in the active site of MraY*_SA_.* **(D)** Table summarizing the interactions between MraY*_SA_* and Compound **1** and **TUN**. Amino acid residue numbers are based on the MraY*_SA_*/MraY*_AA_* sequence alignment. The color-coded squares indicate interactions between specific amino acid residues and the inhibitors, with light gray squares denoting the absence of an interaction based on the docking study. The geometric criteria for interactions are as listed in Figure 3.

The π–π stacking interaction between F226 and the uridine ring in this sub-pocket is believed to be crucial for the binding of all natural product inhibitors to MraY. The geometry of the 3,5- dinitroaryl moiety of compound **1** mimics the pose of the uracil moiety of nucleoside inhibitors in the uridine binding sub-pocket and undergoes the crucial π–π stacking interaction with residue F226 (Figure 5A). Oxygen atoms of the 3,5-dinitro groups interact with D172, L174, and K44 via hydrogen bonds. In the uridine-adjacent sub-pocket, the 1,2,4-triazole core of compound **1** interacted with N169. In the reported co-crystal structures of nucleoside inhibitors with MraY, the ribose moiety of the uridine motif directs the orientation of nucleoside inhibitors toward the hydrophobic binding site. Based on the superimposition of compound **1** and **TUN**, the 1,2,4- triazole core of compound **1** also directs the alkyl chain toward the hydrophobic binding hotspot, which is found to fit into a hydrophobic groove. This groove is thought to be the binding site of lipid substrate carriers, i.e. the C55-P binding site, and the region into which the aliphatic tail of **TUN** and liposidomycins/caprazamycins fit, a class of nucleoside inhibitors that compete with the lipid carriers. This docking study revealed that compound **1** overall adapted a similar binding pose as **TUN** (Figure 5C).

### Structure-activity relationship (SAR) studies of 1,2,4-triazoles as MraY inhibitors

We considered compound **1** as a good starting point to test our hypothesis and conduct an initial round of SAR studies to optimize the potency. Compound **1**, with a molecular weight of 480.54 Da and CLogD 7.4 of 4), showed MraY inhibitory activity of 33% at 100 μM with an IC_50_ 171 µM (Table 1). A wealth of information from the literature guided us through adding essential pharmacophore motifs found in natural product inhibitors of MraY to the central unit of the 1,2,4- triazole-3-thiol series. Design compounds were initially screened against MraY_SA_ at concentrations of 100 μM and 250 μM (Table 1). IC_50_ values were then determined for selected compounds that exhibited substantial MraY inhibition (Table 2). Molecular docking studies of analogs revealed the binding mode of selected compounds and their interactions with key amino acid residues of hotspots in MraY*_SA_*.

**Table 1.**
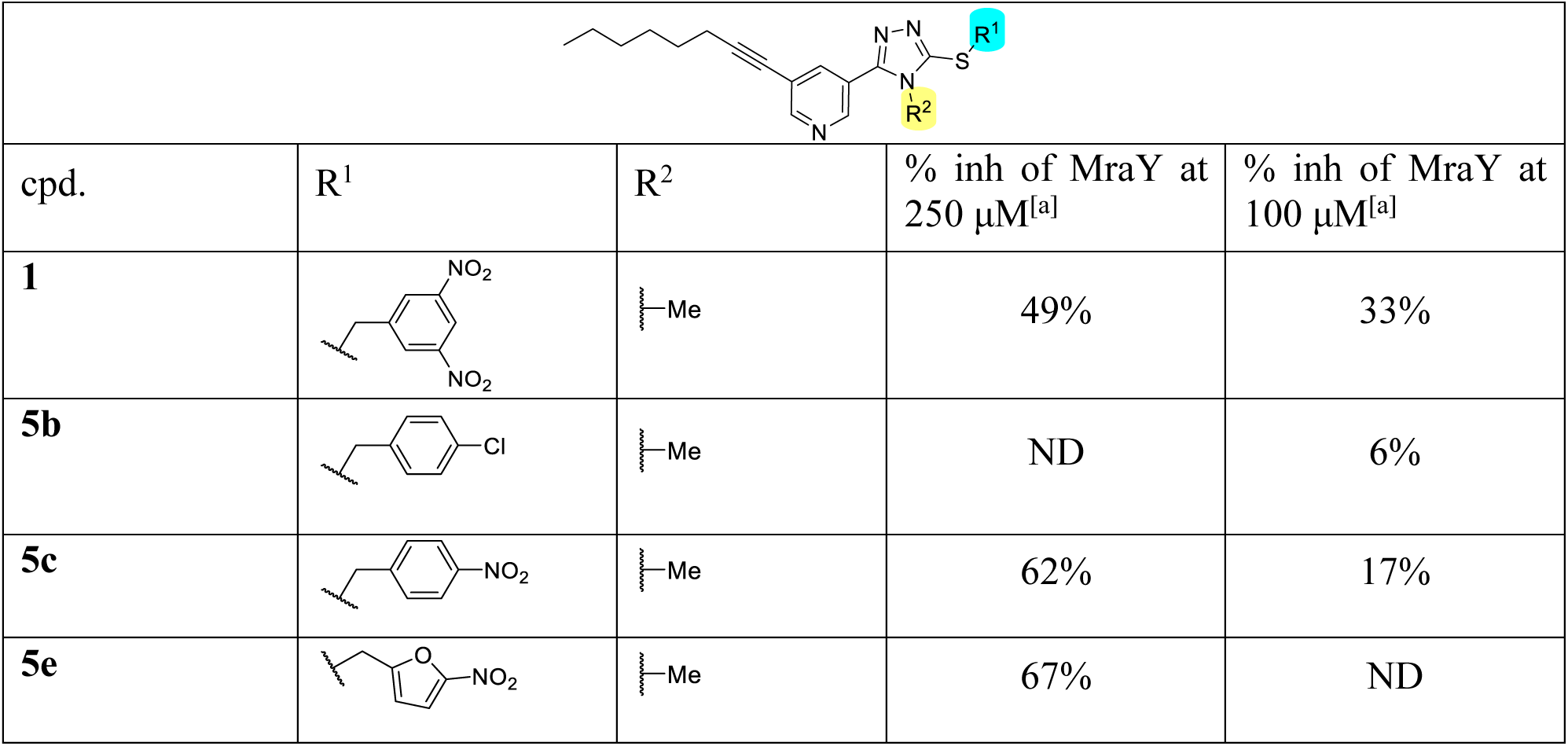
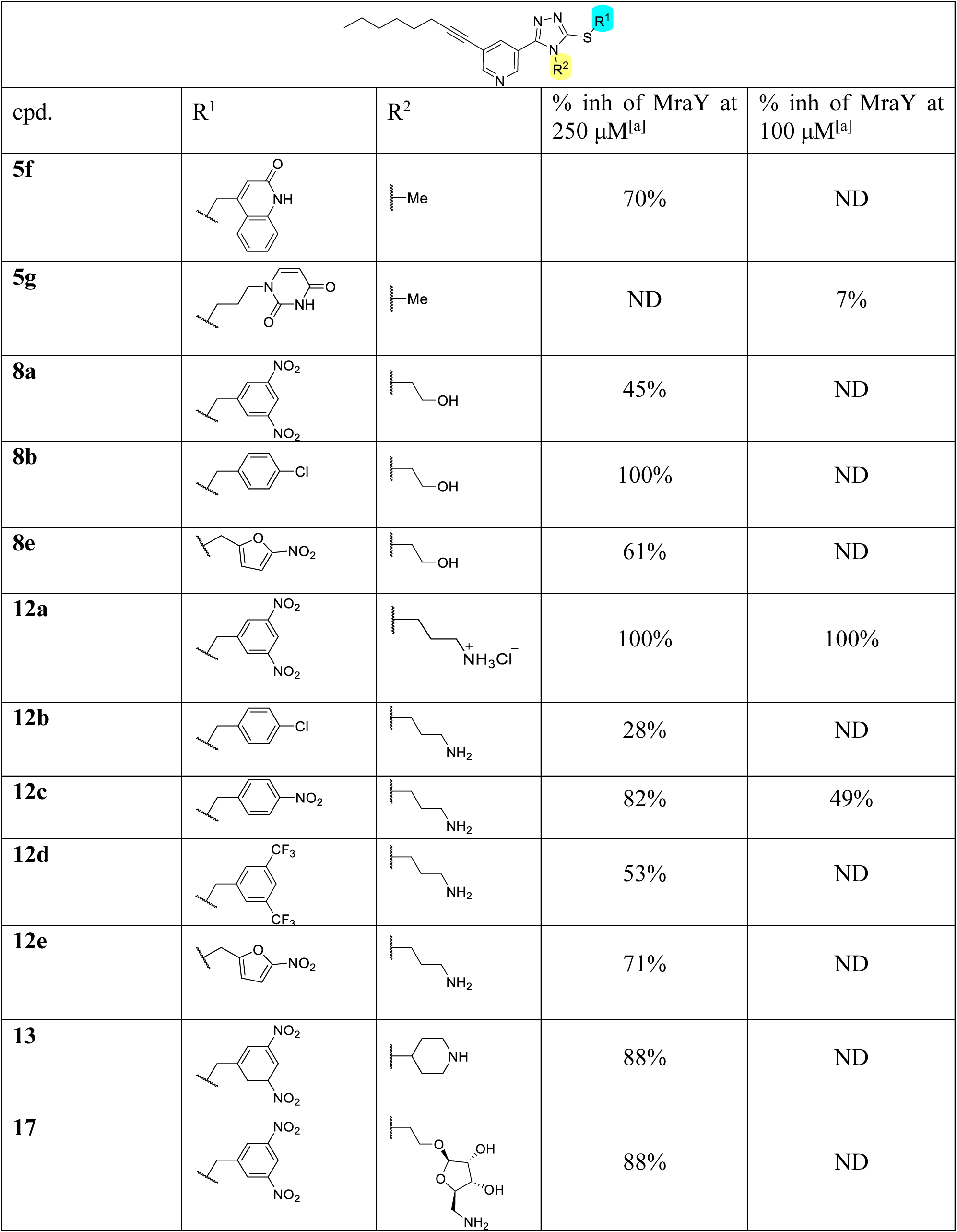
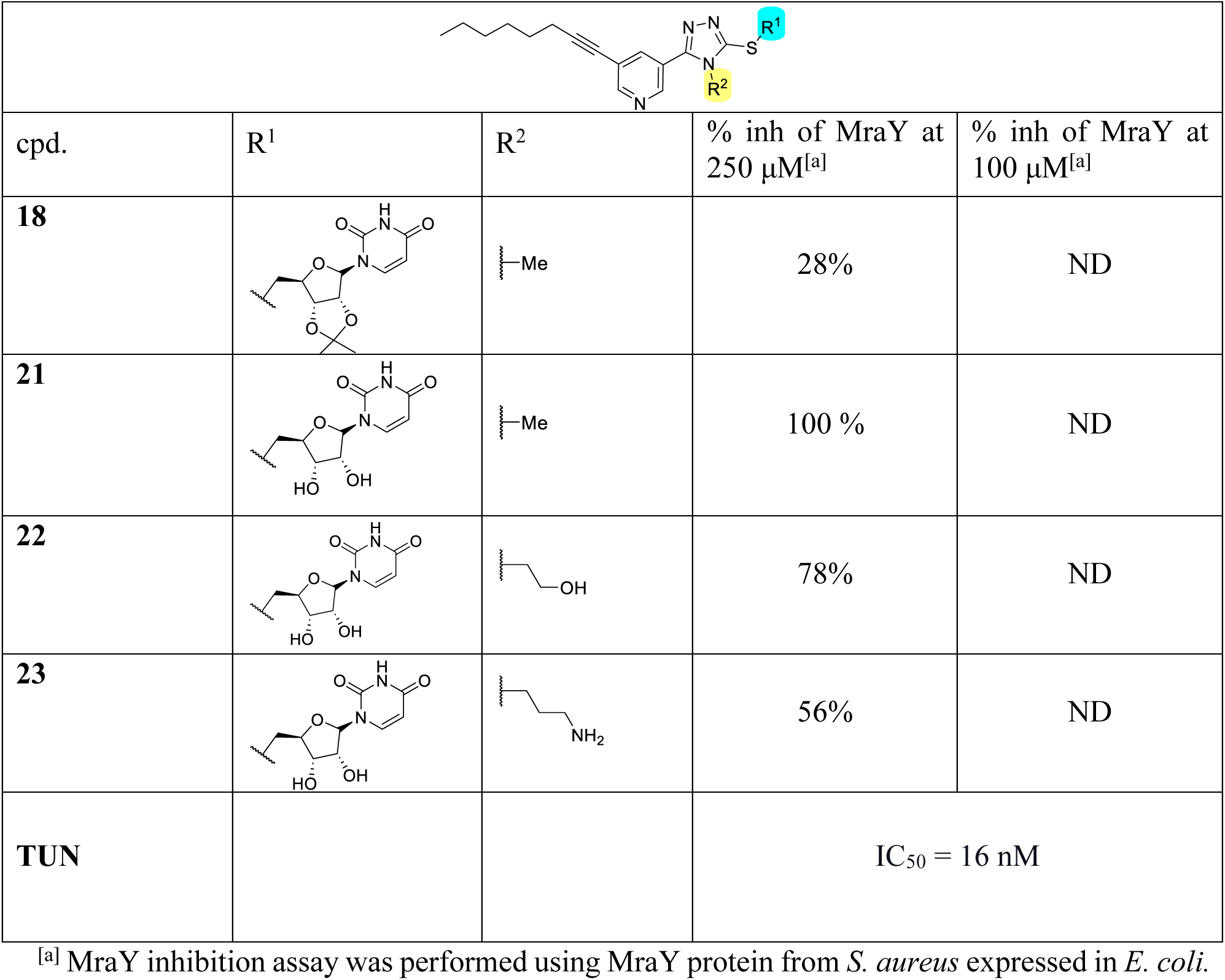
SAR optimization of novel 1,2,4-triazoles inhibiting MraY.

|  |  |  |  |  |
| --- | --- | --- | --- | --- |
| cpd. | R<sup>1</sup> | R<sup>2</sup> | % inh of MraY at 250 μM<sup>[a]</sup> | % inh of MraY at 100 μM<sup>[a]</sup> |
| **1** |  |  | 49% | 33% |
| **5b** |  |  | ND | 6% |
| **5c** |  |  | 62% | 17% |
| **5e** |  |  | 67% | ND |
| cpd. | R<sup>1</sup> | R<sup>2</sup> | % inh of MraY at 250 μM<sup>[a]</sup> | % inh of MraY at 100 μM<sup>[a]</sup> |
| **5f** |  |  | 70% | ND |
| **5g** |  |  | ND | 7% |
| **8a** |  |  | 45% | ND |
| **8b** |  |  | 100% | ND |
| **8e** |  |  | 61% | ND |
| **12a** |  |  | 100% | 100% |
| **12b** |  |  | 28% | ND |
| **12c** |  |  | 82% | 49% |
| **12d** |  |  | 53% | ND |
| **12e** |  |  | 71% | ND |
| **13** |  |  | 88% | ND |
| **17** |  |  | 88% | ND |
| cpd. | R<sup>1</sup> | R<sup>2</sup> | % inh of MraY at 250 μM<sup>[a]</sup> | % inh of MraY at 100 μM<sup>[a]</sup> |
| **18** |  |  | 28% | ND |
| **21** |  |  | 100 % | ND |
| **22** |  |  | 78% | ND |
| **23** |  |  | 56% | ND |
| **TUN** |  |  | IC<sub>50</sub> = 16 nM |  |

<sup>[a]</sup> MraY inhibition assay was performed using MraY protein from *S. aureus* expressed in *E. coli*.

**Table 2.** MraY inhibition activity of selected novel 1,2,4-triazoles analogous.

| Entry | MraY IC<sub>50</sub> (μM) <sup>[a]</sup> |
| --- | --- |
| **1** | 171 ± 36 |
| **12a** | 25 ± 5 |
| **13** | ~100 |
| **17** | 78 ± 5 |
| Tunicamycin | 16 nM |

<sup>[a]</sup> MraY inhibition assay was performed using MraY protein from *S. aureus* expressed in *E. coli* (n = 3).

#### SAR of the eastern region

We first examined replacing the 3,5-dinitrobenzyl ring (occupying the uridine sub-pocket of the MraY binding site) in the eastern region of compound **1** (MraY inhibition of 33% at 100 μM and IC_50_ 171 µM) with several other aromatic groups and non-aromatic groups that would maintain, in theory, the critical π–π interactions in the uridine sub- pocket. Substitution with 4-chlorobenzyl (**5b**) led to a reduction in MraY inhibition (6% at 100 µM). On the contrary, analog **5c** bearing the 4-nitrobenzyl moiety in the eastern region showed similar inhibition in MraY (62% at 250 µM). The nitroimidazole analog **5e** showed comparable inhibitory activity to **5c**. The quinolinone-based O-GlcNAc transferase inhibitors inspired our choice for installing the quinolinone moiety in **5f**.^45^ It is based on a 2-hydroxyquinoline-4- carboxylic acid scaffold that mimics the uridine moiety to bind to the uridine diphosphate (UDP)- binding subpocket of O-GlcNAc transferase (OGT) in GPT enzymes.^46^ This led us to improve MraY percent inhibition (70% at 250 µM) compared to compound **1** (49% at 250 µM). Unfortunately, substituting the 3,5-dinitrobenzyl moiety of **1** with the uracil group connected with a propyl linker led to decreased MraY inhibition (**5g**).

#### SAR of the southern region

The 5-aminoribosyl moiety of **MD2** and the meta-tyrosine moiety of **MUR** are known to fit into the uridine-adjacent sub-pocket of the MraY active site. We hypothesized that introducing a hydrogen bond acceptor or donor that can interact with the T49 or G228 in this sub-pocket would improve MraY activity. To test this hypothesis, we synthesized a series of compounds containing moieties we had explored in the eastern region with an aminopropyl group substituted on the *N*4 of the triazole ring. Compound **12a**, with the 3,5- dinitrobenzyl ring as the eastern region, showed a ∼2-fold improvement in MraY inhibitory activity (100% at 250 µM). 4-Chlorobenzyl (**12b**) (28% at 250 µM) and 4-nitrobenzyl (**12c**) (82% at 250 µM) in the eastern region with methyl substitution in the southern region increased MraY inhibitory activity relative to compounds **5b** and **5c**. The substitution of the eastern region 3,5- di(trifluoromethyl)benzyl (**12d**) (53% at 250 µM) and 5-nitrofuran (**12e**) (71% at 250 µM) showed decreased inhibition compared to **12a**.

Other moieties attempted in the southern region included ethan-1-ol (**8a–8b**, **8e**) and piperidine (**13**). Compound **8a** (45% at 250 µM) did not show a significant improvement relative to compound **1**, but **8e** (61% at 250 µM), **8b** (100% at 250 µM), and **13** (88% at 250 µM) led to slight improvements in MraY inhibitory activity.

Inspired by the 3,4-dihydroxy-3,4-dihyro-2*H*-pyran moiety of **CAR**, we installed a 5- aminoribosyl group in the southern region, which we hypothesized would bind to the uridine- adjacent sub-pocket similar to nucleoside inhibitors.^3^ Compound **17** (88% at 250 µM) displayed a small reduction in MraY inhibition compared to compound **12a**. This might be due to the flexibility caused by the hydrocarbon (n=2) linker attached to the 5-aminoribosyl unit, preventing it from tightly binding to the uridine-adjacent sub-pocket.

#### Nucleoside-based analogs

We next examined whether incorporating the uridine moiety mimicking the nucleoside natural products inhibitors would have an effect on MraY inhibtion. At 250 µM, **18**, **21**, **22,** and **23** showed ∼100%, 78%, 56%, and 28% inhibition, respectively, of the MraY enzyme.

Based on their percent inhibition of MraY, compounds with more than 80% MraY inhibition activity were selected to determine their IC_50_ values against MraY (**Table 2**). Compound **12a** (IC_50_ = 25 µM) demonstrated a roughly 7-fold enhancement in inhibition activity relative to compound **1** (IC_50_ = 171 µM). Other selected compounds, such as compounds **13** (IC_50_ = 100 µM) and **17** (IC_50_ = 78 µM), exhibited similar or slightly worse but still notable inhibitory activities.

### Chemistry

#### Synthesis of eastern and southern regions (***Scheme 1***)

The synthesis of compounds **1** and **5b**–**5d** has been described in our previous publication.^38^ Synthesis of compounds **5**–**17** were carried out using the same protocol with minor modifications where necessary. Briefly, condensation of hydrazide **2** with corresponding isothiocyanates followed by cyclization in the presence of NaOH yielded the respective 1,2,4-triazole-3-thiones (**4**, **7**, **10–11**). For compound **7**, a TBDS protected isothiocyanate **6** was used to prevent side reactions by the hydroxyl group. Fortunately, the TBDS protecting group came off during the cyclization step, eliminating the need for further deprotection. The 1,2,4-triazole-3-thiones (**4**, **7**) were then reacted with the corresponding alkyl or aryl halides to give the desired final products (**1, 5a**–**5g, 8a**–**8b** and **8e**). Analogs **12**–**13** were generated via additional Boc deprotection step of 1,2,4-triazole-3-thiones (**10** and **11**) using 2.5 N HCl in ethanol.

**Scheme 1.**
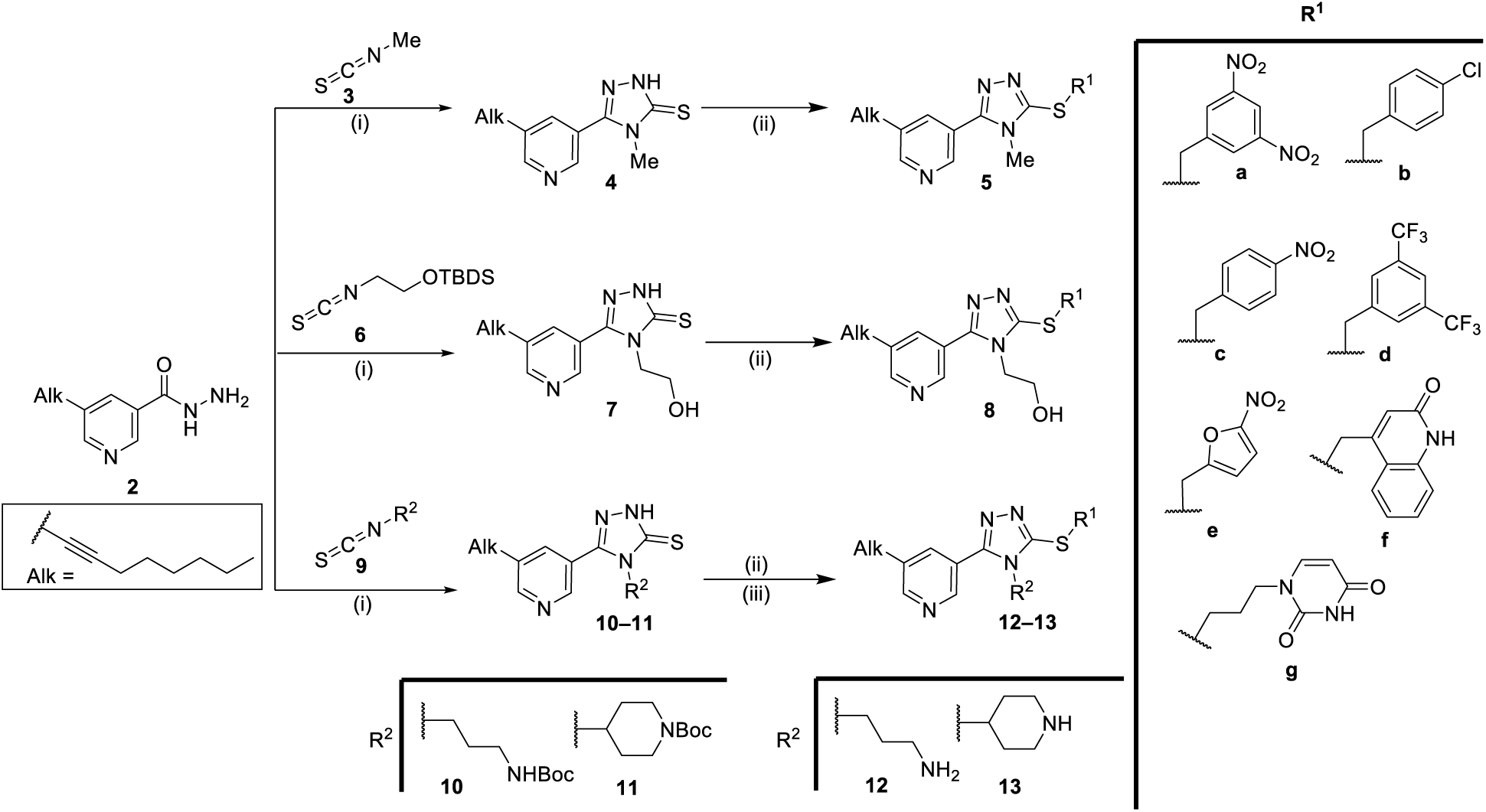
Synthesis of non-nucleoside MraY inhibitors*^[a]^*. *^[a]^*Reagents and conditions: (i) corresponding isothiocyanate, EtOH, reflux, 4 h, then NaOH, 60°C, 3 h, 70–82% (over two steps); (ii) aryl/alkyl halide, NEt_3_, CH_3_CN, 1–24 h, 43–97% or aryl halide, K_2_CO_3_, acetone/MeOH, rt, overnight, 64% for **8**; (iii) 2.5 N HCl/EtOH, rt, 4 h, 30–96%.

### Aminoribosylation of southern region (Scheme 2)

Lewis acid promoted aminoribosylation of **8a** was achieved using glycosyl donor **16a** under BF_3**·**_OEt_2_ conditions to give **14** in 42% yield (**Scheme 2**). Our initial desire to use **16b** for glycosylation was unsuccessful as only trace amount of glycosylation product **15** was detected by HPLC-MS analysis. The azide **14** was converted to a Boc-protected amine (**15**) in a one-pot reaction with a combined yield of 67%. The final compound **17** was obtained following global deprotection using TFA in 80% DCM for 2 h.

**Scheme 2.**
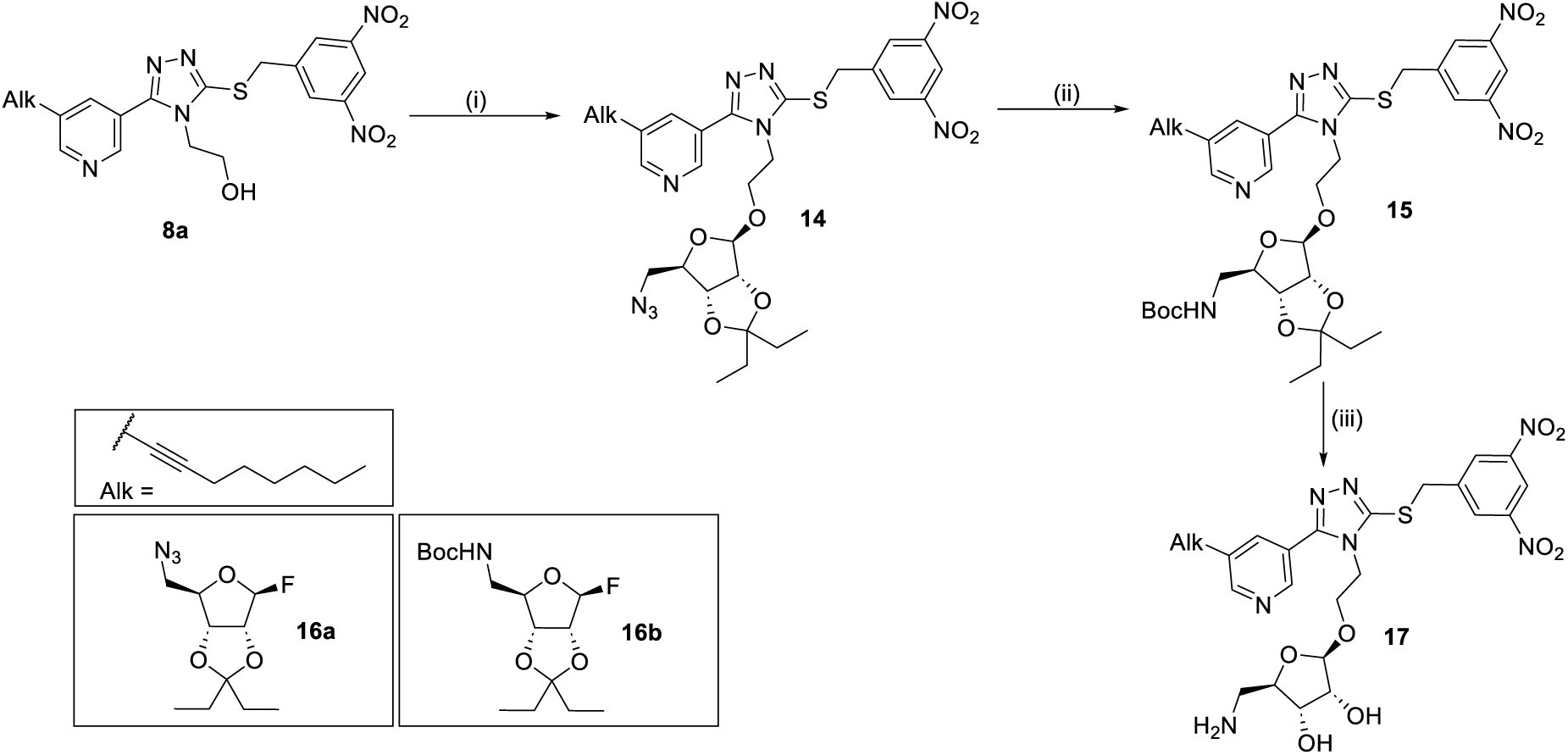
Synthesis of aminoribosylated 1,2,4-triazoles.*^[a]^*. *^[a]^*Reagents and conditions: (i) **16**, BF_3**·**_OEt_2,_ CH_2_Cl_2_, 6 h, 42%; (ii) PPh_3_, H_2_O, THF/toluene, rt 12 h, then Boc_2_O, NaHCO_3_, rt, 2 h, 67% (over two steps); (iii) 80% TFA in CH_2_Cl_2_, rt, 2 h, 40%.

### Synthesis of nucleoside analogs (Scheme 3)

Analogs **21**–**23** were synthesized in two steps from their corresponding 1,2,4-triazole-3-thiones (**4**, **7** and **10**). The final analog **18** was obtained by treating **4** with **24** in the presence of K_2_CO_3_ as a base. Intermediates **19** and **20** were obtained by reacting **24** or **25** with their corresponding 1,2,4-triazole-3-thiones using triethylamine as a base. Intermediates **18**–**20** were then subjected to TFA deprotection to obtain **21**–**23.**

**Scheme 3.**
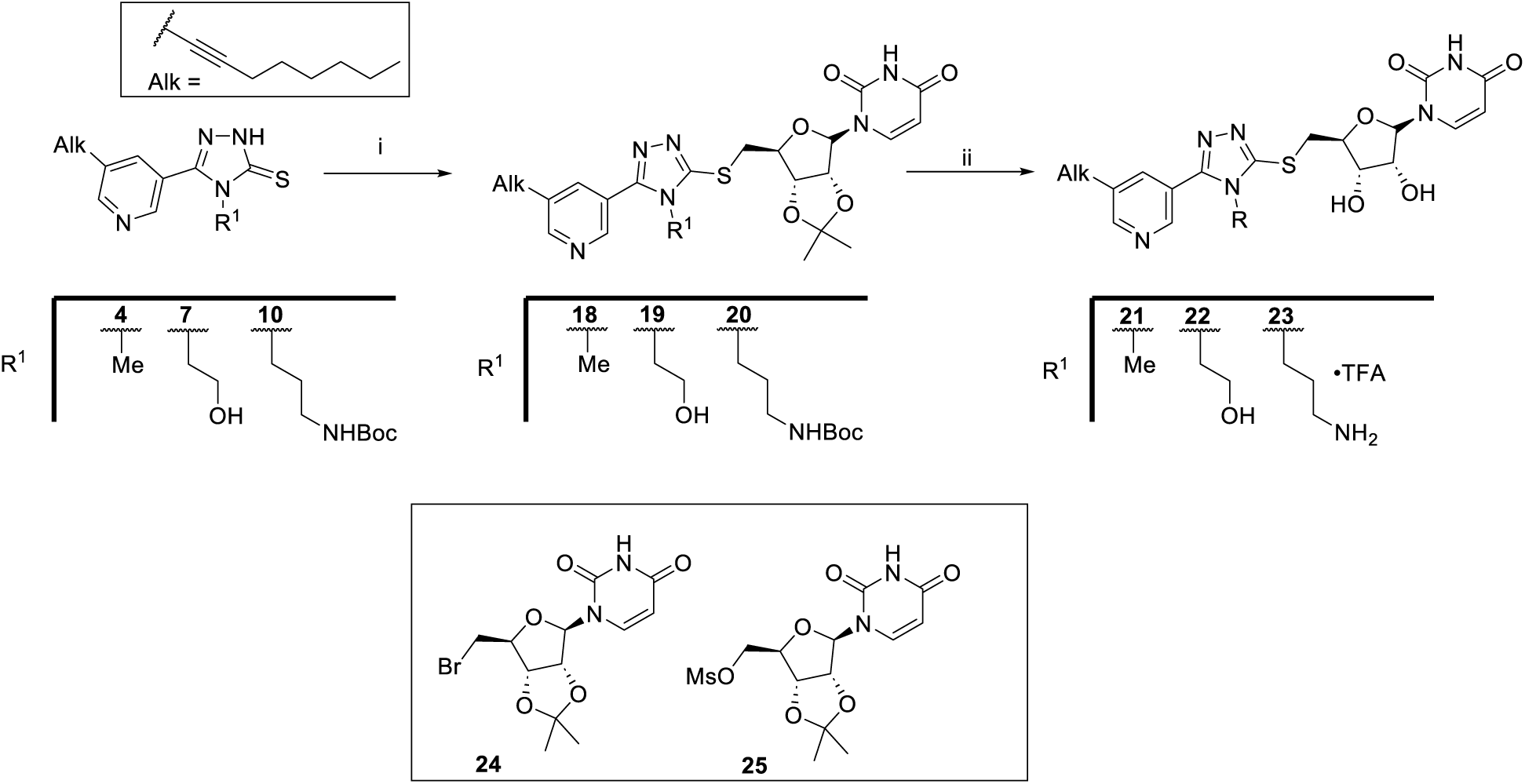
Synthesis of nucleoside analogs.*^[a]^*. *^[a]^*Reagents and conditions: (i) compound **24**, K_2_CO_3_, acetone/MeOH, rt, overnight, 37% for **18**; **24**, NEt_3_, CH_3_CN, overnight, 40% for **19**; and **25**, NEt_3_, CH_3_CN, overnight, 25% for **20.** (ii) 80% TFA in CH_2_Cl_2_, rt, 2 h, 43%–62%.

### Molecular docking of analogs with MraY*_SA_*

Molecular docking studies were conducted on selected compounds to investigate whether the predicted interaction patterns could explain the SAR results for this novel series of small molecule MraY inhibitors. Compounds **12a**, **13**, and **17** (colored green in Figure 6), with more than 80% inhibition activity against MraY at 250 µM inhibitor concentration, were selected and docked into the active site of MraY*_SA_* (salmon in Figure 6) to explore their potential binding modes and interactions. The docking study revealed that the 3,5-dinitroaryl moiety of **12a** adopts a binding pose similar to the uracil moiety of nucleoside inhibitors in the uracil binding sub-pocket and makes the crucial π–π stacking interaction with residue F226, as observed for compound **1** (Figure 6A). Hydrogen bond interactions of the 3,5-dinitroaryl moiety with L174, D172, and K44 were also observed, while the alkyl chain was positioned into the hydrophobic groove. In the uridine-adjacent sub-pocket, the triazole core of **12a** makes a hydrogen bond with N169, which is an important residue known to engage in binding interactions within the uridine-adjacent sub- pocket. However, the aminopropyl moiety attached to the N1 atom of the 1,2,4-triazole ring formed a hydrogen bond interaction and an ionic interaction with D229 in the Mg^2+^ sub-pocket instead of with T49 or G228 in the uridine-adjacent sub-pocket, as it had originally been intended. This interaction was absent with compound **1**, which may explain the enhanced inhibitory activity of **12a** relative to compound **1**. Given that D229, D93, and D94 define the Mg^2+^ cofactor binding site in MraY, we hypothesized that compound **12a** could enable electrostatic interaction between the positively charged amine of the aminopropyl group and H-bond interaction with the Mg^2+^ cofactor binding site. This most likely account for the improved inhibitory activity of **12a** (IC_50_ = 25 µM) against MraY compared to **1** (IC_50_ = 171 µM). Although compound **13** with a piperidine ring attached to N1 of the 1,2,4-triazole ring had a similar binding pose to compound **12a** (Figure 6B) containing an aminopropyl group, the MraY activity of compound **13** (IC_50_ = ∼100 µM) was 4-fold less than compound **12a** and more comparable to compound **1** (IC_50_ = 171 µM). The reason being is the crucial interactions between the aminopropyl moiety of compound **12a** and D229 were not observed in the case compound **13**. Therefore, the improved activity of compound **13** (IC_50_ = ∼100 µM) over compound **1** (IC_50_ = 171 µM) may be attributed to the interactions between the piperidine moiety of compound **13** and loop A of MraY*_SA_*, which we were unable to visualize in our system because the loop A region was removed when we built the protein model based on AlphaFold. Compound **17** exhibited a distinct docking pose (Figure 6C). Docking studies indicated that the π–π stacking interaction between F226 and the 3,5-dinitrophenyl moiety was preserved. Hydrogen bond interactions of the 3,5-dinitroaryl moiety with L174 and D172 were also observed. However, due to the bulky aminoribosyl group of **17**, the 3,5-dinitrophenyl moiety was oriented in the opposite direction, leading to a new hydrogen bond interaction with T49 and the absence of interaction with K44 that was observed in the case with compounds **1** and **12a**. The 5-aminoribosyl unit of compound **17** was observed to interact with N169 rather than the triazole core as in other compounds. Interactions of N166 and A167 with the alkyl chain in the hydrophobic sub-pocket were not observed. These resulted from the 1,2,4-triazole core and the aliphatic chain being further shifted towards the TM9b/Loop E hotspot due to the bulky aminoribosyl group of compound **17.**

**Figure 6.**
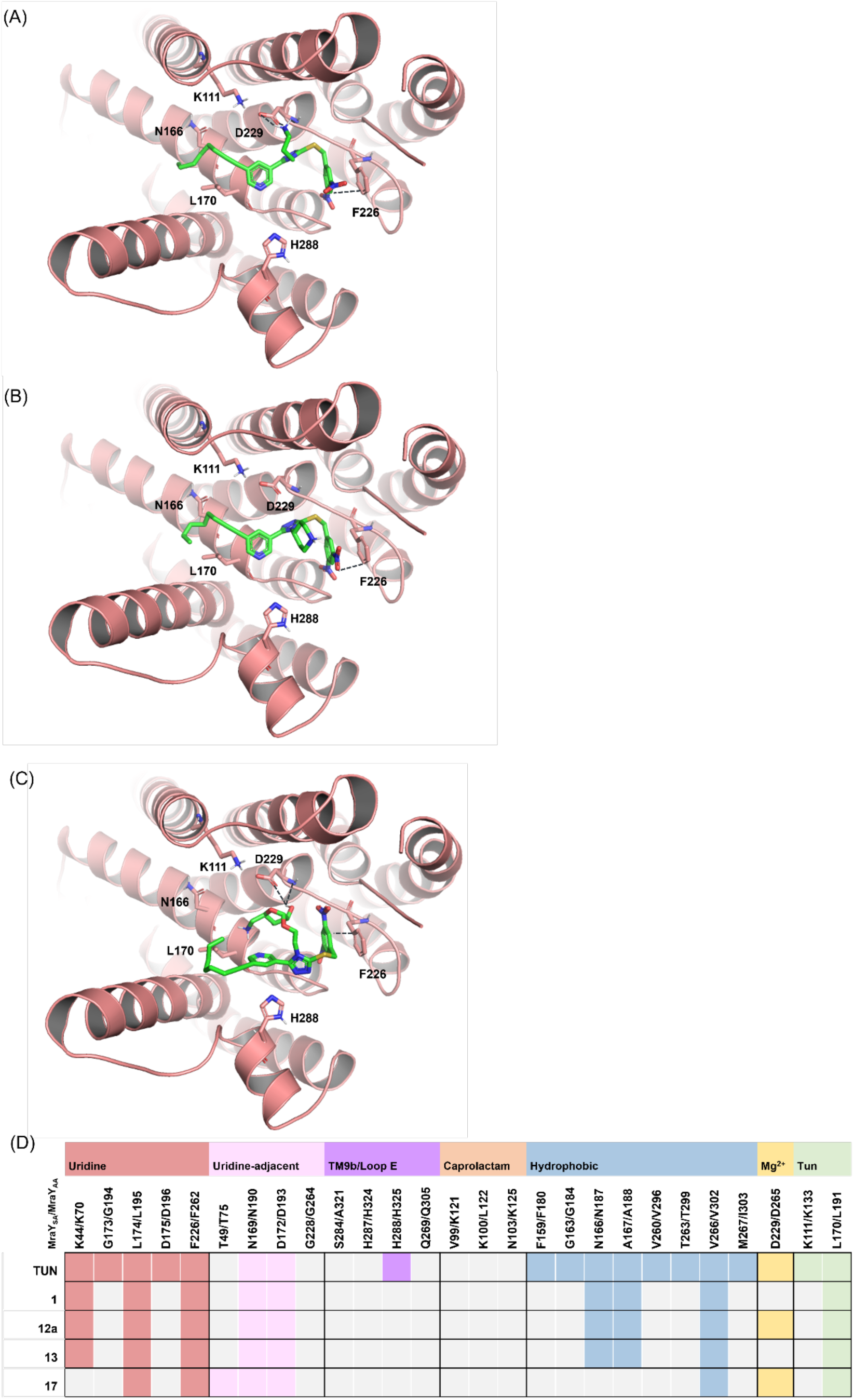
Binding mode of **(A) 12a**, **(B) 13**, and **(C) 17** in the binding site of MraY*_SA_* as predicted using AutoDock Vina.^41,42^ **(D)** Table summarizing the interactions between MraY*_SA_* and compounds **1**, **12a**, **13**, **17,** and **TUN**. Amino acid residue numbers are based on the MraY*_SA_*/MraY*_AA_* sequence alignment. The color-coded squares indicate interactions between specific amino acid residues and the inhibitors, with light gray squares denoting the absence of an interaction. The geometric criteria for interactions are listed in Figure 3.

The docking studies demonstrated the binding modes and the relevant interactions between the compounds and MraY active site. It demonstrated that the 3,5-dinitroaryl moiety of compounds **12a**, **13**, and **17** formed crucial π-π stacking interactions with F226. The interaction between the aminopropyl moiety of compound **12a** and D229 may explain its improved MraY inhibition activity. This provides us valuable insights into their binding modes, which is their primary application.

### In vitro antibacterial activity

We next evaluated the antibacterial profiles of the newly synthesized MraY inhibitors against the ESKAPE panel comprising *Enterococcus faecium*, *Staphylococcus aureus*, *Klebsiella pneumoniae*, *Acinetobacter baumannii*, *Pseudomonas aeruginosa*, and *Enterobacter* spp. The urgent need for effective therapies against these pathogens is becoming increasingly critical. Compounds **12a-e**, **13**, and **17** were selected for screening due to their promising MraY*_SA_* inhibitory activity, and their minimum inhibitory concentration (MIC) values are reported in **Table 3**. Among the analogs screened, compound **12d** showed the best antibacterial profile with MIC of 4 μg/mL against *E. faecalis*, *S. aureus* ATCC 29213, MRSA, and VRSA. Compound **12b** showed MIC values of 4 μg/mL against MRSA and VRSA with 8 μg/mL against *E. faecalis* and S. *aureus* ATCC 29213. Similar MIC values of 8 μg/mL were shown by analogous **12a**, **12c,** and **13** against *S. aureus* ATCC 29213, MRSA, and VRSA. At the same time, **12c** produced better MIC (8 μg/mL) against *E. faecalis* compared to **12a** and **13** with MIC values of 16 μg/mL. Compounds **12e** and **17** showed no to very little antibacterial activity overall.

**Table 3.**
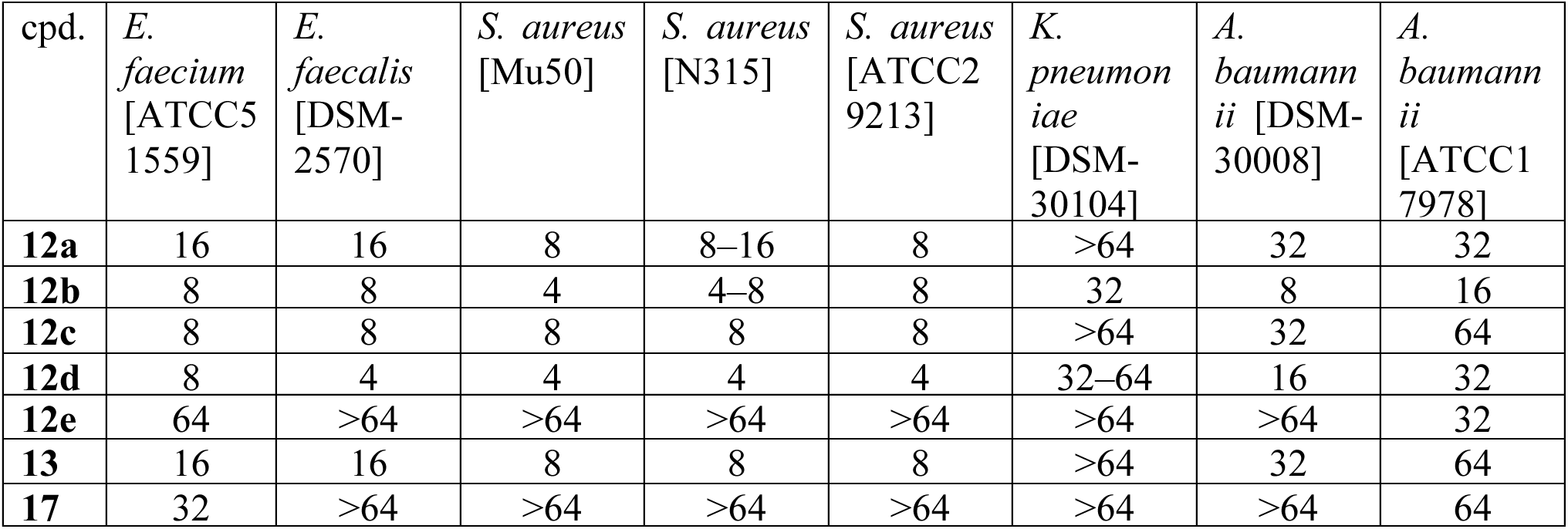
Antibacterial activities (MIC (μg/mL)) of tested compounds against the ESKAPE panel.

| cpd. | *E. faecium*<br>[ATCC51559] | *E. faecalis*<br>[DSM-2570] | *S. aureus*<br>[Mu50] | *S. aureus*<br>[N315] | *S. aureus*<br>[ATCC29213] | *K. pneumoniae*<br>[DSM-30104] | *A. baumannii* [DSM-30008] | *A. baumannii* [ATCC17978] |
| --- | --- | --- | --- | --- | --- | --- | --- | --- |
| **12a** | 16 | 16 | 8 | 8–16 | 8 | >64 | 32 | 32 |
| **12b** | 8 | 8 | 4 | 4–8 | 8 | 32 | 8 | 16 |
| **12c** | 8 | 8 | 8 | 8 | 8 | >64 | 32 | 64 |
| **12d** | 8 | 4 | 4 | 4 | 4 | 32–64 | 16 | 32 |
| **12e** | 64 | >64 | >64 | >64 | >64 | >64 | >64 | 32 |
| **13** | 16 | 16 | 8 | 8 | 8 | >64 | 32 | 64 |
| **17** | 32 | >64 | >64 | >64 | >64 | >64 | >64 | 64 |

#### In vitro anti-*tuberculosis* activity

Since the starting compound **1** had been identified from our prior work as a selective growth inhibitor of *Mtb* (IC_50_ = 0.34 µM),^38^ we also tested all novel analogs in this study for inhibition of the wild-type (WT) *Mtb* Erdman strain using the microplate Alamar Blue assay (MABA). The MABA utilizes the dye resazurin, which turns from dark blue and nonfluorescent to pink and fluorescent when it is reduced to resorufin as a result of cellular metabolism.^47,48^ Compounds that inhibit the *Mtb* growth or survival will decrease or block the color and fluorescence changes. The IC_50_ values for the growth of *Mtb* of all the compounds are summarized in **Table 4**. The IC_50_ value of compound **5a-5f** against *Mtb* was previously determined and reported in our earlier work.^38^ Analogs **5b** and **5c** increased the IC_50_ >15-fold relative to **1.** However, compound **5e** with a nitrofuran group displayed comparable inhibition activity. Meanwhile, compound **5f** was inactive against *Mtb*. There was also a 10-fold reduction in the antitubercular activity of **12a** (IC_50_ of 2.9 µM) compared to that of compound **1** (IC_50_ of 0.34 µM). Analog **8e**, with a nitrofuran moiety in the eastern region, displayed ∼10-fold better anti-TB activity when compared to the aminopropyl analog (**12e**). **13** showed similar inhibition activity compared to compounds **1** and **12a**. At the same time, compound **17** displayed a six-fold reduction in antitubercular activity with an IC_50_ of 19 µM, which is lower than compound **12a** for *Mtb* (IC_50_ of 2.9 µM) inhibition. We think this might be due to its lower inhibitory activity against MraY, possibly in combination with reduced cellular uptake due to the polar aminoribosyl unit. Compound **18,** with acetonide protection of the ribosyl group, showed moderate activity against *Mtb* (IC_50_ of 26 µM). However, compounds **21**, **22,** and **23** containing the uridine moiety were mostly inactive against *Mtb* in the MABA assay, with IC_50_ values of 98 µM, 196 µM, and > 200 µM, respectively. This loss of activity is perhaps due to the polar nature of the uridine unit reflected in the ClogD 7.4, which could lead to poor penetration across the complex mycobacterial cell envelope. The limited correlation between the anti-*Mtb* activity data and the MraY inhibition data suggests (i) that differences in cellular uptake might also play an important role and (ii) that compound **1** and this series of novel compounds may have additional relevant mycobacterial targets other than MraY. Other factors, such as intracellular stability and efflux mechanisms, could also influence the observed outcomes in anti-*Mtb* activity.

**Table 4.**
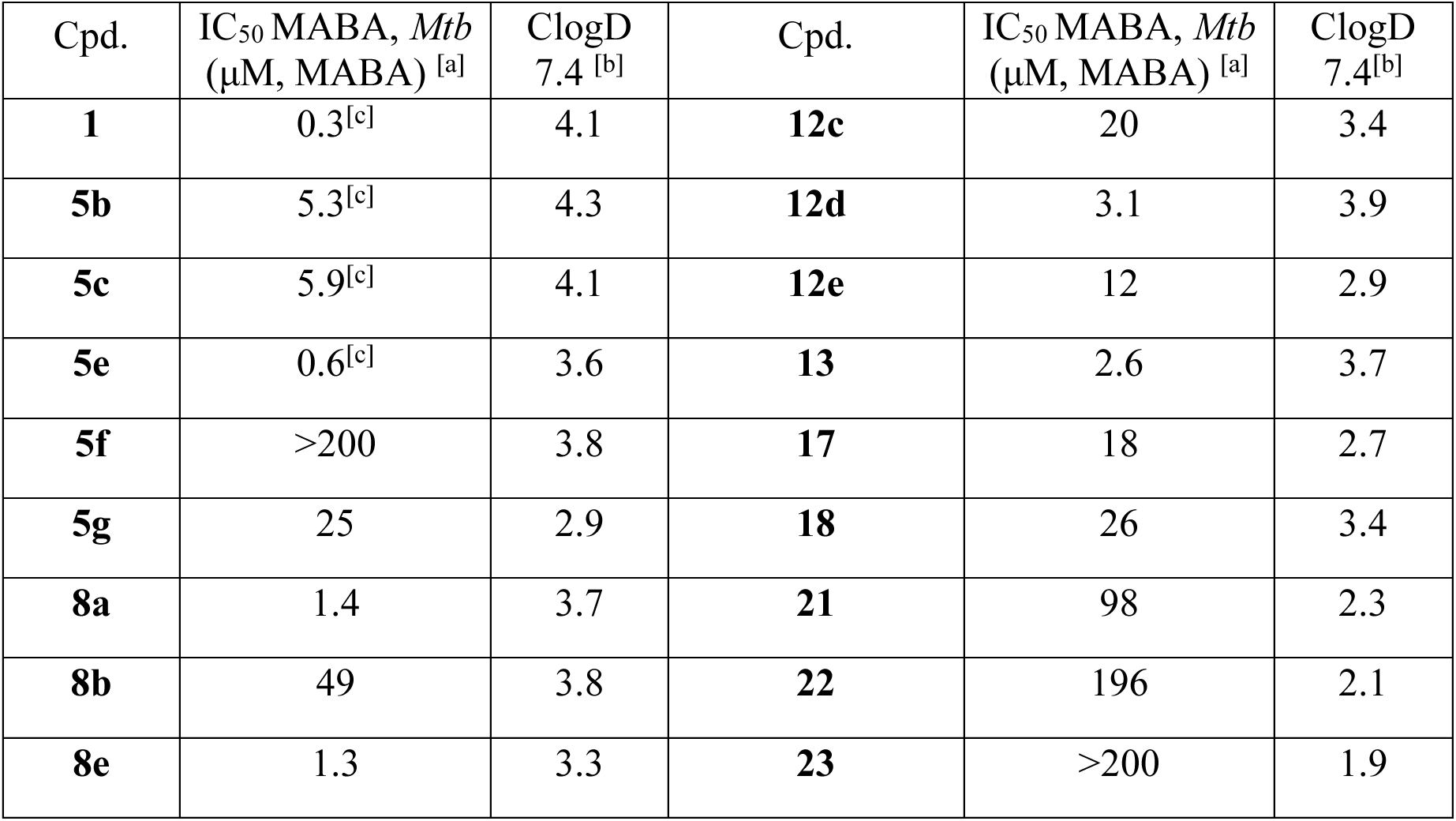
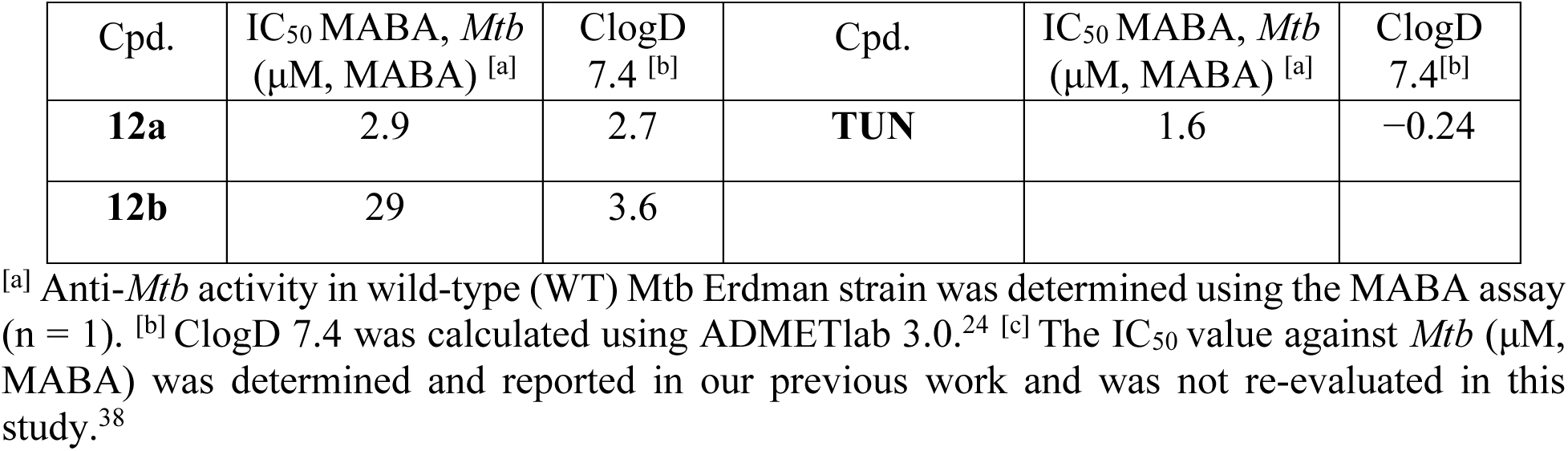
Anti-*Mtb* activities of selected compounds and their ClogD.

## CONCLUSION

In summary, our proof-of-concept study successfully led to the discovery of first-in-class small molecule inhibitors of the MraY protein that are non-nucleosides. We systematically used the structural information available for the MraY active site in our inhibitor design which led us to identify the most potent non-nucleoside inhibitor reported to date, **12a** with an IC_50_ of 25 µM against MraY*_SA_*. This activity is better than that of phloxine B, halogenated fluorescein with antibacterial properties with a reported IC_50_ of 32 µM against MraY from *E. coli*.^31^ Molecular docking studies of compound **12a** and selected analogs of this series suggest interactions with the uridine binding sub-pocket engaging in π–π stacking interactions with the 3,5-nitrophenyl unit, the uridine-adjacent sub-pocket involved in hydrogen-bond interactions, and the hydrophobic sub- pocket with the long aliphatic tail. Although the docking study shed light on the potential binding poses and key interactions between the inhibitors and MraY*_SA_*, the small differences between the docking scores of the compounds hindered us from correlating the docking scores to their inhibition activity against MraY. Therefore, the development of robust computational strategies and tools are required to accurately predict the inhibition activity of newly designed compounds against MraY. Additionally, compounds **12e** and **12b** showed broad spectrum *in vitro* inhibition activity against different strains of *E. faecium* and *S. aureus* including MRSA and VRSA, displaying their potential to serve as structural templates for future development towards antibacterial drug candidates. This series of compounds was also screened against *Mtb*. Even though **12a** had improved MraY inhibitory activity of IC_50_ = 25 µM in the biochemical assays relative to the other analogs, this did not translate to nanomolar potency against *Mtb*. Overall, this series of MraY inhibitors have broad-spectrum antibacterial activity, which is very encouraging. This study presents the very first example of MraY inhibitors that are devoid of the nucleoside moiety, easy to synthesize, and show more drug-like properties compared to their nucleoside-derived counterparts. This novel class of 1,2,4-triazole-based inhibitors thus provides a promising structural framework for development of future antibacterial agents targeting MraY protein as well as against ESKAPE pathogens and *Mtb*. Future directions include further optimization to improve potency, to evaluate and improve drug-like properties as well as PK/PD parameters in order to obtain potential candidates for antibacterial drug development.

## METHODS

### Computational Study

#### Ligand preparation

The 3D structures of the ligands were built in Maestro and then prepared in LigPrep, using the OPLS4 force field.^49,50^ Ionization states of the ligands were generated using Epik (Schrodinger Release 2020-1: Epik, 2020) at physiological pH (7.4).^51^

#### Molecular Docking

Ligands were initially docked to MraY using the program Glide as previous reported docking method.^35^ However, the best scoring docked poses for nearly all ligands lacked π–π stacking interactions between F256 and 4-chlorophenyl ring, thought to be essential for ligand binding to MraY. Therefore, we used AutoDock Vina (ADV) for docking calculations in this study which portrayed the π–π stacking interactions.^41,42^ The MraY and ligand PDB structures were converted into PDBQT format using the DockingPie plugin of PyMOL.^52,53^ A conformational search space cube of dimension 25 Å X 25 Å X 25 Å centered on the Gly186 residue was used for docking with ADV.^42^ An exhaustiveness of 20 and an energy range cut-off of 2 kcal·mol −1 were employed. A total of 20 docking poses were generated for each ligand and then visualized and analyzed using PyMOL.^53^ The best scoring pose of each ligand was selected

### Chemistry

#### General Methods and Instruments

All solvents and reagents were purchased from standard commercial vendors and used without further purification. Synthetic reactions were monitored using thin-layer chromatography (TLC) (Sorbtech silica XG TLC plates) and visualized under UV at 254 nm or with appropriate staining. Purification was performed using medium pressure liquid chromatography (MPLC) on a Biotage Isolera One using Biotage SNAP 10 g–50 g cartridges. NMR spectra were recorded on a Bruker Avance-500 or Bruker Avance-400 spectrometer at 298.15 K. Chemical shifts are reported in ppm using deuterated solvents (CDCl_3_, CD_3_OD, or DMSO-d_6_) for ^1^H and ^13^C NMR. CDCl_3_ (δ = 77.16 ppm), CD_3_OD (δ = 49.00 ppm) or DMSO-*d*_6_ (δ = 39.52 ppm) were used as internal standards for ^13^C NMR. For ^1^H NMR, CDCl_3_ (δ = 7.26 ppm), CD_3_OD (δ = 3.31 ppm) or DMSO-*d*_6_ (δ = 2.50 ppm) or TMS (δ = 0 ppm) were used as internal standards. Data were reported as: s = singlet, br = broad singlet, d = doublet, t = triplet, q = quartet, p = pentet, m = multiplet, b = broad, ap = apparent; coupling constants, *J*, in Hz. For high-resolution mass spectrometry (HRMS), a quadruple-TOF was used to obtain the data both in positive or negative modes. Purity (>95%) was determined using a Dionex Ultimate 3000 UPLC system (Thermo Fisher Scientific).

#### General procedure for 12a and 13

The appropriate halide alkylating agent (1.2 equiv) was added to a solution of the corresponding 1,2,4-triazole-3-thiol (1.0 equiv) and NEt_3_ (2.0 equiv) in CH_3_CN (0.1 M) and stirred overnight at room temperature. Upon completion of the reaction, the solvent was removed under reduced pressure. The residue was then dissolved in EtOAc and washed with water and brine. The extracted organic layer was dried over anhydrous Na_2_SO_4_ and concentrated under low pressure. The resultant residue was purified by MPLC. The solution of Boc-protected substrate in 2.5 N HCl in ethanol (1.5 mL) was stirred at room temperature or for compounds 4 h. The reaction mixture was neutralized with NaHCO_3_ and extracted with CH_2_Cl_2_. The organic phase was washed with water, dried over Na_2_SO_4_, and evaporated under reduced pressure. The residue was purified by MPLC (1–10% MeOH/CH_2_Cl_2_). The solution of corresponding Boc-protected substrate in 2.5 N HCl in ethanol (1.5 mL) was stirred at room temperature or for compounds 4 h. The reaction mixture was neutralized with NaHCO_3_ and extracted with CH_2_Cl_2_. The organic phase was washed with water, dried over Na_2_SO_4_, and evaporated under reduced pressure. The residue was purified by MPLC (1–10% MeOH/CH_2_Cl_2_).

### 3-(3-((3,5-Dinitrobenzyl)thio)-5-(5-(oct-1-yn-1-yl)pyridin-3-yl)-4H-1,2,4-triazol-4-yl)propan-1-amine (12a)

The general procedure was used to synthesize compound **12a**, which was obtained as a brown sticky solid in 40% yield. ^1^H NMR (400 MHz, CDCl_3_) δ 8.88 (t, *J* = 2.1 Hz, 1H), 8.73–8.63 (m, 3H), 8.61 (d, *J* = 2.0 Hz, 1H), 7.92 (t, *J* = 2.1 Hz, 1H), 4.71 (s, 2H), 4.06 (t, *J* = 7.6 Hz, 2H), 3.6 (brs, 2H) 2.76 (t, *J* = 6.8 Hz, 2H), 2.39 (t, *J* = 7.2 Hz, 2H), 1.89 (p, *J* = 7.1 Hz, 2H), 1.58 (p, *J* = 7.2 Hz, 2H), 1.47–1.20 (m, 6H), 0.93–0.83 (m, 3H). ^13^C NMR (101 MHz, CDCl_3_) δ 153.4, 153.1, 150.7, 148.5, 146.4, 141.9, 138.7, 129.6, 123.0, 122.0, 118.2, 96.5, 76.3, 42.7, 38.2, 35.4, 31.6, 31.4, 28.7, 28.5, 22.6, 19.6, 14.1. HRMS (ESI): *m/z* [M + H]^+^ Calcd for [C_25_H_29_N_7_O_4_S + H]^+^ 524.2080, found 524.2087.

### 3-(5-((3,5-Dinitrobenzyl)thio)-4-(piperidin-4-yl)-4H-1,2,4-triazol-3-yl)-5-(oct-1-yn-1- yl)pyridine (13)

The general procedure was used to synthesize compound **13**, which was obtained as a brown solid in 42% yield. ^1^H NMR (500 MHz, CDCl_3_) δ 8.9 (d, *J* = 2.3 Hz, 1H), 8.7 (dd, *J* = 6.5, 2.1 Hz, 3H), 8.5 (d, *J* = 2.1 Hz, 1H), 7.8 (d, *J* = 2.3 Hz, 1H), 4.8 (s, 2H), 4.1 (tt, *J* = 12.5, 4.3 Hz, 1H), 3.3–3.2 (m, 2H), 2.7–2.6 (m, 2H), 2.5–2.3 (m, 4H), 1.8 (dd, *J* = 13.0, 3.6 Hz, 2H), 1.6 (p, *J* = 7.2 Hz, 2H), 1.4 (p, *J* = 7.2 Hz, 2H), 1.4–1.2 (m, 4H), 0.9 (t, *J* = 6.9 Hz, 3H). ^13^C NMR (126 MHz, CDCl_3_) δ 153.8, 153.5, 149.2, 148.6, 147.2, 142.0, 139.2, 129.6, 123.2, 122.0, 118.2, 96.5, 76.4, 55.6, 45.7, 35.7, 31.4, 31.3, 28.7, 28.5, 22.6, 19.6, 14.2. HRMS (ESI): *m/z* [M - H]^-^ Calcd for [C_27_H_31_N_7_O_4_S - H]^−^ 548.2080, found 548.2094.

### 2-(Aminomethyl)-5-(2-(3-((3,5-dinitrobenzyl)thio)-5-(5-(oct-1-yn-1-yl)pyridin-3-yl)-4H- 1,2,4-triazol-4-yl)ethoxy)tetrahydrofuran-3,4-diol (17)

To compound **15** in a 1-gram vial was added 1 mL of 80% TFA in dichloromethane and stirred for 2 h. The solvent was removed under reduced pressure and the residue purified by PTLC 15% MeOH/CH_2_Cl_2._ Compound **17** was obtained in 40% yield. The synthesis and characterization of compound **15** can be found in the supporting information. ^1^H NMR (400 MHz, MeOD) δ 8.90 (t, *J* = 2.2 Hz, 1H), 8.83–8.66 (m, 4H), 8.13 (s, 1H), 4.76–4.64 (m, 3H), 4.24 (t, *J* = 5.4 Hz, 2H), 4.01–3.87 (m, 2H), 3.85–3.75 (m, 1H), 3.73 (d, *J* = 4.5 Hz, 1H), 3.68–3.59 (m, 1H), 3.16–3.09 (m, 1H), 2.61 (dd, *J* = 13.0, 9.8 Hz, 1H), 2.49 (t, *J* = 7.1 Hz, 2H), (p, *J* = 8.2, 7.6, 6.9 Hz, 2H)1.55–1.43 (m, 2H), 1.41–1.29 (m, 4H), 0.96–0.88 (m, 3H). HRMS (ESI): *m/z* [M + H]^+^ Calcd for [C_29_H_35_N_7_O_8_S + H]^+^ 642.2346, found 642.2350.

The synthesis and characterization of the remaining compounds and intermediates can be found in the supporting information.

### Biochemistry & Microbiology

#### Fluorescence-based MraY assay

Fluorescence intensity was determined over time at a specific wavelength (355 nm excitation and 520 nm emission) using a BMG Labtech POLARstar Omega plate reader with 384 wells. MraY-catalyzed reaction was initiated by adding a mixture of crude membrane preparation of MraY from *S. aureus* (1 mL), undecaprenyl phosphate (50 mm), and dansylated Park’s nucleotide 3 (synthetic or semi-synthetic, 7.5 mm) to a buffer solution (100 mm Tris-HCl buffer pH 7.5, 200 mm KCl, 10 mm MgCl_2_, 0.1% Triton X-100, 20 mL total volume). Membrane preparations from non-transfected *E. coli* Lemo21 cells were used as negative controls. To demonstrate partial inhibition of MraY activity, the known MraY inhibitor, tunicamycin (Sigma-Aldrich), was added to the reaction mixture at a concentration of 100 nm.

#### Microplate Alamar Blue Assay (MABA)

WT *Mtb* Erdman or Λ1*cydA*^48^ *Mtb* was cultured in Middlebrook 7H9 liquid media supplemented with 60 μL/L oleic acid, 5 g/L bovine serum albumin, 2 g/L dextrose, 0.003 g/L catalase (OADC), 0.5% glycerol, and 0.05% Tween 80 at 37°C The bacteria were inoculated at a final OD_600_ of 1.6 x 10^-3^ in 200µL per well in 96-well plates with two-fold titrations of compounds. The concentration of DMSO was maintained at 1% for all wells to avoid toxicity. The 96-well plates were incubated in a humidified incubator at 37°C and 5% CO_2_ for one week. After the week incubation, 32.5 μL of a mixture containing an 8:5 ratio of 0.6 mM resazurin (Sigma) dissolved in 1X PBS to 20% Tween 80 was added and the production of fluorescent resorufin was measured after incubation at 37°C in 5% CO_2_ for 24 hours. Relative fluorescence units (RFU) were measured using a BioTek Synergy H1 Microplate Reader H1M at an excitation λ of 530 nm and an emission λ of 590 nm. Media in the absence of bacteria served as the negative control, and media with bacteria in the absence of compound served as the positive control. Percent inhibition was calculated as follows:

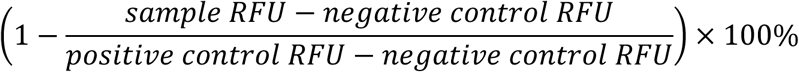

Percent inhibition was input into PRISM GraphPad to calculate IC_50_ values by plotting as a non-linear regression curve.

## NOTES

The authors declare no competing financial interest.

## ACKNOWLEDGMENT

This work was funded by NIH R21 #AI142210 in NIAID primarily and the National Institute of General Medical Sciences of the National Institutes of Health P20GM130460. C.L.S. is supported by a Burroughs Wellcome Fund Investigators in the Pathogenesis of Infectious Disease Award. Support was provided by the University of Mississippi School of Pharmacy. This publication is solely the responsibility of the authors and does not necessarily represent the official view of NIH.

