## Supporting Information for "1,2,4-Triazole-based first-in-class non-nucleoside inhibitors of bacterial enzyme MraY"

### Chemistry Protocols

#### Synthetic Procedure for Intermediates

##### 1. General procedure for synthesis of isothiocyanates (6 and 9)

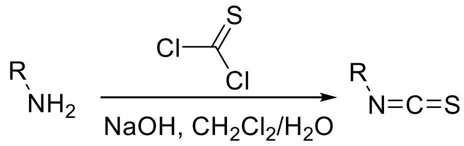

Method 1A: To the corresponding amine (1.0 equiv) dissolved in water (10 mL) and chloroform (10 mL) and brought to 0 °C was added thiophosgene (1.5 equiv) dropwise and NaOH (3 equiv) dissolved in water (5mL). The reaction was brought to rt and stirred for 1.5 h. Water (20 mL) was added to the reaction mixture to quench. The organic phase was dried with anhydrous $Na_2SO_4$ and concentrated under reduced pressure and the residue was purified by MPLC (0–30% EtOAC/Hexane).[1]

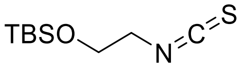

***tert*-Butyl(2-isothiocyanatoethoxy)dimethylsilane (6).** The title compound was synthesized according to method 1A to obtain a white solid (yield = 48%). $^1$H NMR (400 MHz, $CDCl_3$) δ 3.81 (t, *J* = 5.4 Hz, 2H), 3.57 (t, *J* = 5.4 Hz, 2H), 0.91 (s, 9H), 0.10 (s, 6H). Spectroscopic data is consistent with previously reported data for this compound.[2]

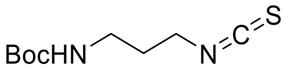

***tert*-Butyl (3-isothiocyanatopropyl)carbamate (9a).** The title compound was synthesized according to method 1A to obtain a yellow solid (yield = 79%). $^1$H NMR (400 MHz, $CDCl_3$) δ

4.69 (s, 1H), 3.56 (t, *J* = 6.5 Hz, 2H), 3.20 (q, *J* = 6.5 Hz, 2H), 1.85 (p, *J* = 6.6 Hz, 2H), 1.40 (s, 10H). $^{13}$C NMR (101 MHz, $CDCl_3$) δ 156.1,130.7, 79.7, 42.8, 37.8, 30.5, 28.5.

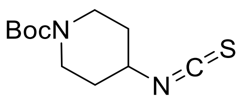

***tert*-Butyl 4-isothiocyanatopiperidine-1-carboxylateYellowish white solid (9b)** The title compound was synthesized according to method 1A to obtain clear liquid (yield = 83%). $^{1}$H NMR (400 MHz, $CDCl_3$) δ 3.88 (tt, *J* = 7.4, 3.8 Hz, 1H), 3.60 (ddd, *J* = 13.9, 7.6, 3.7 Hz, 2H), 3.34 (ddd, *J* = 13.9, 7.5, 3.7 Hz, 2H), 1.89 (ddt, *J* = 14.7, 7.6, 3.7 Hz, 2H), 1.75 (dtd, *J* = 13.6, 7.4, 3.7 Hz, 2H), 1.44 (s, 9H). $^{13}$C NMR (101 MHz, $CDCl_3$) δ 154.6, 132.4, 53.4, 41.0, 32.3, 28.5.

#### 2 General procedure for synthesis of 4,5-substituted-1,2,4-triazole-2-thiones (4, 7, 10 and 11)

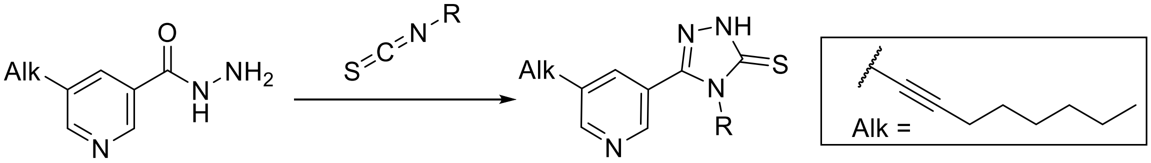

Method 2A: Step1: Acyl hydrazide (1.0 equiv) and corresponding isothiocyanate (1.0 equiv) dissolved in absolute ethanol (10 mL) was refluxed for 4 h. After completion of reaction, as confirmed by TLC, the solvent was removed under reduced pressure to obtain the corresponding 1,4-substituted thiosemicarbazides. Step 2: To the residue was then added 5 mL of 10% aqueous NaOH and stirred for 3 h at 60 °C. Upon completion, the reaction mixture was brought to room temperature and acidified with conc. HCl to a pH of 5–6. The reaction mixture was extracted (x 2) with EtOAc, the organic phase was then washed with brine, dried with anhydrous $Na_2SO_4$, concentrated in vacuo, and purified using MPLC.

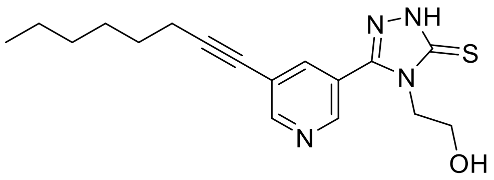

**4-Methyl-5-(5-(oct-1-yn-1-yl)pyridin-3-yl)-2,4-dihydro-3H-1,2,4-triazole-3-thione (4).** The title compound was synthesized according to method 2A to obtain a white solid (yield: 70%). $^1$H NMR (400 MHz, CDCl3) δ 12.80 (s, 1H), 8.86–8.74 (m, 2H), 7.93 (t, *J* = 2.1 Hz, 1H), 3.69 (s, 3H), 2.44 (t, *J* = 7.2 Hz, 2H), 1.61 (q, *J* = 7.4 Hz, 2H), 1.50–1.39 (m, 2H), 1.39–1.24 (m, 4H), 0.89 (t, J = 6.8 Hz, 3H). $^{13}$C NMR (101 MHz, CDCl3) δ 169.0, 154.1, 149.1, 146.6, 138.3, 122.1, 122.1, 96.9, 76.2, 32.4, 31.4, 28.7, 28.4, 22.6, 19.6, 14.2. HRMS (ESI): m/z [M + H] Calcd for [$C_{16}H_{20}N_4S$ + H]$^+$ 301.1487, found 301.1482.

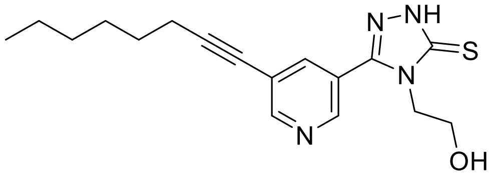

**2-(3-Mercapto-5-(5-(oct-1-yn-1-yl)pyridin-3-yl)-4H-1,2,4-triazol-4-yl)ethan-1-ol (7).** The title compound was synthesized according to method 2A to obtain an off-white solid (yield = 82%). $^1$H NMR (400 MHz, MeOD) δ 8.82 (d, *J* = 2.2 Hz, 1H), 8.68 (d, *J* = 2.1 Hz, 1H), 8.29 (t, *J* = 2.1 Hz, 1H), 4.18 (t, *J* = 5.2 Hz, 2H), 3.94 (t, *J* = 5.2 Hz, 2H), 2.48 (t, *J* = 7.0 Hz, 2H), 1.69–1.57 (m, 2H), 1.55–1.43 (m, 2H), 1.43–1.29 (m, 4H), 0.97–0.87 (m, 3H). $^{13}$C NMR (101 MHz, MeOD) δ 169.35, 154.07, 151.16, 148.77, 140.73, 124.61, 122.98, 96.93, 77.20, 59.22, 32.48, 29.69, 29.51, 23.60, 20.07, 14.39. HRMS (ESI): *m/z* [M + H]$^+$ Calcd for [$C_{17}H_{22}N_4O_{10}S$ + H]$^+$ 331.1593, found 331.1604.

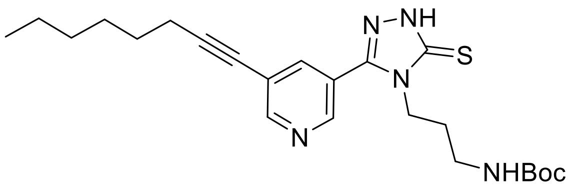

**tert-Butyl (3-(3-(5-(oct-1-yn-1-yl)pyridin-3-yl)-5-thioxo-1,5-dihydro-4H-1,2,4-triazol-4-yl)propyl)carbamate (10).** The title compound was synthesized according to method 2A to obtain an off-white solid (yield = 72%). $^{1}$H NMR (400 MHz, $CDCl_3$) δ 13.06 (s, 1H), 8.86–8.71 (m, 2H), 7.93–7.88 (m, 1H), 5.33–5.25 (m, 1H), 4.22 (t, $J$ = 7.1 Hz, 2H), 3.22–3.03 (m, 2H), 2.43 (t, $J$ = 7.2 Hz, 2H), 1.95–1.79 (m, 2H), 1.61 (p, $J$ = 7.1 Hz, 2H), 1.52–1.36 (m, 11H), 1.36–1.24 (m, 4H), 0.90 (t, $J$ = 6.9 Hz, 3H). $^{13}$C NMR (101 MHz, $CDCl_3$) δ 168.6, 156.2, 154.1, 148.8, 146.5, 138.4, 122.2, 97.0, 79.5, 76.2, 42.4, 37.1, 31.4, 29.3, 28.7, 28.5, 28.4, 22.6, 19.6, 14.2. HRMS (ESI): *m/z* $[M - H]^-$ Calcd for $[C_{23}H_{33}N_5O_2S - H]^-$ 442.2277, found 442.2295.

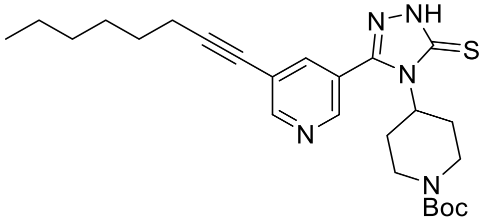

**tert-Butyl 4-(3-mercapto-5-(5-(oct-1-yn-1-yl)pyridin-3-yl)-4H-1,2,4-triazol-4-yl)piperidine-1-carboxylate (11).** The title compound was synthesized according to method 2A to obtain a light-yellow solid (yield = 81%). $^{1}$H NMR (400 MHz, $CDCl_3$) δ 12.22 (s, 1H), 8.80 (d, $J$ = 2.0 Hz, 1H), 8.58 (d, $J$ = 2.2 Hz, 1H), 7.74 (t, $J$ = 2.1 Hz, 1H), 4.59 (s, 1H), 4.33–4.10 (m, 2H), 2.72 (s, 2H), 2.44 (t, $J$ = 7.1 Hz, 2H), 2.29 (s, 2H), 2.20 (s, 2H), 1.79 (d, $J$ = 14.1 Hz, 3H), 1.62 (p, $J$ = 7.2 Hz, 2H), 1.50–1.40 (m, 11H), 1.36–1.28 (m, 4H), 0.93–0.86 (m, 3H). $^{13}$C NMR (101 MHz, $CDCl_3$) δ 167.8, 154.6, 154.1, 148.7, 147.3, 139.4, 122.8, 122.0, 97.1, 80.2, 76.0, 56.6, 43.2, 31.3, 29.6, 28.6,

28.4, 28.3, 22.5, 19.5, 14.1. HRMS (ESI): *m/z* $[M + Cs]^+$ Calcd for $[C_{25}H_{35}N_5O_2S + Cs]^+$ 602.1566, found 602.1587.

##### 3. Synthesis procedure for intermediates (14 and 15)

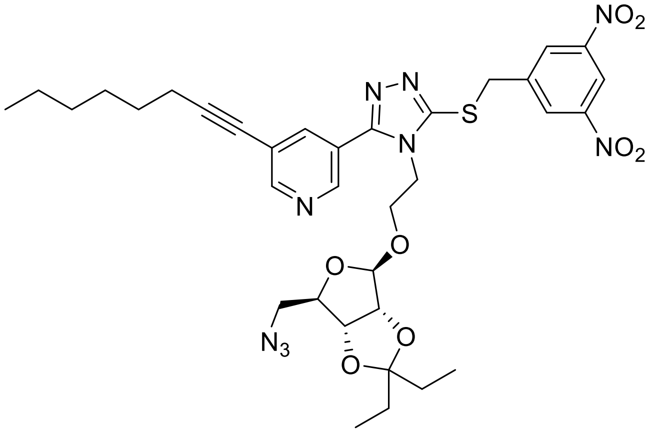

**3-(4-(2-((6-(Azidomethyl)-2,2-diethyltetrahydrofuro[3,4-d][1,3]dioxol-4-yl)oxy)ethyl)-5-((3,5-dinitrobenzyl)thio)-4H-1,2,4-triazol-3-yl)-5-(oct-1-yn-1-yl)pyridine (14).** A mixture of **8a** (72 mg, 1 equiv, 0.14 mmol) and **16a** (35 mg, 1 equiv, 0.14 mmol) in dry $CH_2Cl_2$ (2 mL) was brought to 0 °C. Subsequently, well-dried MS 4Å (500 mg) was added. At every hour, 20 µl of $BF_3.OEt_2$ was added at the same temperature. The reaction was quenched after 6 h with $NaHCO_3$. The mixture was extracted with EtOAc, washed with brine, dried over anhydrous Na2SO4, filtered, and concentrated under reduced pressure and purified by MPLC (mobile phase: 20–50% EtoAC/Hexane). Compound **14** was obtained in 42% yield. $^1$H NMR (500 MHz, $CDCl_3$) δ 8.94 (t, *J* = 2.3 Hz, 1H), 8.77–8.66 (m, 4H), 7.96 (q, *J* = 3.5, 2.9 Hz, 1H), 4.91 (s, 1H), 4.81–4.68 (m, 2H), 4.47 (d, *J* = 6.1 Hz, 1H), 4.39 (d, *J* = 6.0 Hz, 1H), 4.28 (dd, *J* = 8.6, 5.7 Hz, 1H), 4.19–4.07 (m, 2H), 3.95 (ddt, *J* = 13.6, 9.9, 5.1 Hz, 1H), 3.57 (ddd, *J* = 10.8, 6.7, 4.3 Hz, 1H), 3.03 (dd, *J* = 12.7, 6.0 Hz, 1H), 2.95 (dd, *J* = 12.7, 8.2 Hz, 1H), 2.42 (t, *J* = 7.2 Hz, 2H), 1.62 (dq, *J* = 24.0, 7.3 Hz, 4H), 1.53 (q, *J* = 7.4 Hz, 2H), 1.47–1.40 (m, 2H), 1.36–1.27 (m, *J* = 4.6, 4.0 Hz, 4H), 0.93–0.79

(m, 9H). $^{13}$C NMR (126 MHz, $CDCl_3$) δ 153.7, 153.6, 150.6, 148.7, 147.2, 141.9, 138.7, 129.5, 123.1, 121.7, 118.3, 117.5, 109.1, 96.3, 85.9, 85.4, 82.1, 76.4, 65.4, 53.5, 44.6, 35.5, 31.4, 29.4, 28.8, 28.7, 28.5, 22.6, 19.6, 14.2, 8.5, 7.4. HRMS (ESI): *m/z* $[M + H]^+$ Calcd for $[C_{34}H_{41}N_9O_8S + H]^+$ 736.2877, found 736.2886.

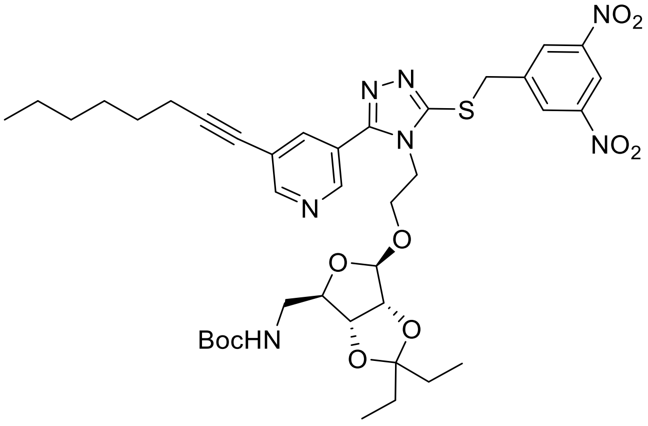

***tert*-Butyl ((6-(2-(3-((3,5-dinitrobenzyl)thio)-5-(5-(oct-1-yn-1-yl)pyridin-3-yl)-4H-1,2,4-triazol-4-yl)ethoxy)-2,2-diethyltetrahydrofuro[3,4-d][1,3]dioxol-4-yl)methyl)carbamate (15)**

To a solution of compound **14** (35 mg, 1.0 equiv, 48 μmol) in THF/toluene (1:1, 2 mL), triphenylphosphine (25 mg, 2 equiv, 95 μmol) and water (43 mg, 43 μL, 50 equiv, 2.4 mmol) were added and the mixture was stirred at room temperature for 12 h. Afterward, Di-*tert*-butyl dicarbonate (21 mg, 21 μL, 2.0 equiv, 95 μmol) and sodium bicarbonate (8.0 mg, 2 equiv, 95 μmol) were added and the mixture was stirred at rt for 2 h. It was then diluted with EtOAc (50 mL), washed with water (50 mL) and brine (50 mL) and dried over anhydrous $Na_2SO_4$. The solvent was evaporated under reduced pressure, and the resultant crude product was purified by Preparative thin layer chromatography (PTLC) (50% EtOAc). Compound 15 was obtained in 67% yield. PTLC was used instead of column chromatography because triphenylphosphine oxide by-product was eluted alongside the title compound during column chromatography. $^1$H NMR (500 MHz, $CDCl_3$) δ 8.94 (t, $J$ = 2.2 Hz, 1H), 8.79–8.66 (m, 4H), 7.98 (d, $J$ = 2.1 Hz, 1H), 4.86 (s, 1H),

4.81–4.65 (m, 3H), 4.52 (d, *J* = 6.0 Hz, 1H), 4.41 (d, *J* = 6.1 Hz, 1H), 4.24–4.14 (m, 2H), 4.10–4.02 (m, 1H), 3.94–3.85 (m, 1H), 3.58–3.46 (m, 1H), 3.03 (t, *J* = 6.6 Hz, 2H), 2.42 (t, *J* = 7.2 Hz, 2H), 2.03 (s, 1H), 1.68–1.57 (m, 4H), 1.52 (q, *J* = 7.6 Hz, 2H), 1.40 (s, 11H), 1.35–1.27 (m, 5H), 0.95–0.73 (m, 10H). $^{13}$C NMR (126 MHz, $CDCl_3$) δ 155.8, 153.7, 153.6, 150.7, 148.7, 147.2, 142.0, 138.8, 129.6, 123.1, 121.7, 118.3, 117.2, 108.6, 96.2, 86.7, 85.4, 82.1, 79.8, 76.6, 65.0, 44.6, 43.6, 35.5, 31.4, 29.8, 29.4, 28.8, 28.8, 28.5, 28.5, 22.6, 19.6, 14.2, 8.5, 7.4. HRMS (ESI): *m/z* $[M + H]^+$ Calcd for $[C_{39}H_{51}N_7O_{10}S + H]^+$ 810.3496, found 810.3483.

**4. General procedure for intermediates (19 and 20)**

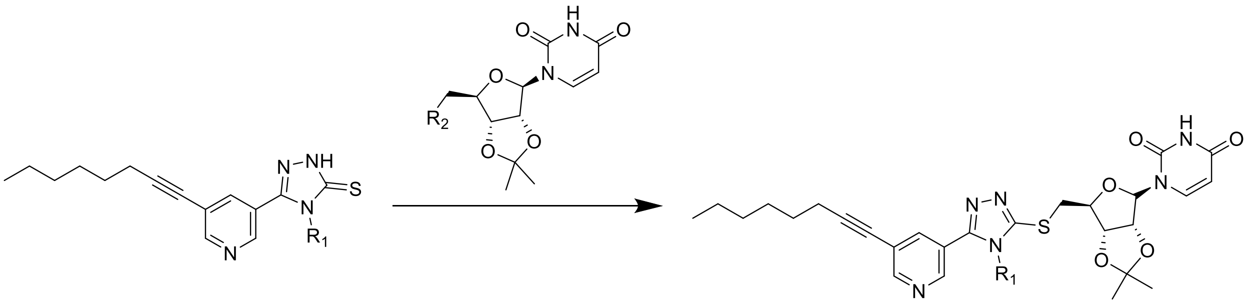

Method 4A: The appropriate halide alkylating agent (1.2 equiv) was added to a solution of the corresponding 1,2,4-triazole-3-thiol (1.0 equiv) and $NEt_3$ (2.0 equiv) in $CH_3CN$ (0.1 M) and stirred overnight at room temperature. Upon completion of the reaction, the solvent was removed under reduced pressure. The residue was then dissolved in EtOAc and washed with water and brine. The extracted organic layer was dried over anhydrous $Na_2SO_4$ and concentrated under low pressure. The resultant residue was purified by MPLC.

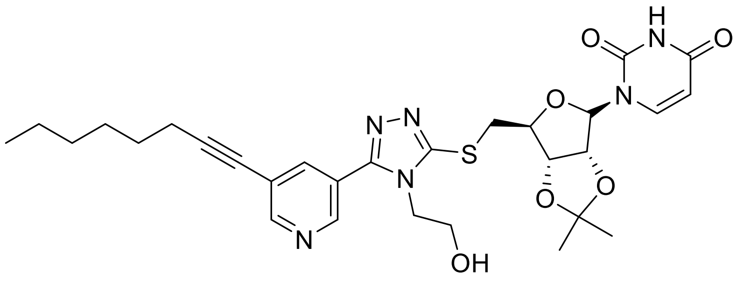

**1-((3aR,4R,6S,6aS)-6-(((4-(2-hydroxyethyl)-5-(5-(oct-1-yn-1-yl)pyridin-3-yl)-4H-1,2,4-triazol-3-yl)thio)methyl)-2,2-dimethyltetrahydrofuro[3,4-d][1,3]dioxol-4-yl)pyrimidine-2,4(1H,3H)-dione (19).** The title compound was synthesized according to method 4A to obtain a white solid (yield = 44%). $^{1}$H NMR (400 MHz, MeOD) δ 8.8 (d, $J$ = 2.1 Hz, 1H), 8.7 (d, $J$ = 2.1 Hz, 1H), 8.2 (t, $J$ = 2.1 Hz, 1H), 7.6 (d, $J$ = 7.9 Hz, 1H), 5.7–5.6 (m, 2H), 5.1 (dd, $J$ = 6.5, 2.0 Hz, 1H), 4.4 (td, $J$ = 6.8, 4.0 Hz, 1H), 4.2 (q, $J$ = 5.2 Hz, 2H), 3.8 (t, $J$ = 5.3 Hz, 2H), 3.6 (dd, $J$ = 13.6, 7.2 Hz, 1H), 3.6 (dd, $J$ = 13.6, 6.5 Hz, 1H), 2.5 (t, $J$ = 7.0 Hz, 2H), 1.69–1.58 (m, 2H), 1.6–1.4 (m, 5H), 1.4–1.3 (m, 7H), 1.0–0.9 (m, 3H). $^{13}$C NMR (101 MHz, $CDCl_3$) δ 163.4, 153.7, 150.0, 147.4, 142.5, 139.3, 122.2, 121.8, 114.8, 102.8, 96.6, 94.8, 86.3, 84.4, 83.0, 76.2, 59.2, 48.3, 47.3, 32.3, 31.3, 28.6, 28.4, 27.1, 25.2, 22.5, 19.5, 14.1. HRMS (ESI): *m/z* $[M + H]^+$ Calcd for $[C_{29}H_{36}N_6O_6S + H]^+$ 597.2495, found 597.2488.

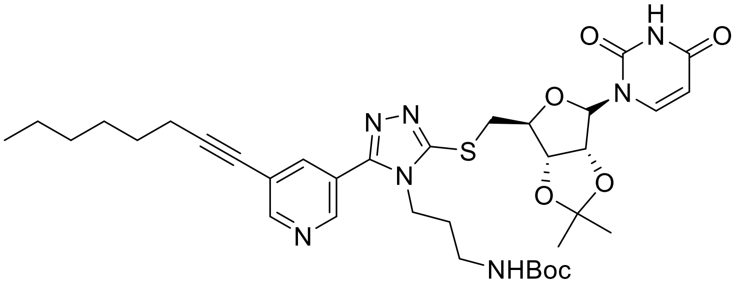

***tert*-Butyl (3-(3-((((3aS,4S,6R,6aR)-6-(2,4-dioxo-3,4-dihydropyrimidin-1(2H)-yl)-2,2-dimethyltetrahydrofuro[3,4-d][1,3]dioxol-4-yl)methyl)thio)-5-(5-(oct-1-yn-1-yl)pyridin-3-yl)-4H-1,2,4-triazol-4-yl)propyl)carbamate (20).** The title compound was synthesized according to method 4A to obtain an off-white solid (yield = 25%). $^{1}$H NMR (400 MHz, $CDCl_3$) δ 9.74 (s, 1H), 8.71 (dd, $J$ = 13.9, 2.1 Hz, 2H), 7.94 (t, $J$ = 2.1 Hz, 1H), 7.27 (d, $J$ = 7.6 Hz, 1H), 5.74 (d, $J$ = 8.1 Hz, 1H), 5.53 (s, 1H), 5.14 (q, $J$ = 6.5 Hz, 1H), 5.01–4.84 (m, 1H), 4.47 (td, $J$ = 6.5, 4.0 Hz, 1H), 4.02 (dt, $J$ = 18.2, 7.6 Hz, 2H), 3.81–3.60 (m, 1H), 3.09 (h, $J$ = 10.3, 8.9 Hz, 2H), 2.44 (t, $J$ = 7.1 Hz, 2H), 1.93–1.80 (m, 2H), 1.63 (p, $J$ = 7.1 Hz, 2H), 1.53 (s, 3H), 1.50–1.38 (m, 11H), 1.37–

1.29 (m, 7H), 0.95–0.88 (m, 3H). $^{13}C$ NMR (101 MHz, $CDCl_3$) δ 163.47, 153.52, 152.60, 150.21, 146.52, 143.31, 138.57, 138.45, 123.26, 121.94, 114.65, 102.90, 96.32, 96.29, 86.58, 84.59, 83.68, 76.52, 50.85, 42.75, 37.55, 35.07, 31.41, 30.82, 28.71, 28.48, 28.45, 27.16, 25.37, 22.64, 19.60, 14.17.

**Synthetic Procedure for Final Analogs**

**5. General procedure for alkylation of 4,5-disubstituted-1,2,4-triazole-2-thiones (5, 8, 12 and 13)**

Method 5A: The appropriate alkylating agent (bromide or chloride) (1.2 equiv) and the corresponding 1,2,4-triazole-3-thione (1.0 equiv) were dissolved in 1:2 parts of MeOH/acetone (0.1 M) at room temperature. Upon addition of $K_2CO_3$ (1.5 equiv), the mixture was stirred overnight at room temperature. The completion of the reaction was monitored by TLC, after which the solvent was evaporated under reduced pressure. The residue was dissolved with EtOAc, washed with water (x2) and brine. The extracted organic layer was dried over anhydrous $Na_2SO_4$, and then concentrated under low pressure. The resultant residue was then purified by MPLC.

Method 5B: The appropriate halide alkylating agent (1.2 equiv) was added to a solution of the corresponding 1,2,4-triazole-3-thiol (1.0 equiv) and $NEt_3$ (2.0 equiv) in $CH_3CN$ (0.1 M) and stirred overnight at room temperature. Upon completion of the reaction, the solvent was removed under reduced pressure. The residue was then dissolved in EtOAc and washed with water and brine. The extracted organic layer was dried over anhydrous $Na_2SO_4$ and concentrated under low pressure. The resultant residue was purified by MPLC.

Method 5C: A solution of Boc-protected substrate in 2.5 N HCl in ethanol (1.5 mL) was stirred at room temperature or for compounds 4 h. The reaction mixture was neutralized with $NaHCO_3$ and extracted with $CH_2Cl_2$. The organic phase was washed with water, dried over anhydrous $Na_2SO_4$, and evaporated under reduced pressure. The residue was purified by MPLC (1–10% MeOH/$CH_2Cl_2$).

The general procedures of the synthesis of the final compounds **5a–5e** are reported in our previous reported study.[3] The synthesis and characterization of the remaining final compounds are described below.

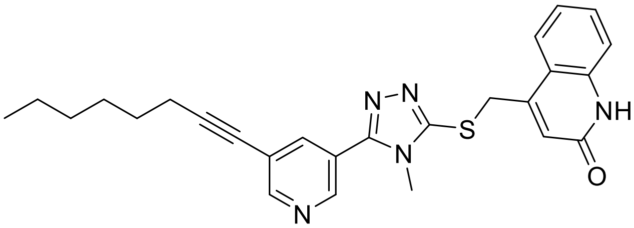

**4-(((4-Methyl-5-(5-(oct-1-yn-1-yl)pyridin-3-yl)-4H-1,2,4-triazol-3-yl)thio)methyl)quinolin-2(1H)-one** (**5f**). The title compound was synthesized according to method 5A to obtain an off-white solid (yield = 58%). $^{1}$H NMR (400 MHz, DMSO) δ 11.75 (s, 1H), 8.80 (d, $J$ = 2.2 Hz, 1H), 8.73 (d, $J$ = 2.1 Hz, 1H), 8.10 (t, $J$ = 2.1 Hz, 1H), 7.88 (dd, $J$ = 8.2, 1.5 Hz, 1H), 7.53 (ddd, $J$ = 8.4, 7.1, 1.3 Hz, 1H), 7.34 (dd, $J$ = 8.3, 1.2 Hz, 1H), 7.23 (ddd, $J$ = 8.3, 7.2, 1.3 Hz, 1H), 6.47 (s, 1H), 4.61 (s, 2H), 3.56 (s, 3H), 2.49 (t, $J$ = 7.6 Hz, 2H), 1.64–1.52 (m, 2H), 1.49–1.37 (m, 2H), 1.30 (ddt, $J$ = 7.2, 4.2, 2.6 Hz, 4H), 0.92–0.83 (m, 2H). $^{13}$C NMR (101 MHz, DMSO) δ 161.2, 152.7, 152.6, 150.3, 147.2, 146.0, 139.1, 137.6, 130.5, 124.7, 123.2, 121.9, 121.8, 120.3, 117.6, 115.8, 95.5, 76.8, 33.6, 31.7, 30.7, 28.0, 27.9, 22.0, 18.7, 13.9. HRMS (ESI): *m/z* $[M + H]^+$ Calcd for $[C_{26}H_{27}ClN_5OS + H]^+$ 458.2014, found 458.2025.

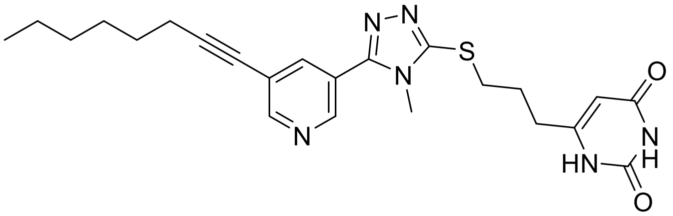

**1-(3-((4-Methyl-5-(5-(oct-1-yn-1-yl)pyridin-3-yl)-4H-1,2,4-triazol-3-yl)thio)propyl)pyrimidine-2,4(1H,3H)-dione (5g).** The title compound was synthesized according to method 5A to obtain a white solid (yield = 64%). $^1$H NMR (400 MHz, $CDCl_3$) δ 8.78 (d, *J* = 17.0 Hz, 2H), 8.59 (brs, 1H), 8.04 (t, *J* = 2.0 Hz, 1H), 7.35 (d, *J* = 7.9 Hz, 1H), 5.74 (dd, *J* = 8.0, 1.4 Hz, 1H), 3.98 (dd, *J* = 7.7, 6.5 Hz, 2H), 3.68 (s, 3H), 3.36 (t, *J* = 6.9 Hz, 2H), 2.47 (d, *J* = 14.2 Hz, 2H), 2.33–2.25 (m, 3H), 1.68–1.59 (m, 2H), 1.52–1.40 (m, 2H), 1.35 (ddd, *J* = 7.6, 5.2, 3.2 Hz, 4H), 0.98–0.87 (m, 3H). $^{13}$C NMR (101 MHz, CDCl3) δ 164.11, 153.27, 152.95, 152.11, 151.27, 146.68, 144.63, 138.17, 122.87, 121.71, 102.51, 96.07, 76.44, 47.33, 31.83, 31.30, 29.78, 28.82, 28.58, 28.38, 22.53, 19.48, 14.07. HRMS (ESI): *m/z* $[M + H]^+$ Calcd for $[C_{23}H_{28}N_6O_2S + H]^+$ 453.2073, found 453.2076.

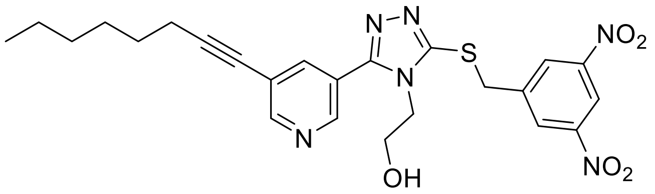

**2-(3-((3,5-Dinitrobenzyl)thio)-5-(5-(oct-1-yn-1-yl)pyridin-3-yl)-4H-1,2,4-triazol-4-yl)ethan-1-ol (8a).** The title compound was synthesized according to method 5B to obtain a white solid (yield = 74%). $^1$H NMR (400 MHz, $CDCl_3$) δ 8.9 (t, *J* = 2.1 Hz, 1H), 8.8–8.8 (m, 1H), 8.7–8.6 (m, 3H), 8.1 (t, *J* = 2.1 Hz, 1H), 7.3 (s, 2H), 4.6 (s, 2H), 4.1 (t, *J* = 5.1 Hz, 2H), 4.0 (t, *J* = 5.1 Hz, 2H), 2.4 (t, *J* = 7.2 Hz, 2H), 1.6 (p, *J* = 7.0 Hz, 4H), 1.4 (p, *J* = 7.2 Hz, 3H), 1.4–1.2 (m, 4H), 0.9–0.8 (m, 3H). $^{13}$C NMR (126 MHz, $CDCl_3$) δ 153.7, 153.5, 151.1, 148.6, 147.3, 141.5, 139.0, 129.5, 122.6, 121.7, 118.3, 96.4, 76.4, 60.3, 53.6, 47.6, 36.0, 31.4, 28.7, 28.5, 22.6, 19.6, 14.2. HRMS (ESI): *m/z* $[M + H]^+$ Calcd for $[C_{24}H_{26}N_6O_5S + H]^+$ 511.1764, found 511.1793.

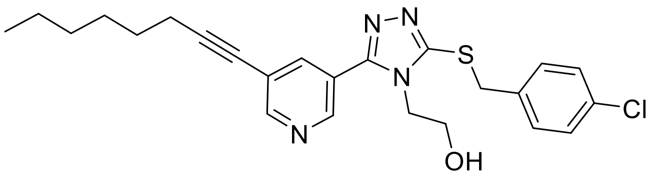

**2-(3-((4-Chlorobenzyl)thio)-5-(5-(oct-1-yn-1-yl)pyridin-3-yl)-4H-1,2,4-triazol-4-yl)ethan-1-ol (8b).** The title compound was synthesized according to method 5B to obtain a white solid (yield = 43%). $^{1}$H NMR (500 MHz, $CDCl_3$) δ 8.80 (s, 1H), 8.66 (s, 1H), 7.23 (dt, *J* = 17.2, 9.1 Hz, 4H), 4.29 (s, 2H), 3.92 (dt, *J* = 29.1, 4.7 Hz, 4H), 2.43 (t, *J* = 7.2 Hz, 2H), 1.61 (p, *J* = 7.2 Hz, 2H), 1.46 (q, *J* = 7.7 Hz, 2H), 1.39–1.22 (m, 4H), 0.91 (t, *J* = 6.9 Hz, 3H). $^{13}$C NMR (126 MHz, $CDCl_3$) δ 153.5, 153.5, 151.9, 147.7, 139.1, 134.9, 133.9, 130.5, 129.0, 123.0, 121.5, 96.0, 60.6, 47.4, 37.8, 31.5, 28.8, 28.6, 22.7, 19.7, 14.2. HRMS (ESI): *m/z* $[M + H]^+$ Calcd for $[C_{24}H_{27}ClN_4OS + H]^+$ 455.1672, found 455.1679.

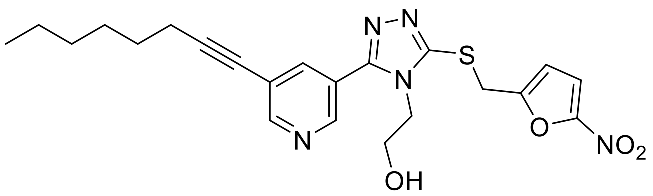

**2-(3-(((5-Nitrofuran-2-yl)methyl)thio)-5-(5-(oct-1-yn-1-yl)pyridin-3-yl)-4H-1,2,4-triazol-4-yl)ethan-1-ol (8e).** The title compound was synthesized according to method 5B to obtain a white solid (yield = 47%). $^{1}$H NMR (400 MHz, $CDCl_3$) δ 8.9–8.5 (m, 2H), 8.1 (d, *J* = 2.0 Hz, 1H), 7.2 (d, *J* = 3.7 Hz, 1H), 6.6 (d, *J* = 3.7 Hz, 1H), 4.4 (s, 2H), 4.1–4.0 (m, 2H), 4.0–3.9 (m, 2H), 2.4 (t, *J* = 7.2 Hz, 2H), 1.6 (p, *J* = 7.1 Hz, 2H), 1.5–1.4 (m, 2H), 1.3 (tt, *J* = 7.6, 3.5 Hz, 4H), 0.9–0.8 (m, 3H). $^{13}$C NMR (101 MHz, $CDCl_3$) δ 153.8, 153.6, 153.5, 151.8, 151.0, 147.4, 139.1, 122.7, 121.7 112.8, 112.7, 96.2, 76.5, 60.5, 47.6, 31.4, 29.9, 28.8, 28.5, 22.7, 19.6, 14.2.

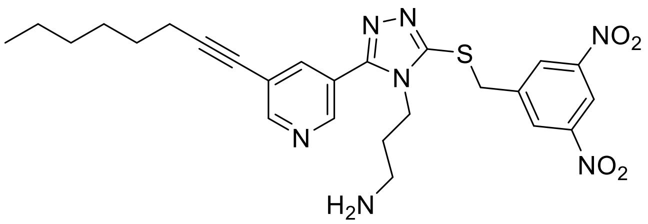

**3-(3-((3,5-Dinitrobenzyl)thio)-5-(5-(oct-1-yn-1-yl)pyridin-3-yl)-4H-1,2,4-triazol-4-yl)propan-1-amine (12a).** The title compound was synthesized according to method 5B followed by method 5C to obtain a brown sticky solid (yield = 40%). $^1$H NMR (400 MHz, Methanol-$d_4$) δ 8.99 (s, 2H), 8.92 (s, 1H), 8.80 (s, 2H), 8.50 (s, 1H), 4.82 (s, 2H), 4.28 (s, 2H), 2.95 (s, 2H), 2.54 (t, $J$ = 6.8 Hz, 2H), 2.07 (s, 2H), 1.66 (s, 2H), 1.50 (s, 2H), 1.36 (dd, $J$ = 7.2, 3.8 Hz, 4H), 0.97–0.89 (m, 3H). $^{13}$C NMR (126 MHz, Methanol-$d_4$) δ 154.5, 154.3, 152.4, 149.9, 148.0, 143.5, 140.0, 130.5, 124.3, 123.4, 121.3, 118.9, 97.4, 77.1, 43.4, 38.1, 36.7, 32.5, 29.7, 29.5, 23.6, 20.1, 14.4. HRMS (ESI): *m/z* $[M + H]^+$ Calcd for $[C_{25}H_{29}N_7O_4S + H]^+$ 524.2080, found 524.2181.

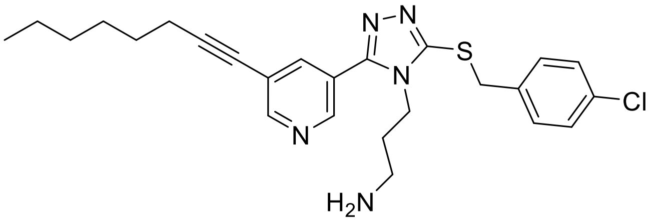

**3-(3-((4-Chlorobenzyl)thio)-5-(5-(oct-1-yn-1-yl)pyridin-3-yl)-4H-1,2,4-triazol-4-yl)propan-1-amine (12b)**. The title compound was synthesized according to method 5B followed by method 5C to obtain a brown sticky solid (yield = 30%). $^1$H NMR (400 MHz, $CDCl_3$) δ 8.68 (dd, $J$ = 13.1, 2.1 Hz, 1H), 7.95 (t, $J$ = 2.1 Hz, 0H), 7.36–7.23 (m, 1H), 4.02–3.88 (m, 1H), 2.68 (t, $J$ = 6.8 Hz, 1H), 2.43 (t, $J$ = 7.1 Hz, 1H), 1.77 (p, $J$ = 6.7 Hz, 1H), 1.61 (p, $J$ = 7.1 Hz, 1H), 1.51–1.23 (m, 2H), 0.91 (q, $J$ = 5.0, 3.1 Hz, 1H). $^{13}$C NMR (101 MHz, $CDCl_3$) δ 153.4, 152.6, 151.7, 146.5, 138.8, 135.3, 133.9, 130.7, 129.0, 123.4, 121.9, 96.3, 76.5, 42.6, 38.3, 37.4, 32.1, 31.4, 28.7, 28.5, 22.7, 19.6, 14.2. HRMS (ESI): *m/z* $[M + H]^+$ Calcd for $[C_{25}H_{30}ClN_5S + H]^+$ 468.1989, found 468.1988.

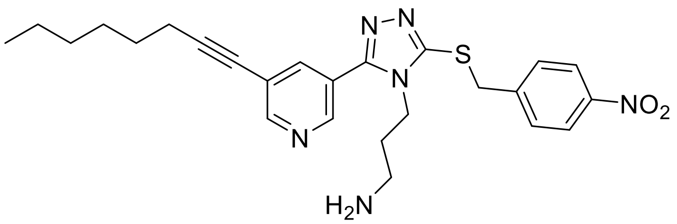

**3-(3-((4-Nitrobenzyl)thio)-5-(5-(oct-1-yn-1-yl)pyridin-3-yl)-4H-1,2,4-triazol-4-yl)propan-1-amine (12c).** The title compound was synthesized according to method 5B followed by method 5C to obtain a brown sticky solid (yield = 40%). $^1$H NMR (500 MHz, $CDCl_3$) δ 8.7–8.7 (m, 2H), 8.2 (d, $J$ = 8.7 Hz, 2H), 8.0 (t, $J$ = 2.1 Hz, 1H), 7.6 (d, $J$ = 8.5 Hz, 2H), 4.6 (s, 2H), 4.0 (t, $J$ = 7.6 Hz, 2H), 2.6 (t, $J$ = 6.6 Hz, 2H), 2.4 (t, $J$ = 7.1 Hz, 2H), 1.7 (p, $J$ = 6.7 Hz, 2H), 1.6 (h, $J$ = 6.4, 5.8 Hz, 2H), 1.4 (td, $J$ = 12.4, 10.4, 5.3 Hz, 3H), 1.4–1.3 (m, $J$ = 4.4, 3.8 Hz, 5H), 0.9 (t, $J$ = 6.9 Hz, 3H). $^{13}$C NMR (126 MHz, $CDCl_3$) δ 153.5, 152.9, 151.2, 147.6, 146.6, 144.5, 138.6, 130.3, 124.0, 123.2, 121.9, 96.3, 76.5, 42.7, 38.5, 36.4, 32.9, 31.4, 29.8, 28.7, 28.5, 22.8, 22.7, 19.6, 14.2. HRMS (ESI): *m/z* $[M + H]^+$ Calcd for $[C_{25}H_{30}N_6O_2S + H]^+$ 479.2229, found 479.2220.

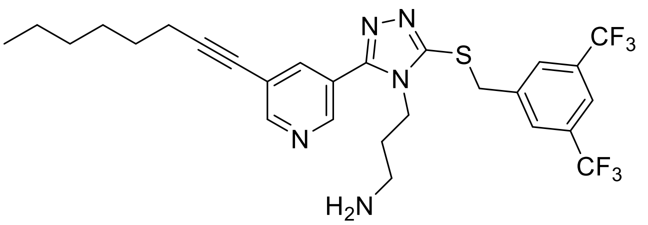

**3-(3-((3,5-bis(Trifluoromethyl)benzyl)thio)-5-(5-(oct-1-yn-1-yl)pyridin-3-yl)-4H-1,2,4-triazol-4-yl)propan-1-amine (12d).** The title compound was synthesized according to method 5B followed by method 5C to a obtain brown solid (yield = 50%). $^1$H NMR (500 MHz, $CDCl_3$) δ 8.69 (d, $J$ = 2.1 Hz, 1H), 8.64 (d, $J$ = 2.0 Hz, 1H), 7.94 (d, $J$ = 1.8 Hz, 1H), 7.90 (s, 2H), 7.77 (s, 1H), 4.61 (s, 2H), 3.99 (t, $J$ = 7.7 Hz, 2H), 2.71 (t, $J$ = 6.9 Hz, 2H), 2.41 (t, $J$ = 7.2 Hz, 2H), 1.80 (p, $J$

= 6.9 Hz, 2H), 1.59 (p, $J$ = 7.2 Hz, 2H), 1.42 (p, $J$ = 7.2 Hz, 2H), 1.36–1.24 (m, $J$ = 4.6, 3.8 Hz, 5H), 0.88 (t, $J$ = 6.9 Hz, 3H). $^{13}$C NMR (126 MHz, $CDCl_3$) δ 153.4, 152.9, 151.0, 146.4, 139.7, 138.8, 132.1 (q, $J$ = 33.5 Hz), 129.5 (d, $J$ = 4.0 Hz), 123.2 (q, $J$ = 272.8 Hz), 123.2, 122.0, 121.9 (p, $J$ = 3.5 Hz), 96.4, 76.4, 38.2, 36.4, 31.8, 31.4, 28.7, 28.5, 19.6, 14.1.

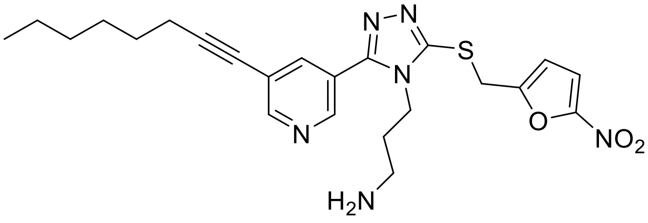

**3-(3-(((5-Nitrofuran-2-yl)methyl)thio)-5-(5-(oct-1-yn-1-yl)pyridin-3-yl)-4H-1,2,4-triazol-4-yl)propan-1-amine (12e).** The title compound was synthesized according to method 5B followed by method 5C to obtain a brown solid (yield = 96%). $^{1}$H NMR (400 MHz, MeOD) δ 8.79–8.72 (m, 2H), 8.13 (q, $J$ = 2.6, 2.0 Hz, 1H), 7.41 (d, $J$ = 3.8 Hz, 1H), 6.73 (d, $J$ = 3.7 Hz, 1H), 4.63 (s, 2H), 4.23 (t, $J$ = 7.7 Hz, 2H), 2.92–2.84 (m, 2H), 2.51 (t, $J$ = 7.1 Hz, 2H), 2.10–1.88 (m, 2H), 1.66 (p, $J$ = 7.5, 6.9 Hz, 2H), 1.57–1.46 (m, 2H), 1.43–1.31 (m, 4H), 1.01–0.90 (m, 3H). $^{13}$C NMR (101 MHz, MeOD) δ 155.5, 154.6, 154.3, 152.1, 148.1, 140.1, 124.4, 123.5, 113.8 (d, $J$ = 1.8 Hz), 97.4, 77.1, 49.6, 49.4, 49.2, 49.0, 48.6, 48.3, 43.3, 42.2, 37.7, 32.5, 32.5, 31.1, 29.7, 29.5, 29.0, 23.6, 20.1, 20.1, 14.4. HRMS (ESI): $m/z$ [M + Cs]$^+$ Calcd for $[C_{23}H_{28}N_6O_3S + Cs]^+$ 603.1155, found 603.1179.

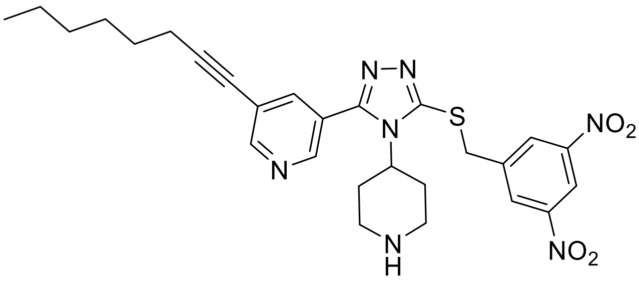

**3-(5-((3,5-Dinitrobenzyl)thio)-4-(piperidin-4-yl)-4H-1,2,4-triazol-3-yl)-5-(oct-1-yn-1-yl)pyridine (13).** The title compound was synthesized according to method 5B followed by method 5C to obtain a brown solid (yield = 42%). $^1$H NMR (500 MHz, $CDCl_3$) δ 8.9 (d, *J* = 2.3 Hz, 1H), 8.7 (dd, *J* = 6.5, 2.1 Hz, 3H), 8.5 (d, *J* = 2.1 Hz, 1H), 7.8 (d, *J* = 2.3 Hz, 1H), 4.8 (s, 2H), 4.1 (tt, *J* = 12.5, 4.3 Hz, 1H), 3.3–3.2 (m, 2H), 2.7–2.6 (m, 2H), 2.5–2.3 (m, 4H), 1.8 (dd, *J* = 13.0, 3.6 Hz, 2H), 1.6 (p, *J* = 7.2 Hz, 2H), 1.4 (p, *J* = 7.2 Hz, 2H), 1.4–1.2 (m, 4H), 0.9 (t, *J* = 6.9 Hz, 3H). $^{13}$C NMR (126 MHz, $CDCl_3$) δ 153.8, 153.5, 149.2, 148.6, 147.2, 142.0, 139.2, 129.6, 123.2, 122.0, 118.2, 96.5, 76.4, 55.6, 45.7, 35.7, 31.4, 31.3, 28.7, 28.5, 22.6, 19.6, 14.2. HRMS (ESI): *m/z* $[M - H]^-$ Calcd for $[C_{27}H_{31}N_7O_4S - H]^-$ 548.2080, found 548.2094.

**7. Synthesis of final analog 17**

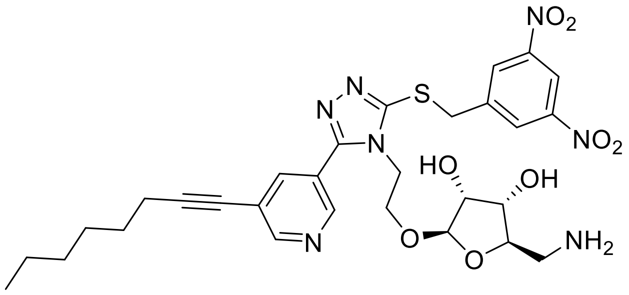

**2-(Aminomethyl)-5-(2-(3-((3,5-dinitrobenzyl)thio)-5-(5-(oct-1-yn-1-yl)pyridin-3-yl)-4H-1,2,4-triazol-4-yl)ethoxy)tetrahydrofuran-3,4-diol (17).** To *tert*-butyl ((6-(2-(3-((3,5-dinitrobenzyl)thio)-5-(5-(oct-1-yn-1-yl)pyridin-3-yl)-4H-1,2,4-triazol-4-yl)ethoxy)-2,2-diethyltetrahydrofuro[3,4-d][1,3]dioxol-4-yl)methyl)carbamate (18 mg, 22 μmol) in a 1 gram vial was added 1 mL of 80% TFA in dichloromethane and stirred for 2 h. The solvent was removed under reduced pressure and the residue purified by PTLC 15% MeOH/$CH_2Cl_2$. Compound **17** was obtained in 40% yield. $^1$H NMR (400 MHz, MeOD) δ 8.90 (t, *J* = 2.2 Hz, 1H), 8.83–8.66 (m, 4H), 8.13 (s, 1H), 4.76–4.64 (m, 3H), 4.24 (t, *J* = 5.4 Hz, 2H), 4.01–3.87 (m, 2H), 3.85–3.75 (m, 1H),

3.73 (d, *J* = 4.5 Hz, 1H), 3.68–3.59 (m, 1H), 3.16–3.09 (m, 1H), 2.61 (dd, *J* = 13.0, 9.8 Hz, 1H), 2.49 (t, *J* = 7.1 Hz, 2H), (p, *J* = 8.2, 7.6, 6.9 Hz, 2H)1.55–1.43 (m, 2H), 1.41–1.29 (m, 4H), 0.96–0.88 (m, 3H). $^{13}$C NMR (101 MHz, MeOD) δ 154.9, 154.2, 152.9, 149.9, 148.3, 143.5, 140.3, 130.4, 124.6, 123.2, 118.9, 109.0, 97.4, 80.2, 77.0, 75.7, 74.2, 66.7, 46.1, 45.2, 37.0, 32.5, 29.7, 29.5, 23.6, 20.1, 14.4. HRMS (ESI): *m/z* $[M + H]^+$ Calcd for $[C_{29}H_{35}N_7O_8S + H]^+$ 642.2346, found 642.2350.

**8. Synthesis of final analog 18–23**

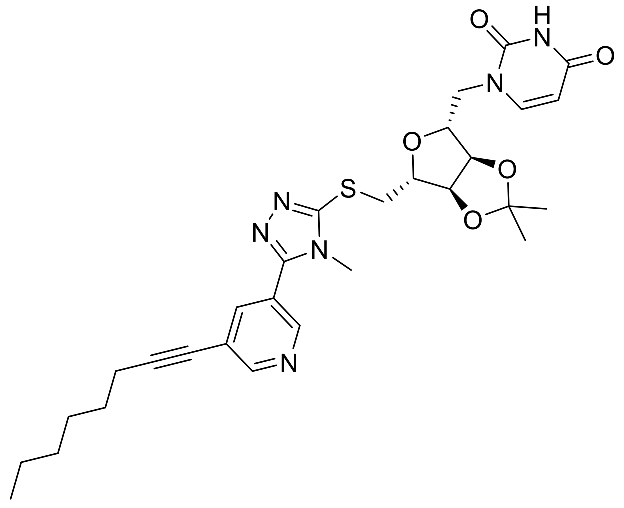

**1-((3aR,4R,6S,6aS)-2,2-dimethyl-6-(((4-methyl-5-(5-(oct-1-yn-1-yl)pyridin-3-yl)-4H-1,2,4-triazol-3-yl)thio)methyl)tetrahydrofuro[3,4-d][1,3]dioxol-4-yl)pyrimidine-2,4(1H,3H)-dione (18)** To 1-((3aR,4R,6S,6aS)-6-(bromomethyl)-2,2-dimethyltetrahydrofuro[3,4-d][1,3]dioxol-4-yl)pyrimidine-2,4(1H,3H)-dione (98 mg, 1.3 equiv, 0.28 mmol), 4-methyl-5-(5-(oct-1-yn-1-yl)pyridin-3-yl)-4H-1,2,4-triazole-3-thiol (65 mg, 1 equiv, 0.22 mmol) and $K_2CO_3$ (45 mg, 1.5 equiv, 0.32 mmol) was added acetone (1.5 mL) and MeOH (0.5 mL) and stirred overnight. The solvent was removed under low pressure and the residue dissolved with EtOAc, and washed with water (x2), brine. The extracted organic layer was dried over anhydrous $Na_2SO_4$, and then concentrated under low pressure. Then reaction mixture was purified by MPLC (gradient 5% MeOH DCM). Compound **18** was obtained in 37% yield. $^{1}$H NMR (400 MHz, MeOD) δ 8.77 (d, *J* = 2.1 Hz, 1H), 8.70 (d, *J* = 2.0 Hz, 1H), 8.11 (t, *J* = 2.0 Hz, 1H), 7.58 (d, *J* = 8.0 Hz, 1H), 5.67 (d, *J* = 1.9 Hz, 1H), 5.64 (d, *J* = 7.9 Hz, 1H), 5.13 (dd, *J* = 6.5, 1.9 Hz, 1H), 4.40–4.29 (m, 1H), 3.74 (s, 3H), 3.59 (dd, *J* = 13.7, 7.4 Hz, 1H), 3.50 (dd, *J* = 13.7, 6.2 Hz, 1H), 2.50 (t, *J* = 7.1 Hz, 2H), 1.70–1.60 (m, 2H), 1.56–1.45 (m, 5H), 1.42–1.29 (m, 7H), 0.96–0.90 (m, 3H). $^{13}$C NMR (126 MHz, MeOD) δ 166.1, 154.3, 154.0, 153.9, 151.8, 147.9, 145.2, 139.7, 124.5, 123.2, 115.4, 102.9, 97.1, 96.5, 88.2, 85.7, 85.0, 77.2, 37.0, 32.8, 32.5, 29.7, 29.5, 27.4, 25.4, 23.6, 20.1, 14.4. HRMS (ESI): *m/z* $[M + H]^+$ Calcd for $[C_{23}H_{28}N_6O_2S + H]^+$ 453.2073, found 453.2085.

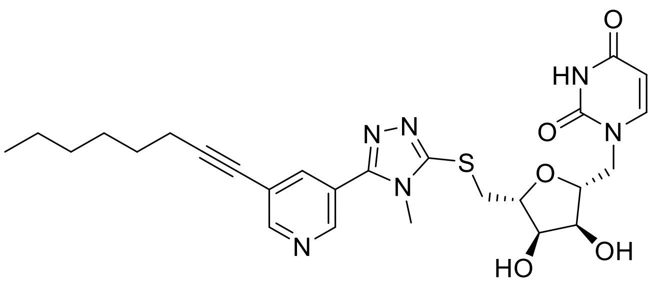

**1-((2R,3R,4S,5S)-3,4-Dihydroxy-5-(((4-methyl-5-(5-(oct-1-yn-1-yl)pyridin-3-yl)-4H-1,2,4-triazol-3-yl)thio)methyl)tetrahydrofuran-2-yl)pyrimidine-2,4(1H,3H)-dione (21).** 1-((3aR,4R,6S,6aS)-2,2-dimethyl-6-(((4-methyl-5-(5-(oct-1-yn-1-yl)pyridin-3-yl)-4H-1,2,4-triazol-3-yl)thio)methyl)tetrahydrofuro[3,4-d][1,3]dioxol-4-yl)pyrimidine-2,4(1H,3H)-dione (30 mg, 1 equiv, 53 µmol) was added 80% TFA in water (1 mL) and stirred at room temperature for 4 h. The solvent was evaporated under reduced pressure and the residue triturated with diethyl ether. Compound **21** was obtained in 43% yield. $^{1}$H NMR (400 MHz, MeOD) δ 8.77 (d, *J* = 2.1 Hz, 1H), 8.72 (d, *J* = 2.0 Hz, 1H), 8.12 (t, *J* = 2.1 Hz, 1H), 7.60 (d, *J* = 8.2 Hz, 1H), 5.80 (d, *J* = 4.6 Hz, 1H), 5.72 (d, *J* = 8.1 Hz, 1H), 4.33–4.26 (m, 1H), 4.18 (dt, *J* = 7.6, 4.6 Hz, 1H), 4.12 (t, *J* = 5.4 Hz, 1H), 3.75 (s, 3H), 3.62 (dd, *J* = 13.9, 4.5 Hz, 1H), 3.55 (dd, *J* = 13.9, 7.7 Hz, 1H), 2.51 (t, *J* =

7.0 Hz, 2H), 1.66 (p, *J* = 8.0, 7.5, 7.0 Hz, 2H), 1.56–1.47 (m, 2H), 1.43–1.34 (m, 4H), 0.99–0.91 (m, 3H). $^{13}$C NMR (126 MHz, MeOD) δ 165.9, 154.4, 154.2, 154.1, 152.2, 147.9, 142.9, 139.7, 124.5, 123.2, 103.1, 97.1, 92.1, 84.0, 77.2, 74.5, 73.9, 37.2, 32.7, 32.5, 29.7, 29.5, 23.6, 20.1, 14.4. HRMS (ESI): *m/z* $[M + H]^+$ Calcd for $[C_{25}H_{30}N_6O_5S + H]^+$ 527.2076, found 572.2065.

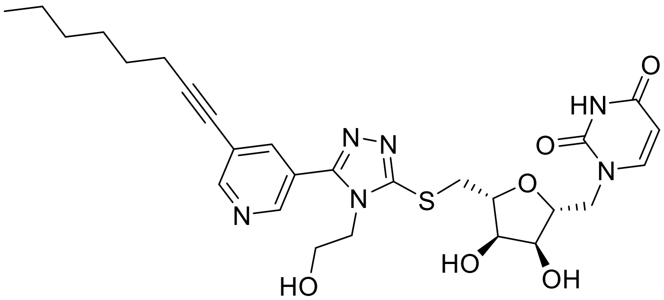

**1-((2R,3R,4S,5S)-3,4-dihydroxy-5-(((4-(2-hydroxyethyl)-5-(5-(oct-1-yn-1-yl)pyridin-3-yl)-4H-1,2,4-triazol-3-yl)thio)methyl)tetrahydrofuran-2-yl)pyrimidine-2,4(1H,3H)-dione (22)**

To 1-((3aR,4R,6S,6aS)-6-(((4-(2-Hydroxyethyl)-5-(5-(oct-1-yn-1-yl)pyridin-3-yl)-4H-1,2,4-triazol-3-yl)thio)methyl)-2,2-dimethyltetrahydrofuro[3,4-d][1,3]dioxol-4-yl)pyrimidine-2,4(1H,3H)-dione (10 mg, 1 equiv, 17 µmol) was added 80% TFA in water (1 mL) and stirred at room temperature for 4 h. The solvent was evaporated under reduced pressure and the residue triturated with diethyl ether. Compound **22** was obtained in _% yield. $^1$H NMR (500 MHz, MeOD) δ 8.90–8.77 (m, 2H), 8.70 (s, 1H), 8.23 (d, *J* = 1.8 Hz, 1H), 7.61 (d, *J* = 8.1 Hz, 1H), 5.79 (d, *J* = 4.6 Hz, 1H), 5.72 (d, *J* = 8.1 Hz, 1H), 4.29 (t, *J* = 5.1 Hz, 1H), 4.19 (tq, *J* = 7.6, 4.9, 3.7 Hz, 3H), 4.12 (t, *J* = 5.4 Hz, 1H), 3.79 (t, *J* = 5.3 Hz, 2H), 3.67 (dd, *J* = 13.9, 4.7 Hz, 1H), 3.59 (dd, *J* = 14.0, 7.7 Hz, 1H), 2.48 (t, *J* = 7.1 Hz, 2H), 1.64 (p, *J* = 7.1 Hz, 2H), 1.54–1.44 (m, 2H), 1.36 (dp, *J* = 7.5, 4.4, 3.7 Hz, 4H), 1.03–0.84 (m, 3H). $^{13}$C NMR (101 MHz, MeOD) δ 165.9, 154.9, 154.0, 152.2, 148.6, 142.9, 140.6, 124.9, 123.1, 103.1, 97.0, 92.1, 83.8, 77.2, 74.6, 73.9, 64.9, 60.7, 44.7,

37.2, 32.5, 29.7, 29.5, 23.6, 20.1, 14.4. HRMS (ESI): *m/z* $[M + H]^+$ Calcd for $[C_{26}H_{32}N_6O_6S + H]^+$ 557.2182, found 557.2193.

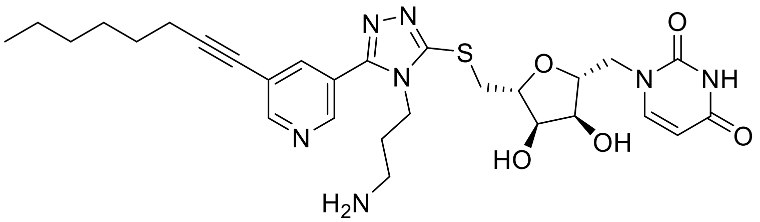

**1-((2R,3R,4S,5S)-5-(((4-(3-aminopropyl)-5-(5-(oct-1-yn-1-yl)pyridin-3-yl)-4H-1,2,4-triazol-3-yl)thio)methyl)-3,4-dihydroxytetrahydrofuran-2-yl)pyrimidine-2,4(1H,3H)-dione 2,2,2-trifluoroacetate (23).** *tert*-Butyl (3-(3-((((3aS,4S,6R,6aR)-6-(2,4-dioxo-3,4-dihydropyrimidin-1(2H)-yl)-2,2-dimethyltetrahydrofuro[3,4-d][1,3]dioxol-4-yl)methyl)thio)-5-(5-(oct-1-yn-1-yl)pyridin-3-yl)-4H-1,2,4-triazol-4-yl)propyl)carbamate (20 mg, 1 equiv, 28 μmol) was added 80% TFA in water (1 mL) and stirred at room temperature for 4 h. The solvent evaporated under reduced pressure and the residue triturated with diethyl ether. Compound **23** was obtained in 62% yield. $^1$H NMR (400 MHz, MeOD) δ 12.67 (dd, *J* = 15.5, 1.9 Hz, 2H), 12.02 (t, *J* = 2.0 Hz, 1H), 11.54 (d, *J* = 8.1 Hz, 1H), 9.73 (d, *J* = 4.6 Hz, 1H), 9.67 (d, *J* = 8.1 Hz, 1H), 9.44 (s, 2H), 8.26 (dd, *J* = 5.6, 4.6 Hz, 1H), 8.22–8.02 (m, 4H), 7.64 (dd, *J* = 13.9, 4.3 Hz, 1H), 7.55 (dd, *J* = 13.9, 8.4 Hz, 1H), 6.86–6.77 (m, 3H), 6.44 (t, *J* = 7.1 Hz, 3H), 5.94 (p, *J* = 6.9 Hz, 2H), 5.65–5.53 (m, 3H), 5.50–5.38 (m, 2H), 5.38–5.25 (m, 5H), 4.92–4.84 (m, 3H). NMR (101 MHz, MeOD) δ 165.9, 154.6, 154.5, 153.8, 152.3, 147.9, 142.9, 140.0, 124.4, 123.5, 103.2, 97.4, 92.1, 83.9, 77.1, 74.5, 74.1, 43.2, 37.8, 37.3, 32.5, 29.7, 29.5, 28.9, 23.6, 20.1, 14.4. HRMS (ESI): *m/z* $[M + H]^+$ Calcd for $[C_{27}H_{35}N_7O_5S + H]^+$ 570.2499, found 570.2469. Purity: 98.44.

#### $^{1}H$ and $^{13}C$ NMR spectrum of the final selected compounds

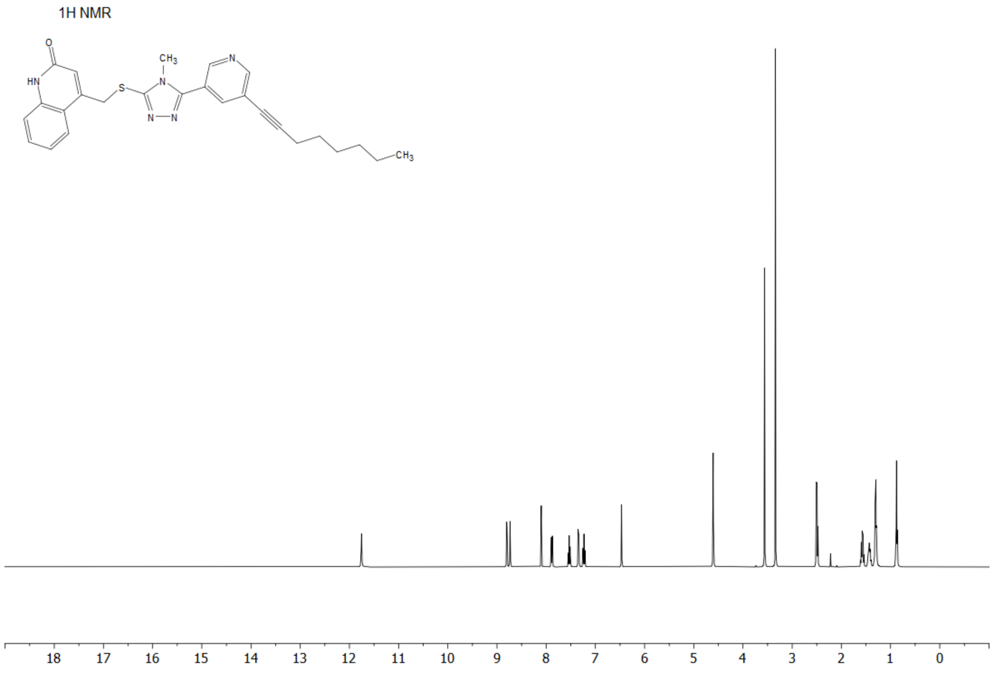

The $^{1}$H NMR spectrum of compound **5f**

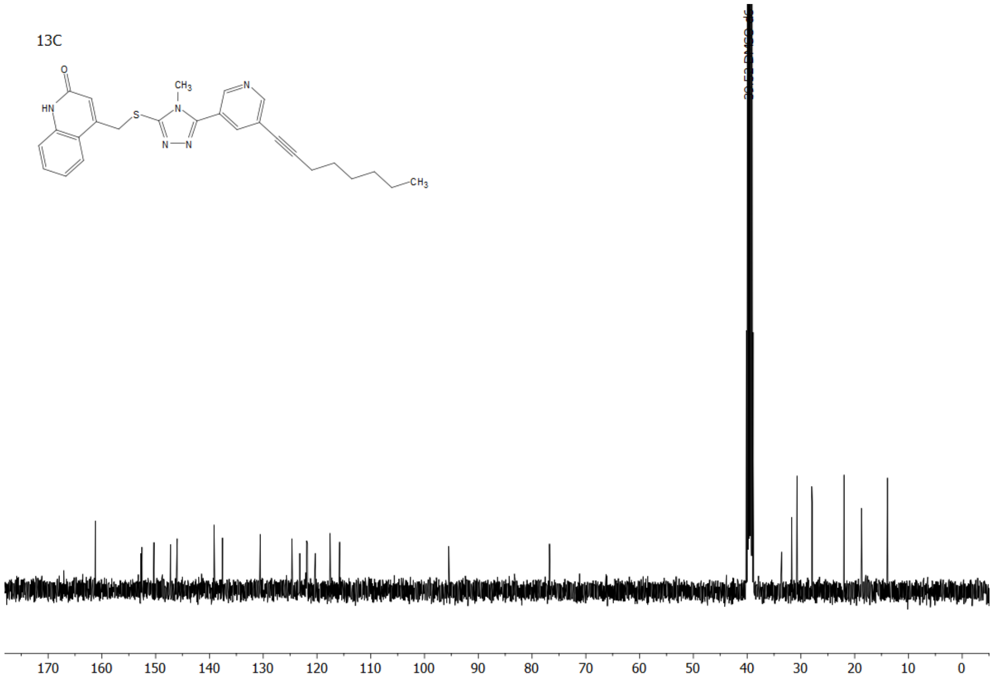

The $^{13}$C NMR spectrum of compound **5f**

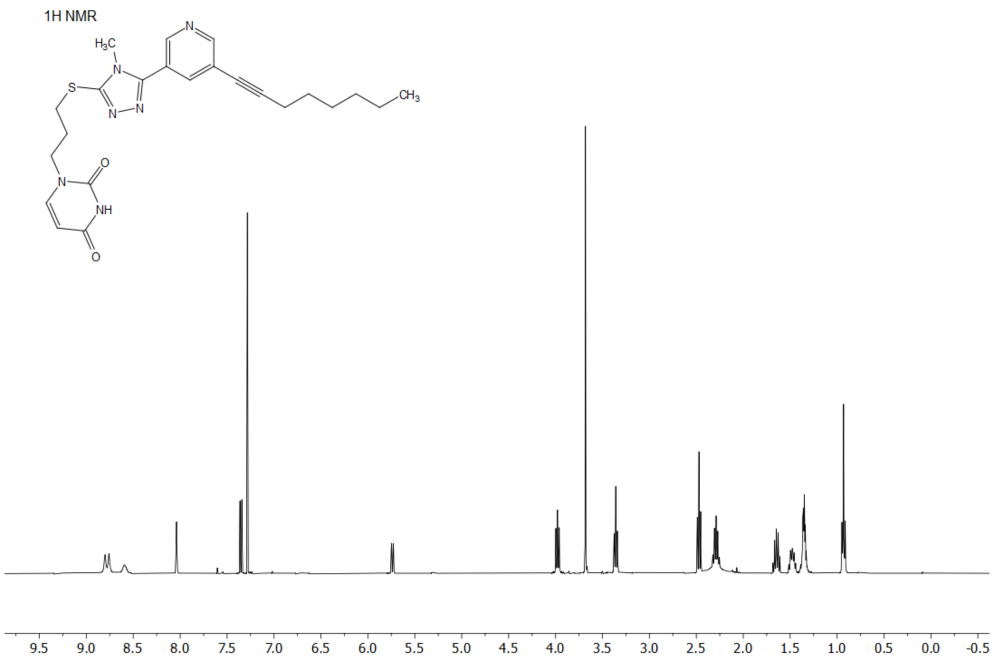

The $^{1}H$ NMR spectrum of compound **5g**

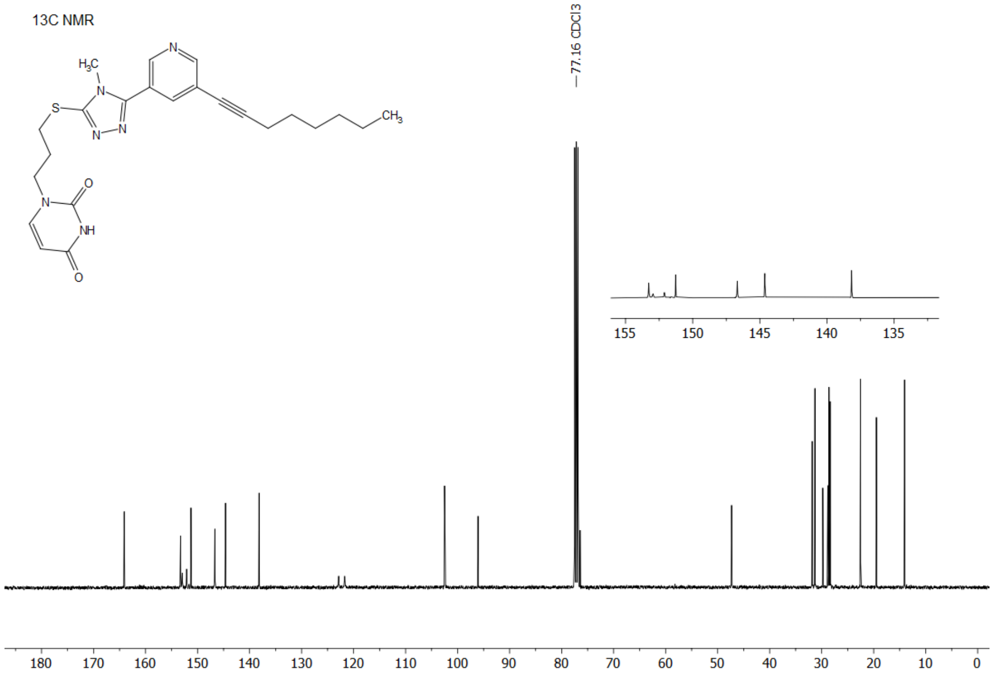

The $^{13}C$ NMR spectrum of compound **5g**

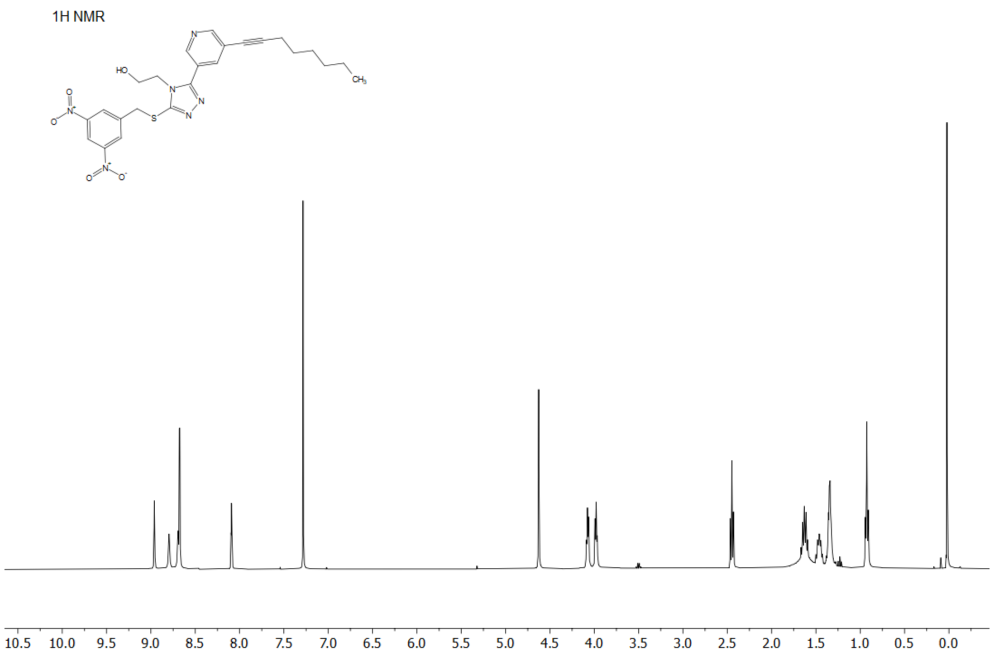

The $^{1}H$ NMR spectrum of compound **8a**

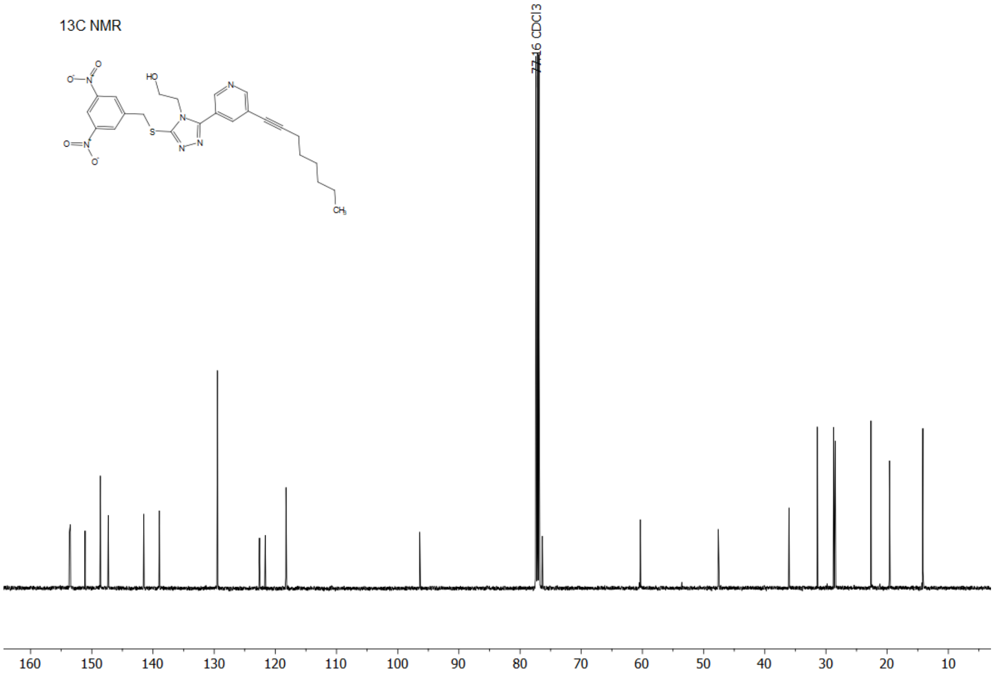

The $^{13}C$ NMR spectrum of compound **8a**

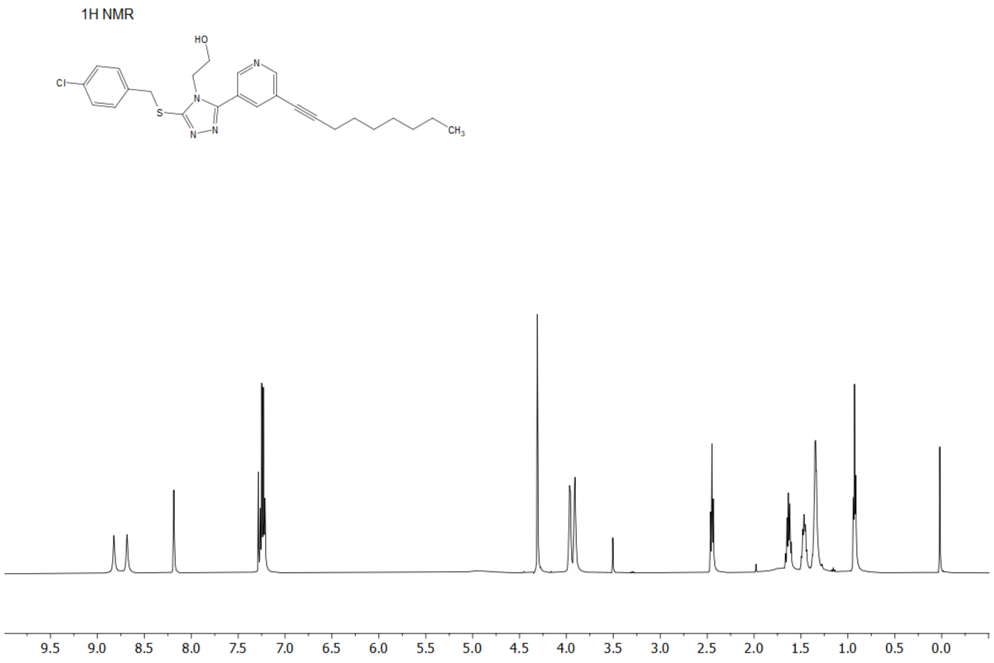

The $^{1}H$ NMR spectrum of compound **8b**

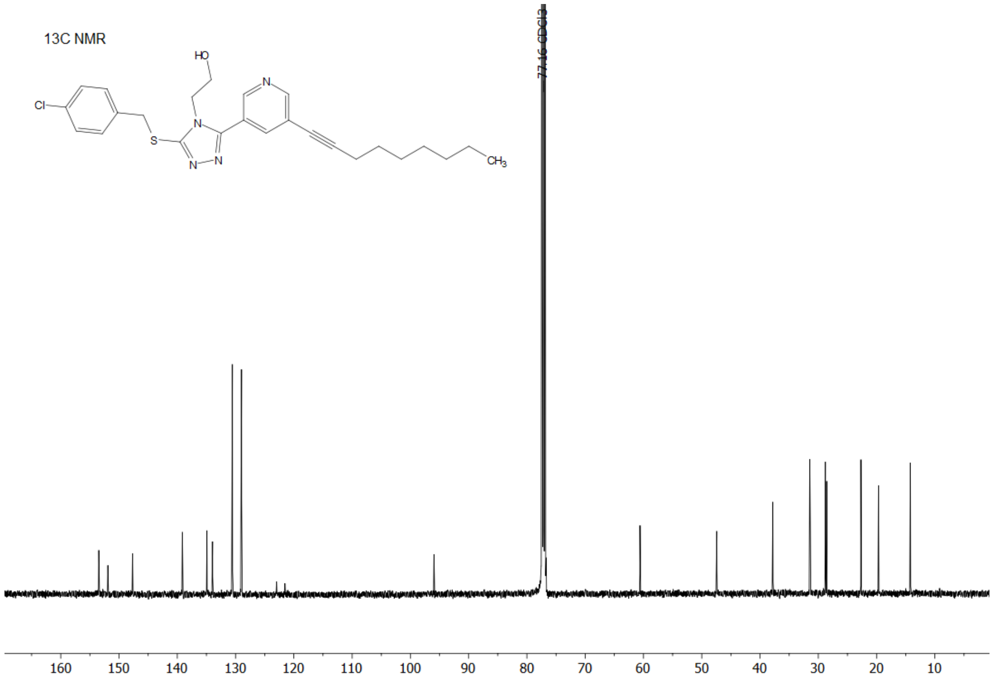

The $^{13}C$ NMR spectrum of compound **8b**

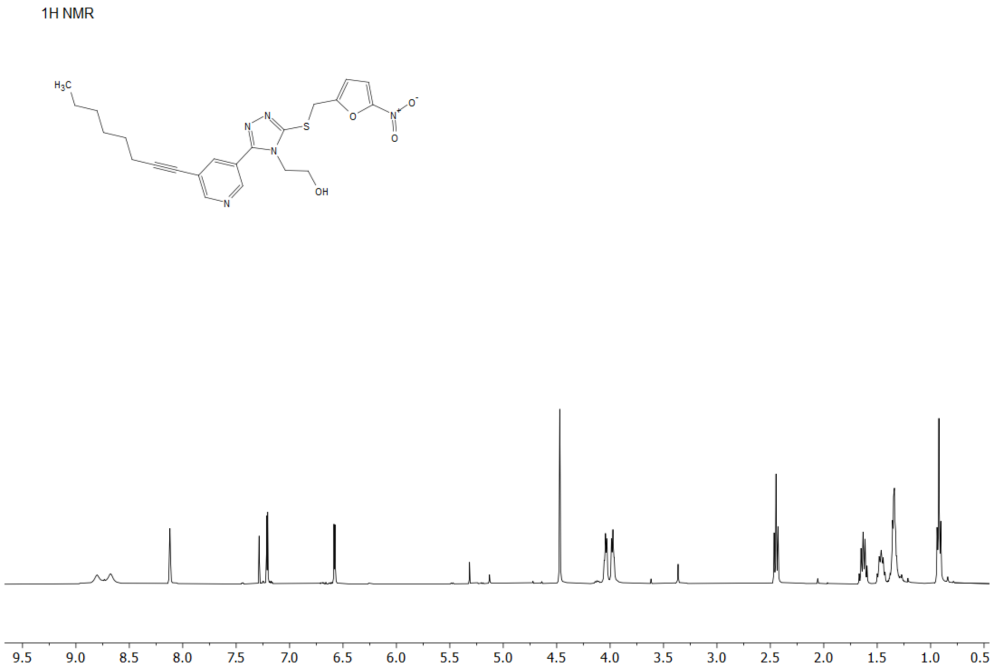

The $^{1}H$ NMR spectrum of compound **8e**

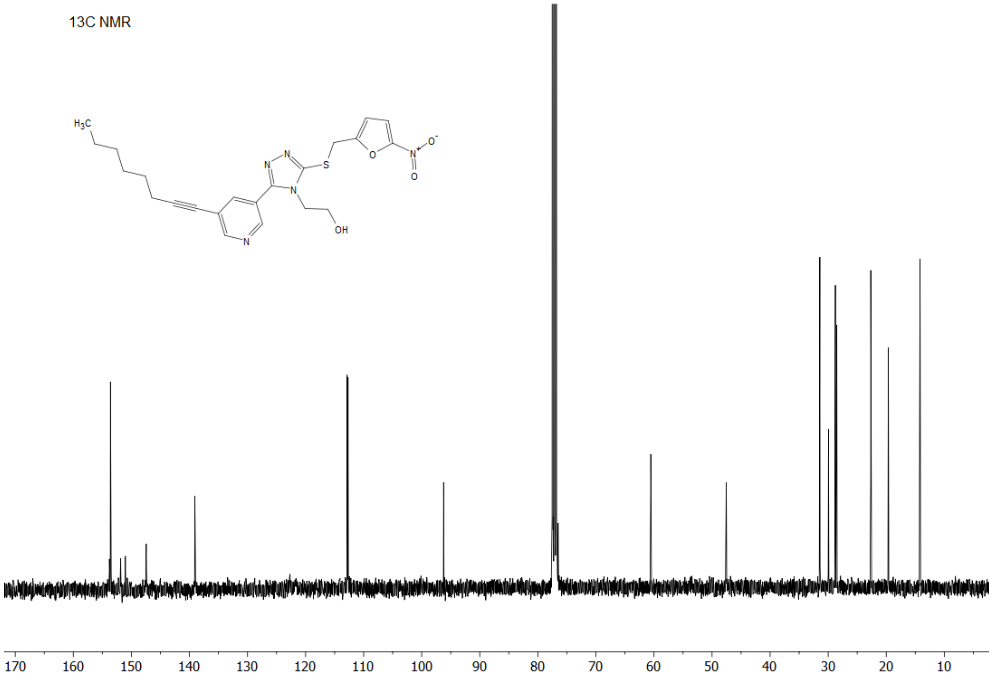

The $^{13}C$ NMR spectrum of compound **8e**

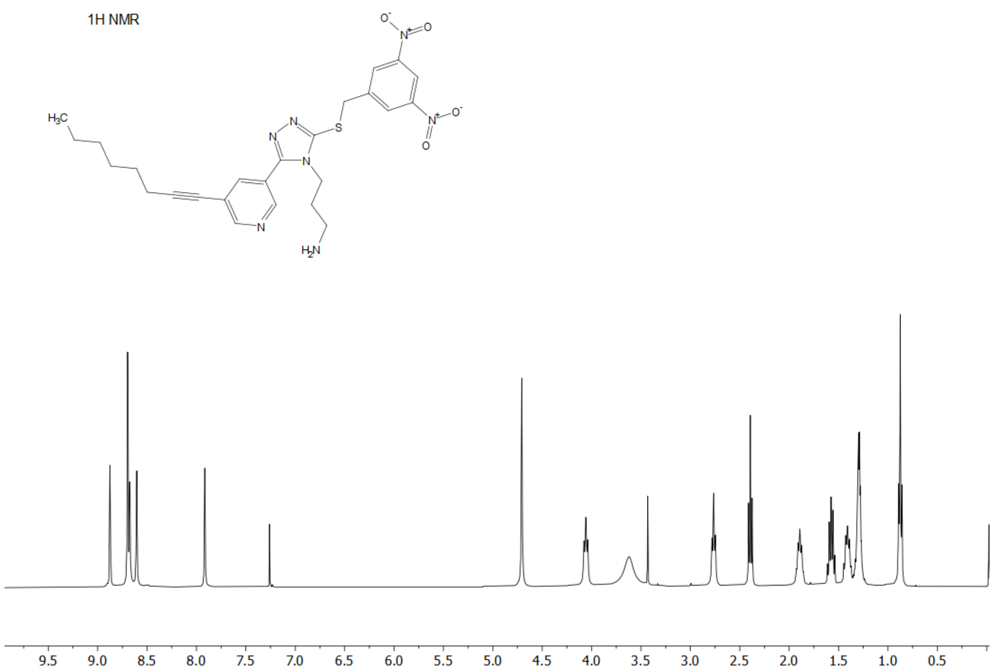

The $^{1}H$ NMR spectrum of compound **12a**

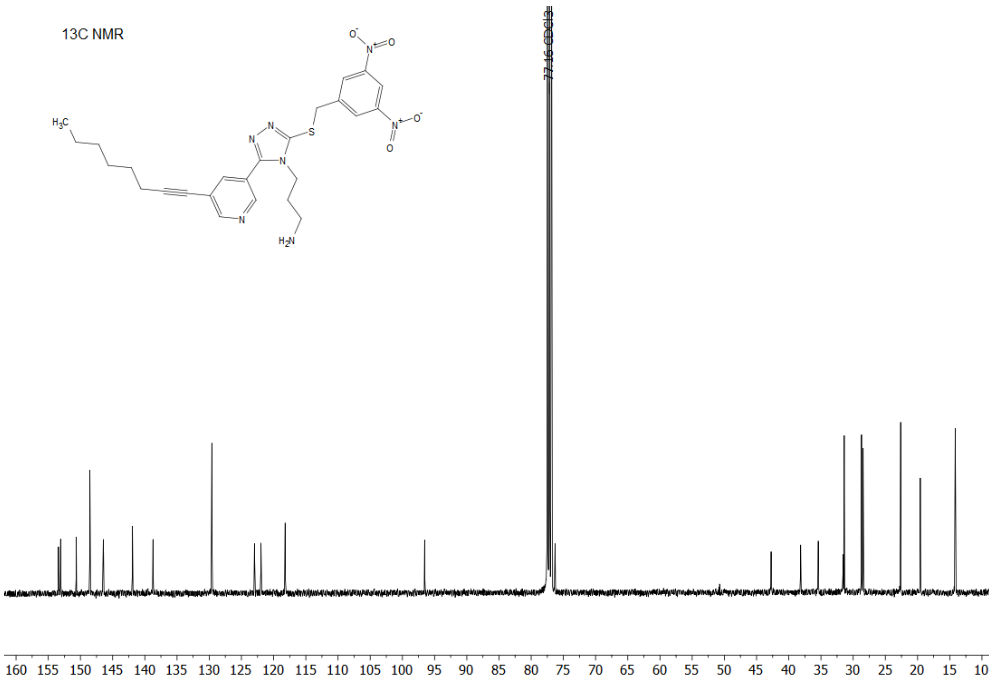

The $^{13}C$ NMR spectrum of compound **12a**

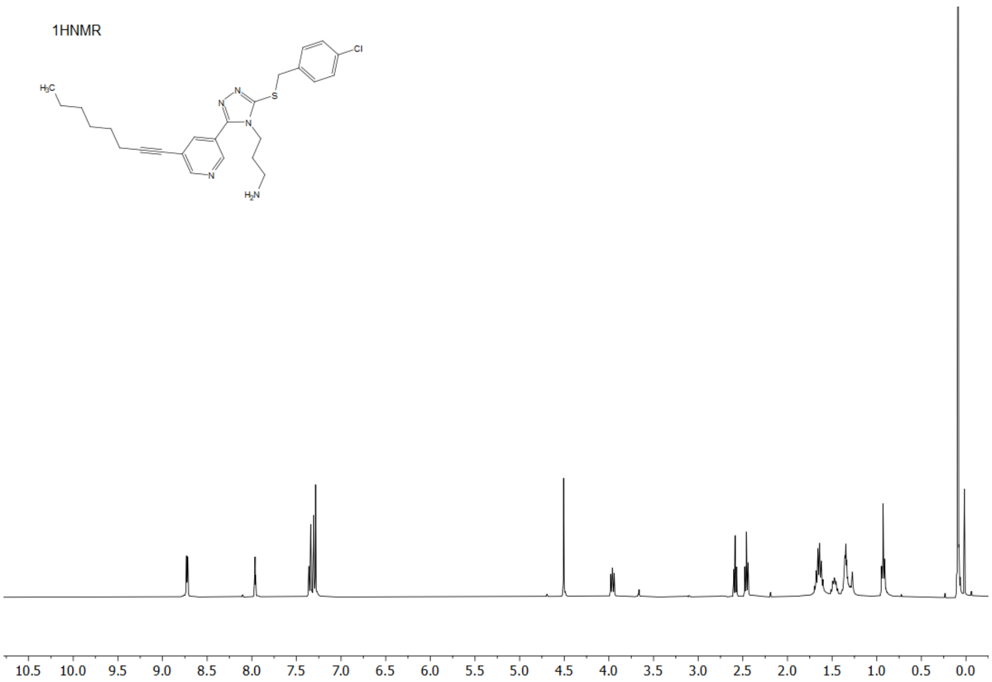

The $^{1}H$ NMR spectrum of compound **12b**

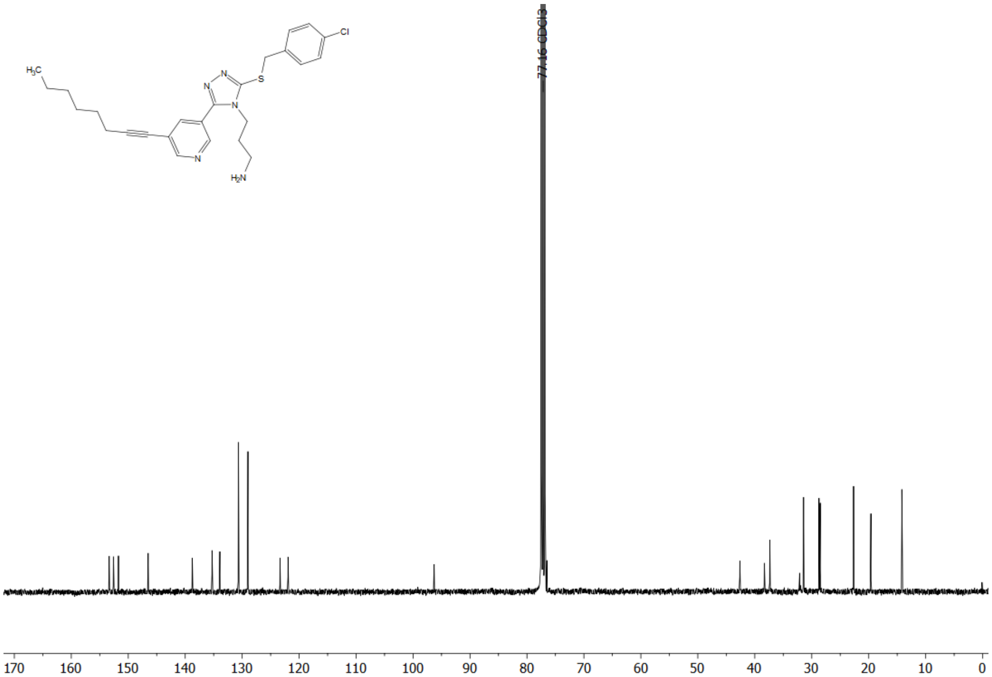

The $^{13}C$ NMR spectrum of compound **12b**

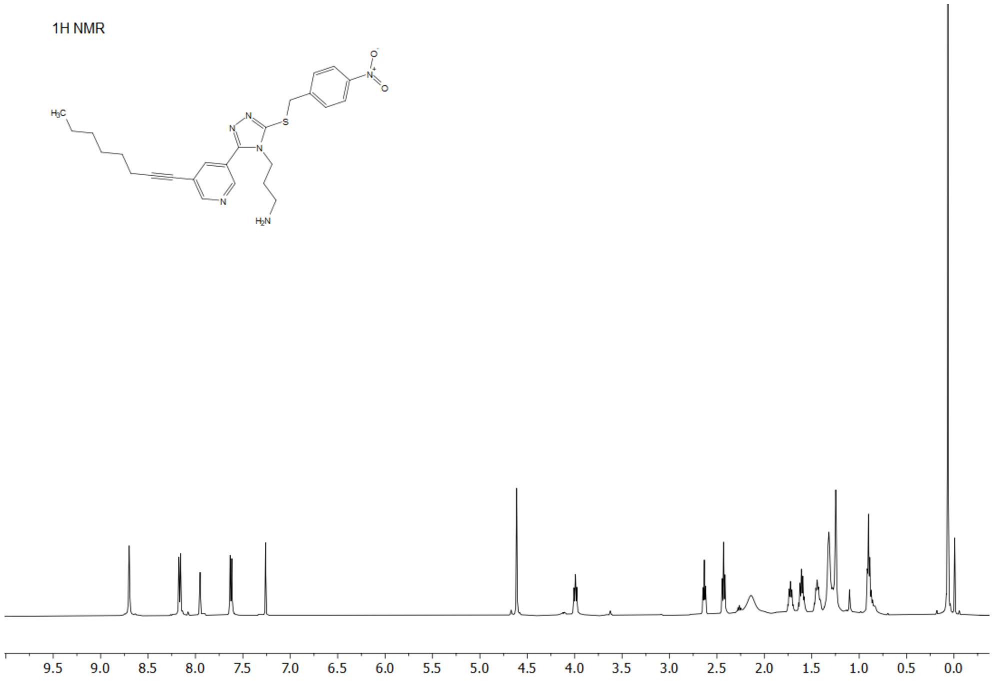

The $^{1}H$ NMR spectrum of compound **12c**

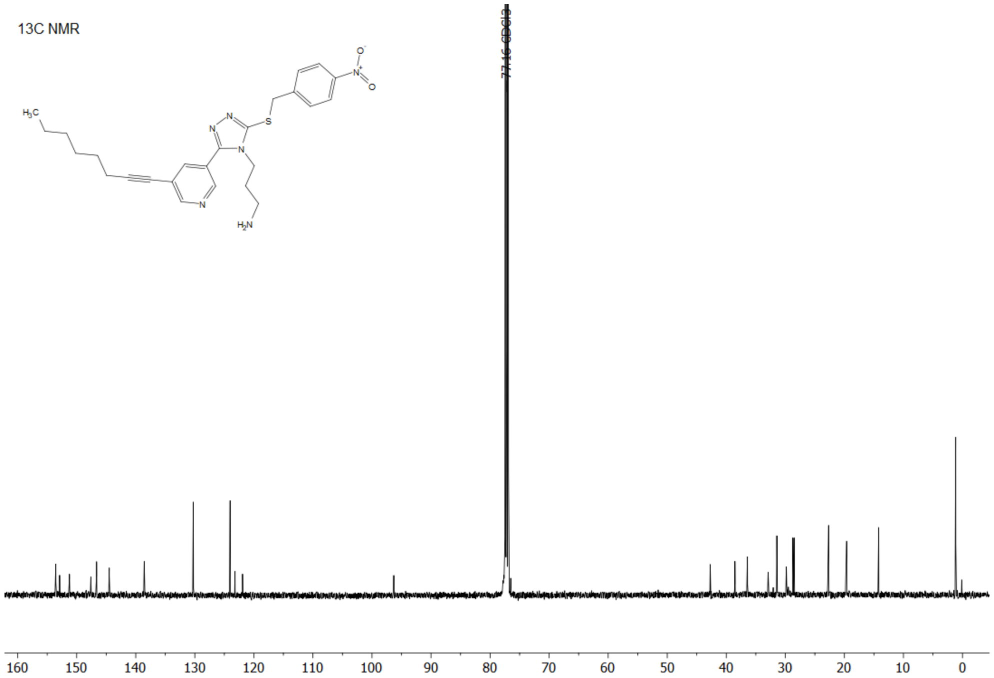

The $^{13}C$ NMR spectrum of compound **12c**

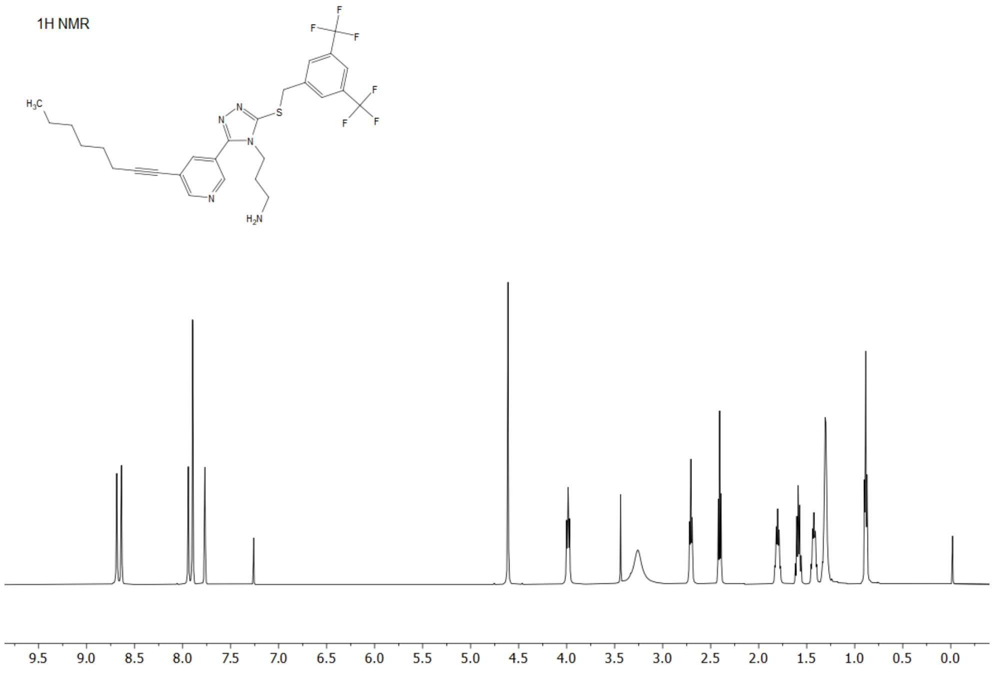

The $^{1}H$ NMR spectrum of compound **12d**

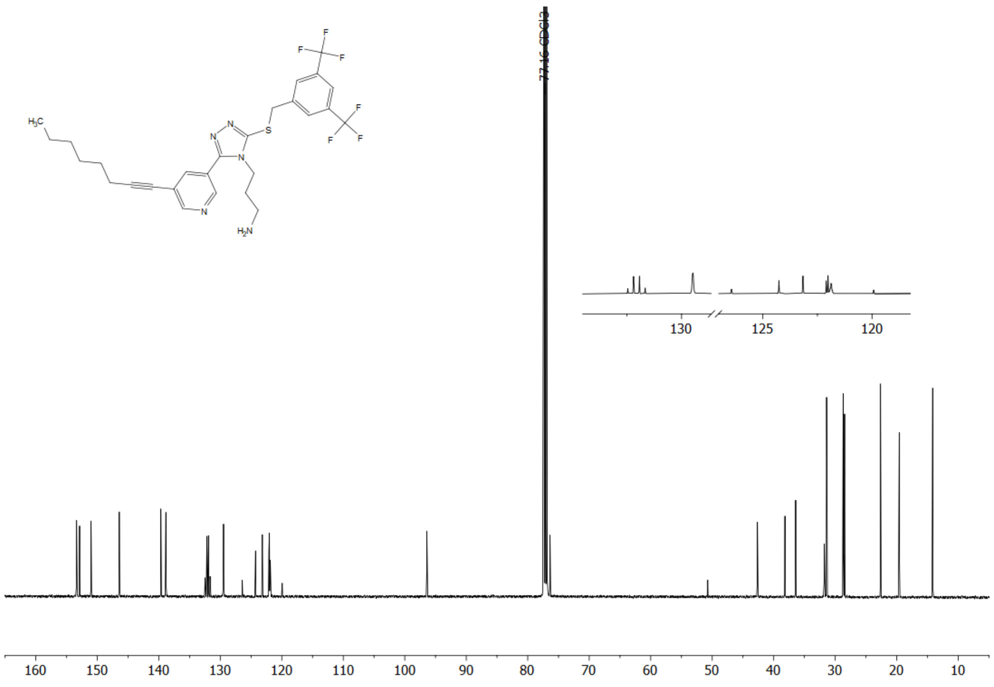

The $^{13}C$ NMR spectrum of compound **12d**

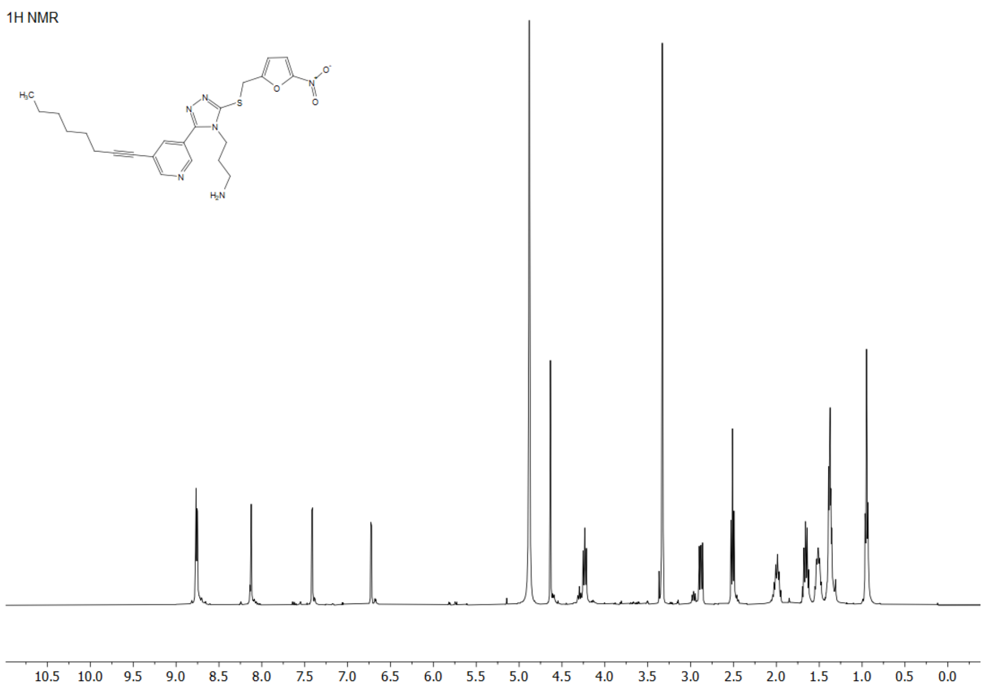

The $^{1}H$ NMR spectrum of compound **12e**

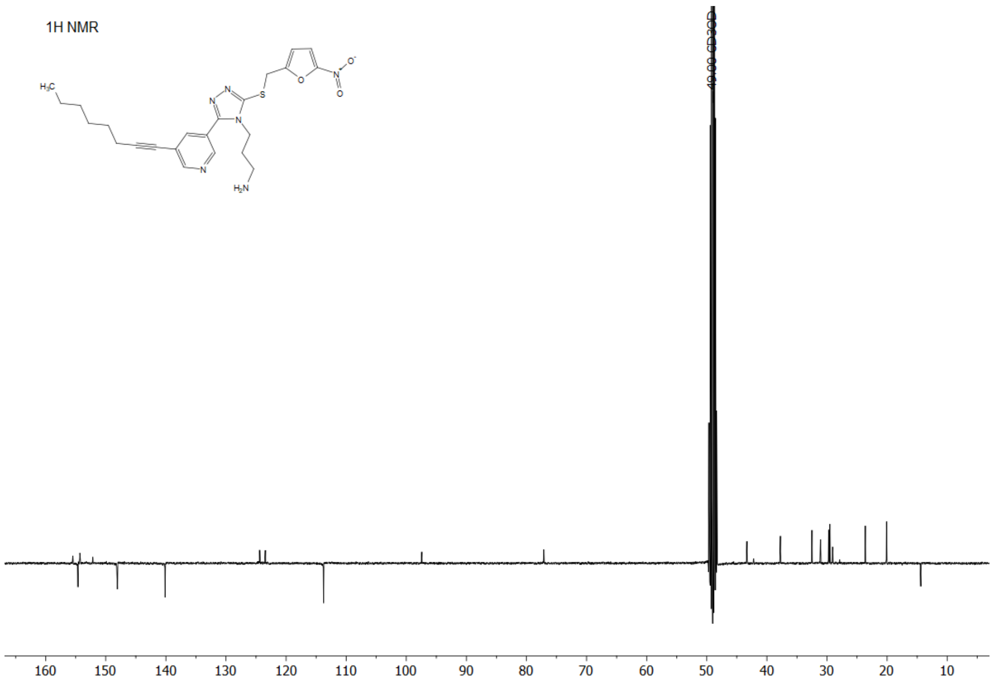

The $^{13}C$ NMR spectrum of compound **12e**

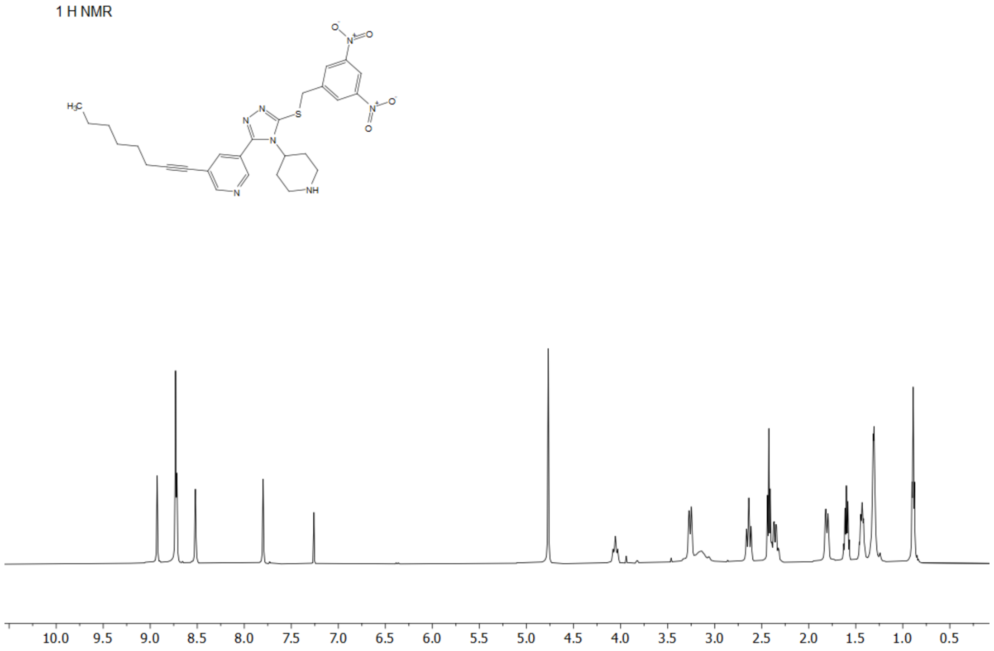

The $^{1}H$ NMR spectrum of compound **13**

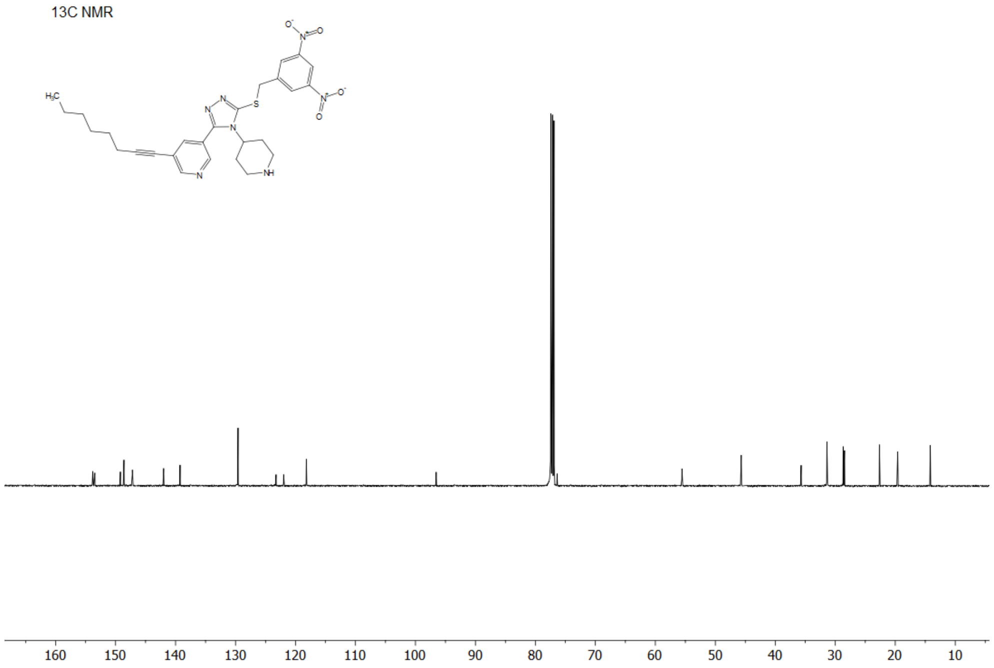

The $^{13}C$ NMR spectrum of compound **13**

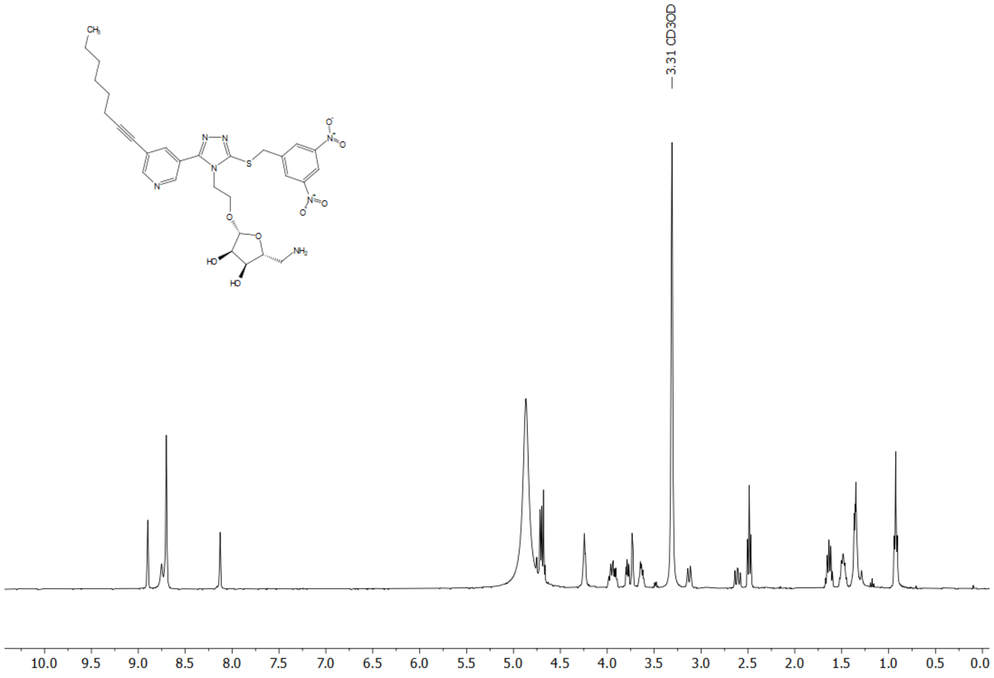

The $^{1}H$ NMR spectrum of compound **17**

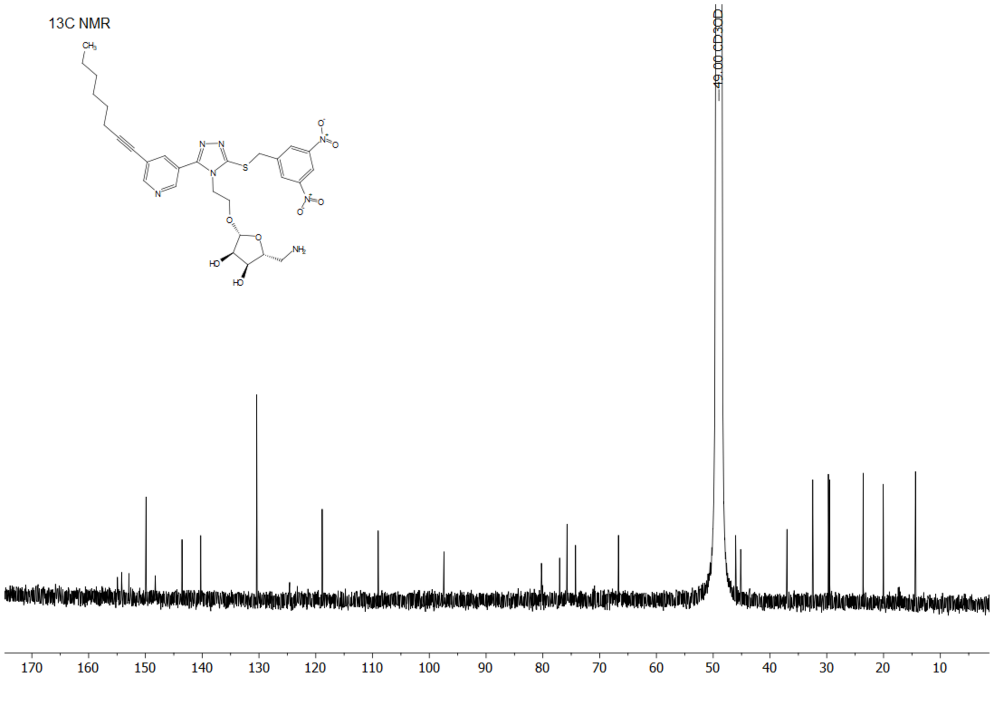

The $^{13}C$ NMR spectrum of compound **17**

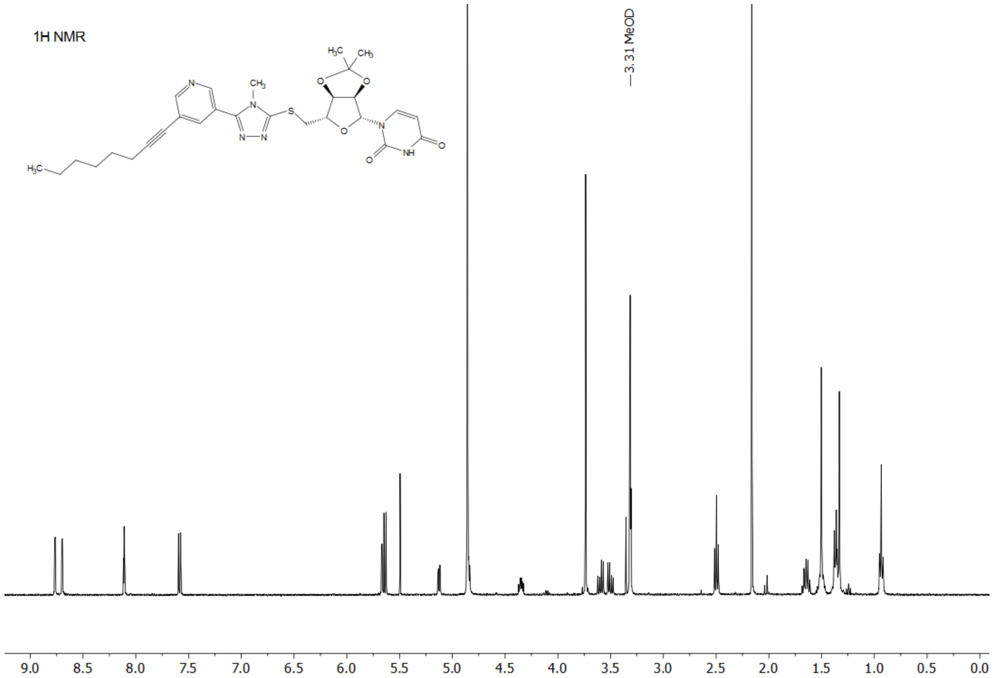

The $^{1}H$ NMR spectrum of compound **18**

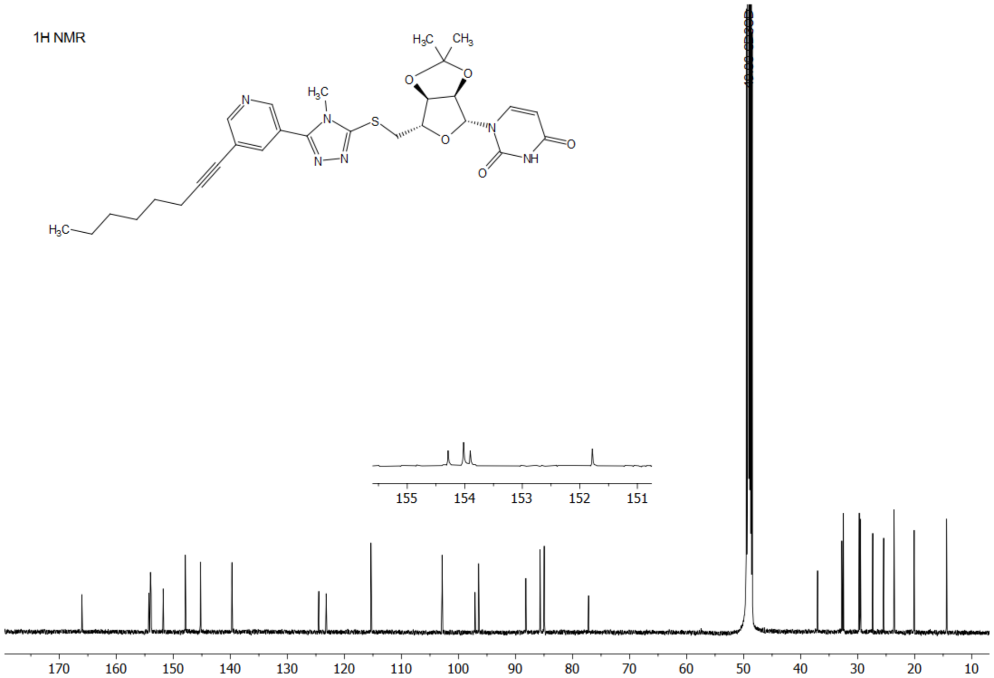

The $^{1}H$ NMR spectrum of compound **18**

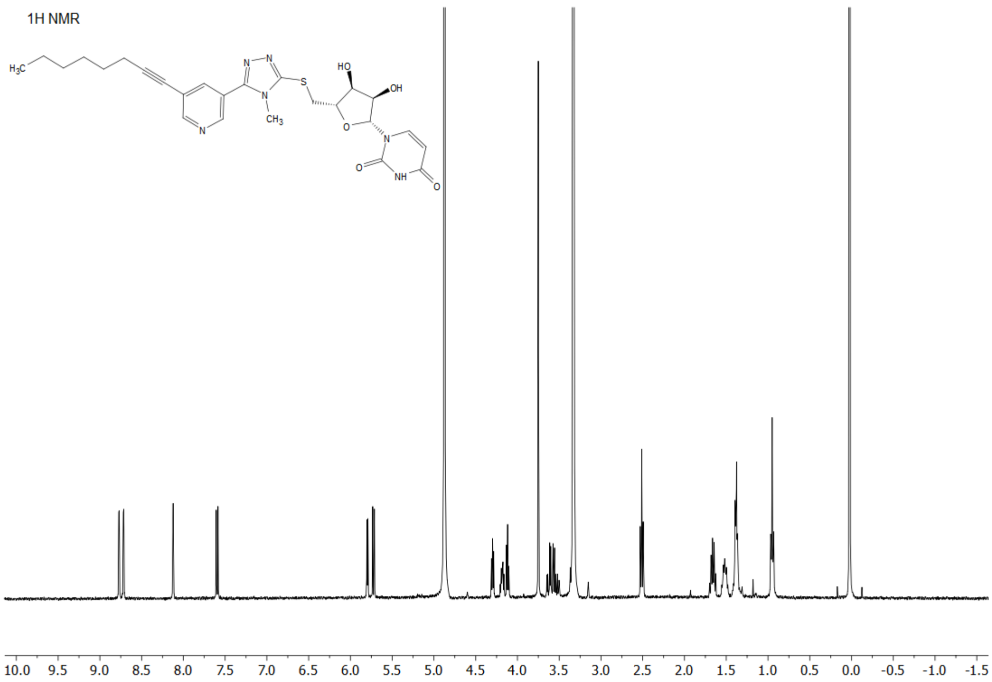

The $^1H$ NMR spectrum of compound **21**

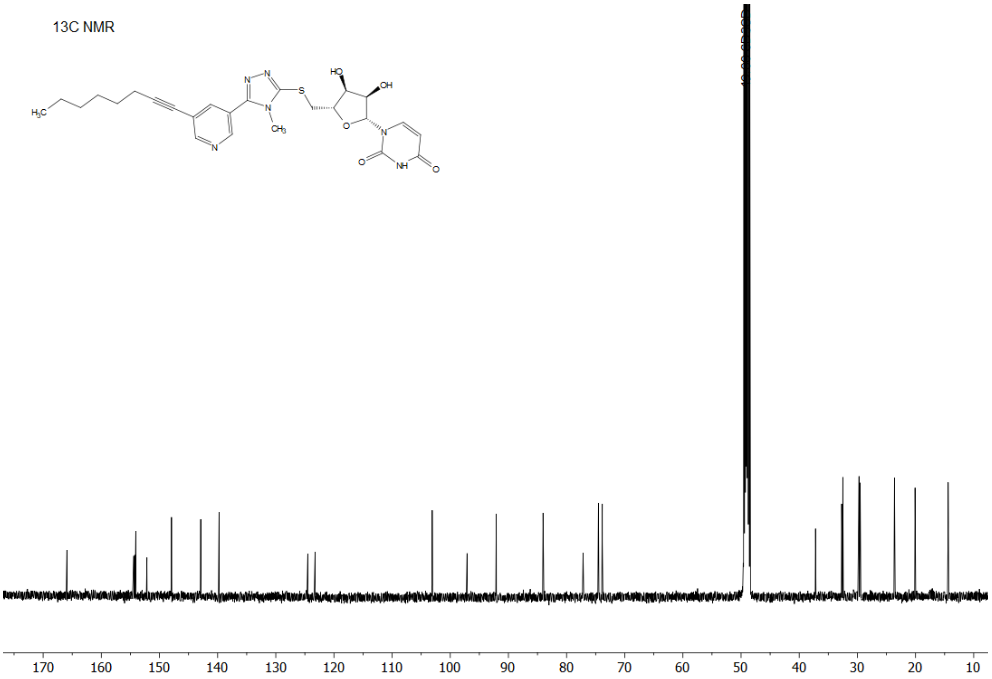

The $^{13}C$ NMR spectrum of compound **21**

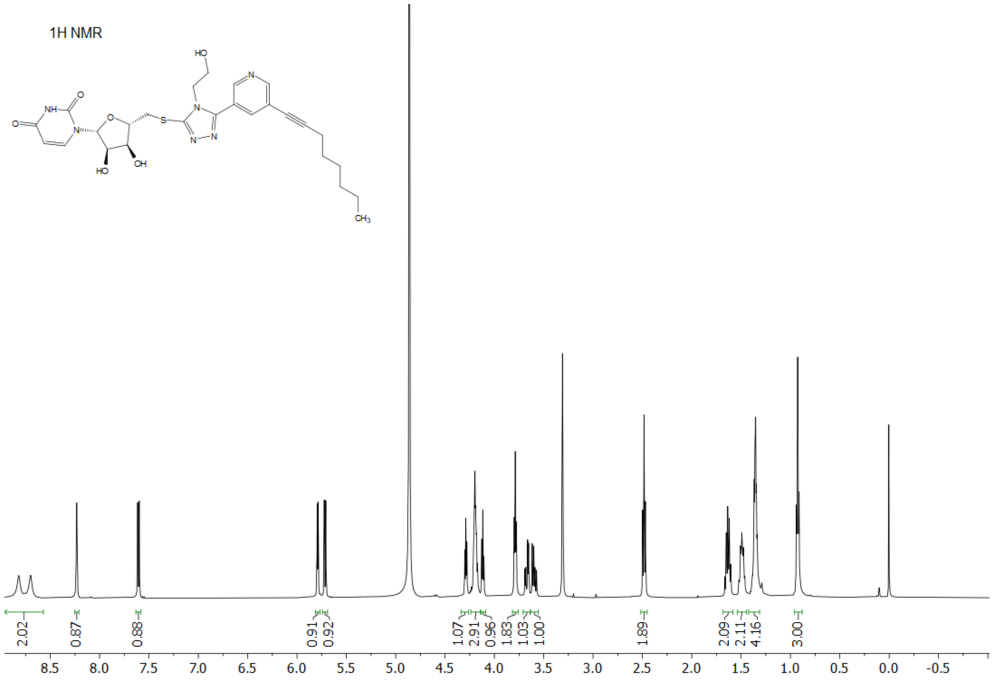

The $^{1}$H NMR spectrum of compound **22**

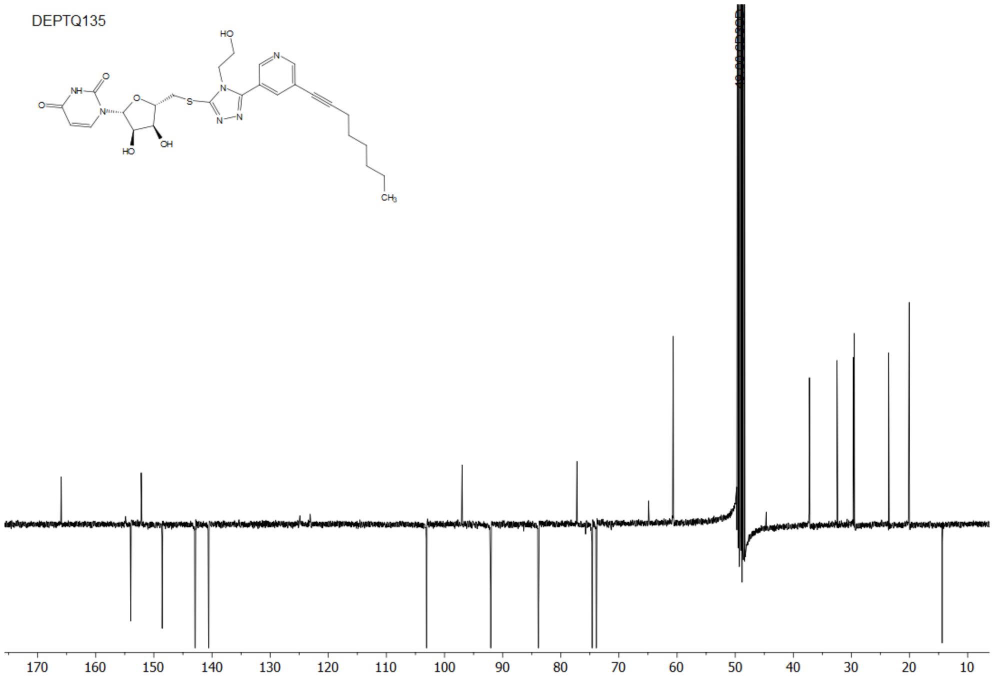

The $^{13}$C NMR spectrum of compound **22**

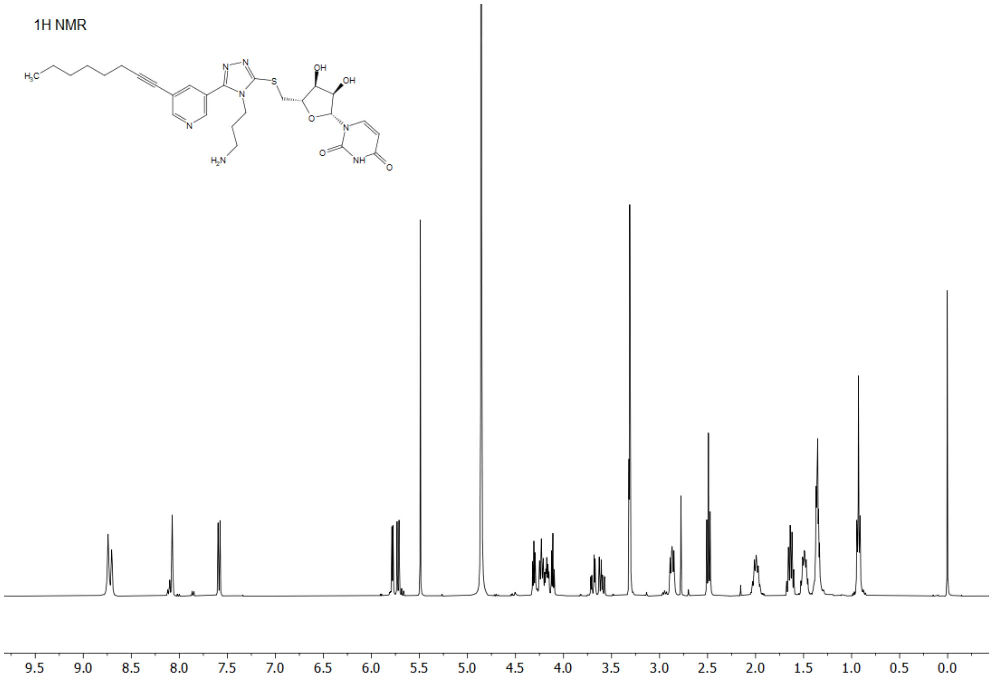

The $^{1}H$ NMR spectrum of compound **23**

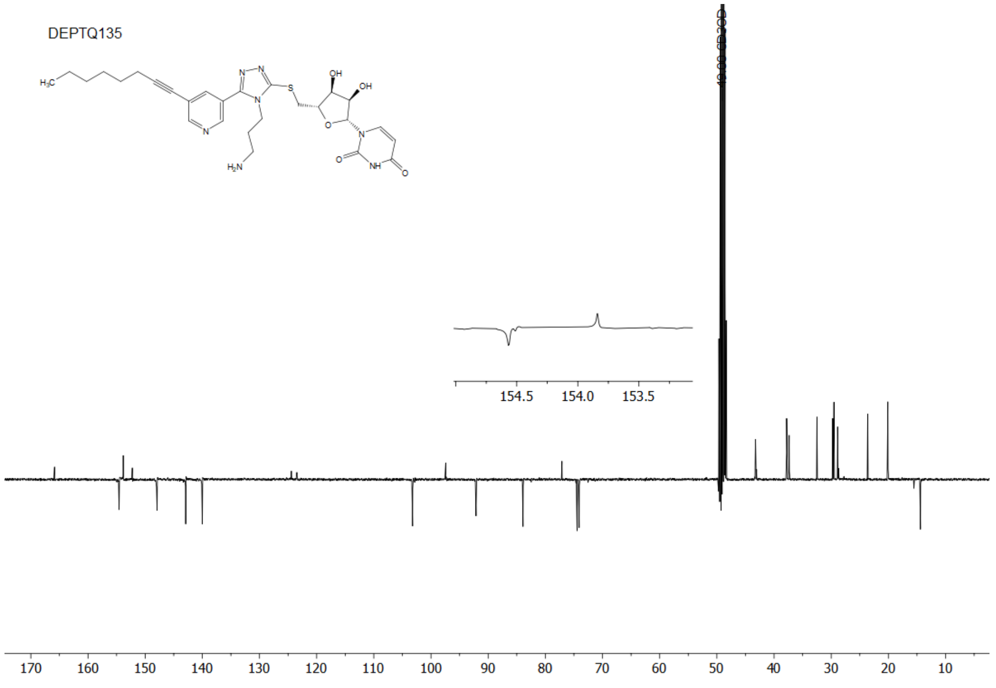

The $^{13}C$ NMR spectrum of compound **23**
